# Inferring Neural Activity Before Plasticity: A Foundation for Learning Beyond Backpropagation

**DOI:** 10.1101/2022.05.17.492325

**Authors:** Yuhang Song, Beren Millidge, Tommaso Salvatori, Thomas Lukasiewicz, Zhenghua Xu, Rafal Bogacz

## Abstract

For both humans and machines, the essence of learning is to pinpoint which components in its information processing pipeline are responsible for an error in its output — a challenge that is known as *credit assignment*. How the brain solves credit assignment is a key question in neuroscience, and also of significant importance for artificial intelligence. It has long been assumed that credit assignment is best solved by backpropagation, which is also the foundation of modern machine learning. However, it has been questioned whether it is possible for the brain to implement backpropagation and learning in the brain may actually be more efficient and effective than backpropagation. Here, we set out a fundamentally different principle on credit assignment, called *prospective configuration*. In prospective configuration, the network first infers the pattern of neural activity that should result from learning, and then the synaptic weights are modified to consolidate the change in neural activity. We demonstrate that this distinct mechanism, in contrast to backpropagation, (1) underlies learning in a well-established family of models of cortical circuits, (2) enables learning that is more efficient and effective in many contexts faced by biological organisms, and (3) reproduces surprising patterns of neural activity and behaviour observed in diverse human and animal learning experiments. Our findings establish a new foundation for learning beyond backpropagation, for both understanding biological learning and building artificial intelligence.

---

The credit assignment problem^1^ lies at the very heart of learning. *Backpropagation*^2–5^, as a simple yet effective credit assignment theory, has powered notable advances in artificial intelligence since its inception^6–11^. It has also gained a predominant place in understanding learning in the brain^1, 12–21^. Due to this success, much recent work has focused on understanding how biological neural networks could learn in a way similar to backpropagation^22–31^: although many proposed models do not implement backpropagation exactly, they nevertheless try to approximate backpropagation, and much emphasis is placed on how close this approximation is^22–28, 32–34^. However, learning in the brain is superior to backpropagation in many critical aspects — for example, compared to the brain, backpropagation requires many more exposures to a stimulus to learn^35^ and suffers from catastrophic interference of newly and previously stored information^36, 37^. This raises the question of whether using backpropagation to understand learning in the brain should be the main focus of the field.

Here, we propose that the brain instead solves credit assignment with a fundamentally different principle, which we call *prospective configuration*. In prospective configuration, before synaptic weights are modified, neural activity changes across the network so that output neurons better predict the target output; only then are the synaptic weights (weights, for short) modified to consolidate this change in neural activity. By contrast, in backpropagation the order is reversed — weight modification takes the lead and the change in neural activity is the result that follows.

We identify prospective configuration as a principle that is implicitly followed by a well-established family of neural models with solid biological groundings, namely, energy-based networks. They include Hopfield networks^38^ and predictive coding networks^39^, which have been successfully used to describe information processing in the cortex^40–46^. To support the theory of prospective configuration, we show that it can both yield efficient learning, which humans and animals are capable of, and reproduce data from experiments on human and animal learning. Thus, on the one hand, we demonstrate that prospective configuration performs more efficient and effective learning than backpropagation in various situations faced by biological systems, such as learning with deep structures, online learning, learning with a limited amount of training examples, learning in changing environments, continual learning with multiple tasks, and reinforcement learning. On the other hand, we demonstrate that patterns of neural activity and behaviour in diverse human and animal learning experiments, including sensorimotor learning, fear conditioning and reinforcement learning, can be naturally explained by prospective configuration, but not by backpropagation.

Guided by the belief that backpropagation is the foundation of biological learning, previous work showed that energy-based networks can closely approximate backpropagation. However, to achieve it, the networks were set up in an unnatural way, such that the neural activity was prevented from substantially changing before weight modification, by constraining the supervision signal to be infinitely small (e.g., as in equilibrium propagation^24^ and in previous studies employing predictive coding networks^25, 47^) or last an infinitely short time^33, 48^. In contrast, we reveal that the energy-based networks without these unrealistic constrains follow the distinct principle of prospective configuration rather than backpropagation, and are superior in both learning efficiency and accounting for data on biological learning.

Below, we first introduce prospective configuration with an intuitive example, show how it originates from energy-based networks, describe its advantages and quantify them in a rich set of biological-relevant learning tasks. Finally, we show that it naturally explains patterns of neural activity and behaviour in diverse learning experiments.

## Results

### Prospective configuration: an intuitive example

To optimally plan behaviour, it is critical for the brain to predict future stimuli — for example, to predict sensations in some modalities on the basis of other modalities^49^. If the observed outcome differs from the prediction, the weights in the whole network need to be updated so that prediction in the “output” neurons are corrected. Backpropagation computes how the weights should be modified to minimize the error on the output, and this weight update results in the change of neural activity when the network next makes the prediction. In contrast, we propose that the activity of neurons is first adjusted to a new configuration, so that the output neurons better predict the observed outcome (target pattern); the weights are then modified to reinforce this configuration of neural activity. We call this configuration of neural activity “prospective”, since it is the neural activity that the network *should produce* to correctly predict the observed outcome. In agreement with the proposed mechanism of prospective configuration, it has indeed been widely observed in biological neurons that presenting the outcome of a prediction triggers changes in neural activity — for example, in tasks requiring animals to predict a fruit juice delivery, the reward triggers rapid changes in activity not only in the gustatory cortex, but also in multiple cortical regions^50, 51^.

To highlight the difference between backpropagation and prospective configuration, consider a simple example in Fig. 1a. Imagine a bear seeing a river. In the bear’s mind, the sight generates predictions of hearing water and smelling salmon. On that day, the bear indeed smelled the salmon but did not hear the water, perhaps due to an ear injury, and thus the bear needs to change its expectation related to the sound. Backpropagation (Fig. 1b) would proceed by backpropagating the negative error, so as to reduce the weights on the path between the visual and auditory neurons. However, this also entails a reduction of the weights between visual and olfactory neurons that would compromise the expectation of smelling the salmon, the next time the river is visited; even though the smell of salmon was present and correctly predicted. These undesired and unrealistic side effects of learning with backpropagation are closely related with the phenomenon of catastrophic interference, where learning a new association destroys previously learned memories^36, 37^. This example shows that, with backpropagation, even learning one new aspect of an association may interfere with the memory of other aspects of the same association.

**Fig. 1.**
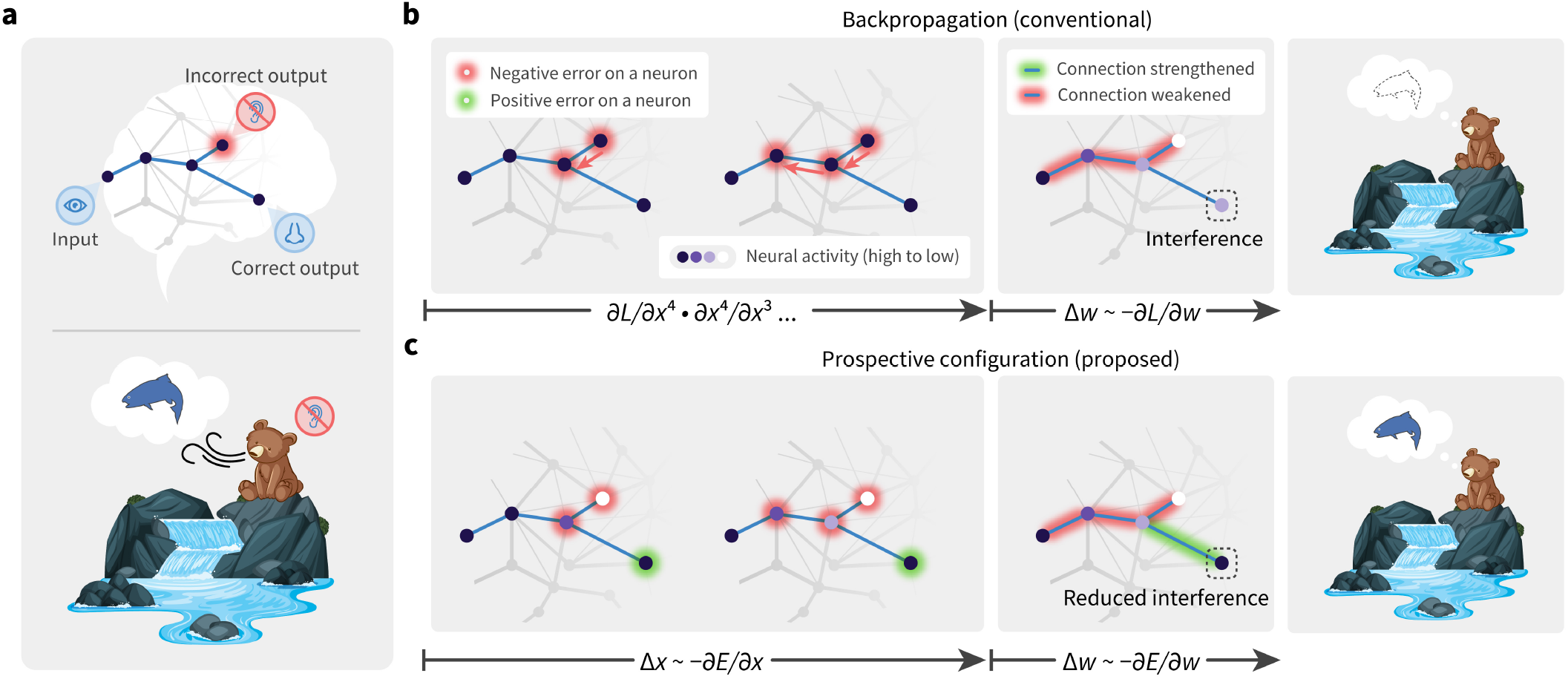
Prospective configuration avoids interference during learning. ▶ **a** | An abstract (top) and a concrete (bottom) example of a task inducing interference during learning. One stimulus input (seeing the water) triggers two prediction outputs (hearing the water and smelling the salmon). One output is correct (smelling the salmon), while the other one is an error (not hearing the water). Backpropagation produces interference during learning: not hearing the water reduces the expectation of smelling the salmon (panel b), although the salmon was indeed smelled. Prospective configuration, on the other hand, avoids such interference (panel c). ▶ **b** | In backpropagation, negative error propagates from the error output into hidden neurons (left). This causes a weakening of some connections, which on the next trial improves the incorrect output, but it also reduces the prediction of the correct output, thus introducing interference (middle and right). ▶ **c** | In prospective configuration, neural activity settles into a new configuration (purple of different intensity) before weight modification (left). This configuration corresponds to the activity that should be produced after learning, i.e., is “prospective”. Hence it foresees the positive error on the correct output, and modifies the connections to improve the incorrect output, while maintaining the correct output (middle and right).

In contrast, prospective configuration assumes that learning starts with the neurons being configured to a new state — which corresponds to a pattern enabling the network to correctly predict the observed outcome. The weights are then modified to consolidate this state. This behaviour can “foresee” side effects of potential weight modifications and compensate for them dynamically — Fig. 1c: to correct the negative error on the incorrect output, the hidden neurons settle to their prospective state of lower activity, and as a result, a positive error is revealed and allocated to the correct output. Consequently, prospective configuration increases the weights connecting to the correct output, while backpropagation does not (cf. middle plots of Fig. 1b and c). Hence, prospective configuration is able to correct the side effects of learning an association effectively, efficiently, and with little interference.

### Origin of prospective configuration: energy-based networks

To shows how prospective configuration naturally arises in energy-based networks, we introduce a physical machine analog, that provides an intuitive understanding of energy-based networks, and how they produce the mechanism of prospective configuration.

Energy-based networks have been widely and successfully used in describing biological neural systems^38, 39, 53–55^. In these models, a neural circuit is described by a dynamical system driven by reducing an abstract “energy”, e.g., reflecting errors made by the neurons; see Methods. Neural activity and synaptic weights change to reduce this energy, hence they can be considered as “movable parts” of the dynamical system. We show below that energy-based networks are mathematically equivalent to a physical machine (we call it *energy machine*), where the energy function has an intuitive interpretation and its dynamics are straightforward — the energy machine simply adjusts its movable parts to reduce energy.

As shown in Fig. 2a–b, the energy machine includes nodes sliding on vertical posts, connected with each other via rods and springs. Translating from energy-based networks to the energy machine, the neural activity maps to the vertical position of a solid node, a connection maps to a rod (blue arrow) pointing from one node to another (where the weight determines how the end position of the rod relates to the initial position), and the energy function maps to the elastic potential energy of springs with nodes attached on their both ends (the natural length of the springs is zero). Different energy functions and networks structures result in different energy-based networks, corresponding to energy machines with different configurations and combinations of nodes, rods, and springs. In Fig. 2, we present the energy machine of predictive coding networks^25, 40, 52^, because they are most accessible and established to be closely related to backpropagation^25, 33^.

**Fig. 2.**
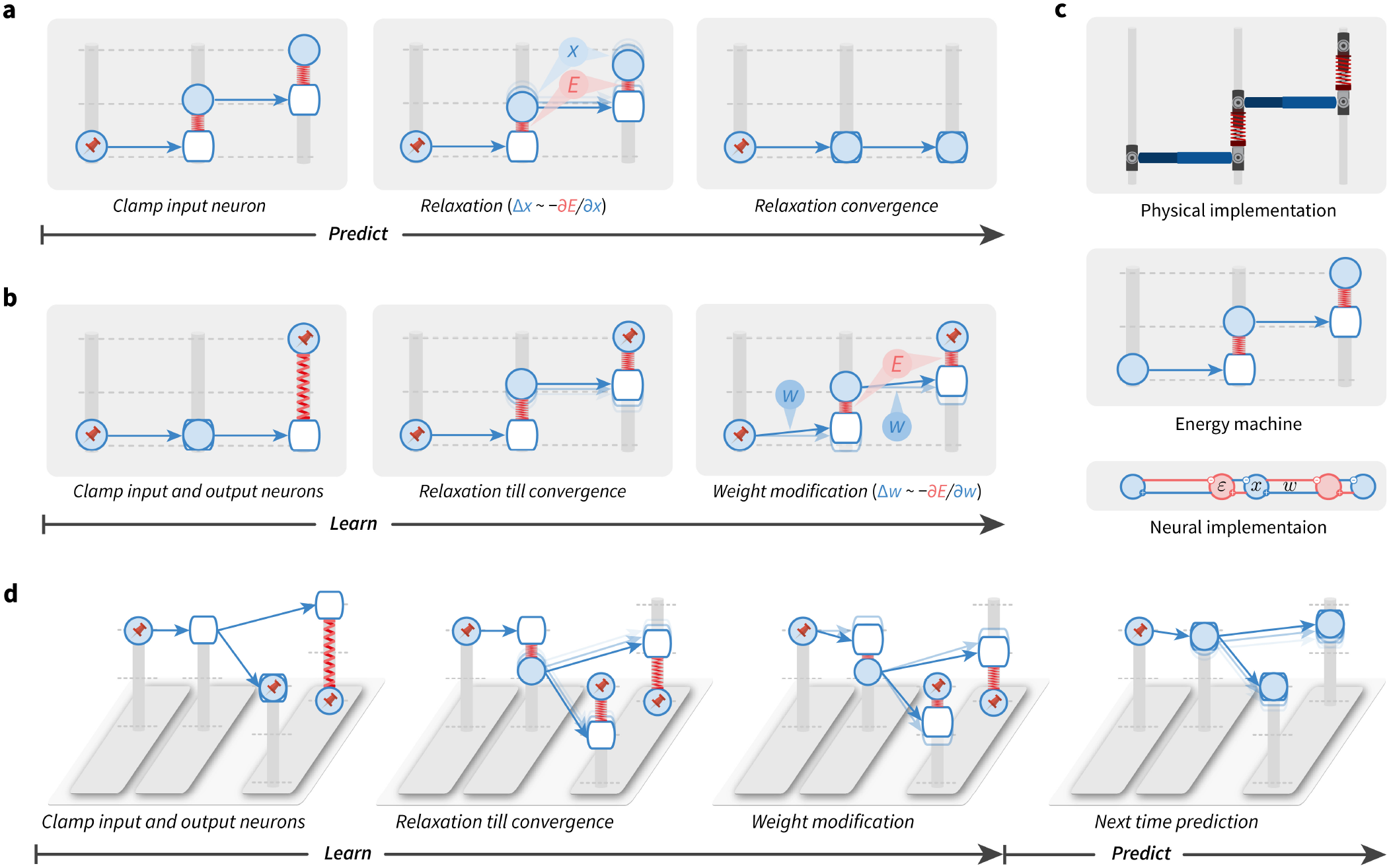
The energy machine reveals a new understanding of energy-based networks, the mechanism of prospective configuration, and its theoretical advantages. A subset of energy-based networks can be visualized as mechanical machines that perform equivalent computations. Here, we present one of them, predictive coding networks^25,40,52^. In the energy machine, the activity of a neuron corresponds to a height of a node (represented by a solid circle) sliding on a post. The input to the neuron is represented by a hollow node on the same post. A synaptic connection corresponds to a rod pointing from a solid to a hollow node. The synaptic weight determines how the input to a post-synaptic neuron depends on the activity of pre-synaptic neuron, hence it influences the angle of the rod. In energy-based networks, relaxation (i.e., neural dynamics) and weight modification (i.e., weight dynamics) are both driven by minimizing the energy, thus correspond to relaxing the energy machine by moving the nodes and tuning the rods, respectively. ▶ **a-b** | Predictions (a) and learning (b) in energy-based networks, visualized by the energy machine. The pin indicates that the neural activity is fixed to the input or target pattern. Here, it is revealed that the relaxation infers the prospective neural activity, towards which the weights are then modified, a mechanism that we call prospective configuration. ▶ **c** | The physical implementation (top) and the connectivity of a predictive coding network^25,40,52^ (bottom), which has a dynamics mathematically equivalent to the energy machine in the middle (see Methods for details). ▶ **d** | The learning problem in Fig. 1, visualized by the energy machine, which learns to improve the incorrect output while not interfering with the correct output, thanks to the mechanism of prospective configuration.

The dynamics of energy-based networks, which are driven by minimizing the energy function, maps to the relaxation of the energy machine, which is driven by reducing the total elastic potential energy on the springs. A prediction with energy-based networks involves clamping the input neurons to the provided stimulus and updating the activity of the other neurons, which corresponds to fixing one side of the energy machine and letting the energy machine relax by moving nodes (Fig. 2a). Learning with energy-based networks involves clamping the input and output neurons to the corresponding stimulus, first letting the activity of the remaining neurons converge and then updating weights, which corresponds to fixing both sides of the energy machine and letting the energy machine relax first by moving nodes and then by tuning rods (Fig. 2b).

The energy machine reveals the essence of energy-based networks: the relaxation before weight modification lets the network settle to a new configuration of neural activity, corresponding to those that would have occurred after the error was corrected by the modification of weights, i.e., prospective activity (thus, we call this mechanism prospective configuration). For example, the second layer “neuron” in Fig. 2b increases its activity, and this increase in activity would also be caused by the subsequent weight modification (of the connection between the first and the second neurons). In simple terms, the relaxation in energy-based networks infers the prospective neural activity after learning, towards which the weights are then modified. This distinguishes it from backpropagation, where the weights modification takes the lead, and the change in neural activity is the result that follows.

The bottom part of Fig. 2c shows the connectivity of a predictive coding network^25, 40, 52^, which has a dynamics mathematically equivalent to the energy machine shown above it. Predictive coding networks include neurons (blue) corresponding to nodes on the posts, and separate neurons encoding prediction errors (red) corresponding to springs. For details, see Methods and Extended Data Fig. 1, where we list equations describing predictive coding networks, show how they map on the neural implementation and the proposed energy machine.

Using the energy machine, Fig. 2d simulates the learning problem from Fig. 1. Here, we can see that prospective configuration indeed foresees the result of learning and its side effects, through relaxation. Hence, it learns to avoid interference within one iteration, which would otherwise take multiple iterations for backpropagation.

### Advantages of prospective configuration: reduced interference and faster learning

Here we quantify interference in the above scenario and demonstrate how the reduced interference translates into an advantage in performance. In all simulations in the main text prospective configuration is implemented in predictive coding networks (see Methods, other energy-based models are considered in Extended Data Figures and Supplementary Information). Fig. 3a compares the activity of output neurons in the example in Fig. 1, between backpropation and prospective configuration. Initially both output neurons are active (top right corner), and the output should change towards a target in which one of the neurons is inactive (red vector). Learning with prospective configuration results in changes on the output (purple solid vector) that are aligned better with the target than those for backpropagation (purple dotted vector). Following the first update of weights, we simulate multiple iterations until the network is able to correctly predict the target. Here, “iteration” refers to each time the agent is presented with stimuli and conducts one weight update because of the stimulus (a trial-by-trial iteration). Within each iteration, it contains: (1) numerical integration procedure of relaxation of energy-based networks, which captures its continuous process; (2) one update of weights at the end of the above procedure. Although the output from backpropagation can reach the target after multiple iterations, the output for the “correct neuron” diverges from the target during learning and then comes back - it is particularly undesired effect in biological learning, where networks can be “tested” at any point during the learning process, because it may lead to incorrect decisions affecting chances for survival. By contrast, prospective configuration substantially reduces this effect.

**Fig. 3.**
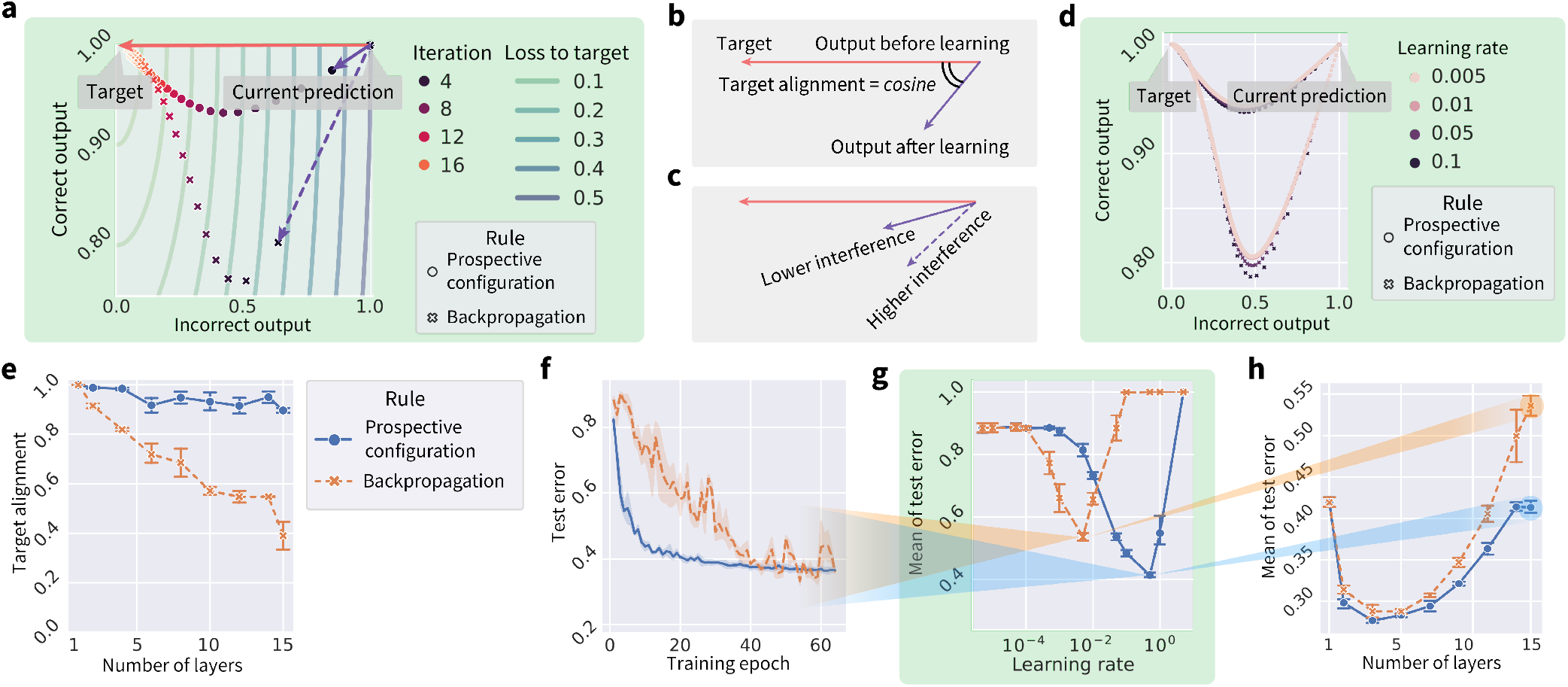
Learning with prospective configuration changes the activity of output neurons in a direction more aligned towards the target. ▶ **a** | Simulation of network from Fig. 1 showing changes of the correct and incorrect output neurons during training (“Iteration”), trained with both learning rules. Here, learning with prospective configuration (purple solid vector) aligns better with the target (red vector), than for backpropagation (purple dashed vector). ▶ **b** | The interference can be quantified by “target alignment”: the cosine similarity of the direction of target (red vector) and the direction of learning (purple vector). ▶ **c** | Higher target alignment indicates less interference and vice versa. ▶ **d** | The same experiment as in panel a repeated with a learning rate ranging from to 0.5 represented by the size of the markers, where it is shown that the choice of learning rate slightly changes the trajectories for both methods but the conclusion holds irrespective of the learning rate. ▶ **e** | Target alignment of randomly generated networks trained with both learning rules, as a function of depth of the network. Here, target alignment drops as the network gets deeper, demonstrating the difficulty of training deep structures. However, prospective configuration maintains much higher target alignment along the way. ▶ **f** | Classification error during training on FashionMNIST^56^ dataset containing images of clothing belonging to different categories, for both learning rules, with a deep neural network of 15 layers. ▶ **g** | Mean of the classification error over training epochs (reflecting how fast test error drops), as a function of learning rate. Results in the panels f and h are for the learning rates giving the minima of the corresponding curves in this panel. ▶ **h** | Mean of classification error of other network depths. Each point is from learning rate independently optimized for each learning rule in the corresponding setup of network depth. In panels e–h, prospective configuration demonstrates notable advantage as the structure gets deep.

Although backpropagation modifies the weights to directly reduce the cost in the space of weights (i.e., performs gradient descent), surprisingly and rather subversively, it does not push the resulting output activity directly towards the target. To illustrate this, Fig. 3a visualizes the cost with contour lines. Changing the activity of output neurons according to the gradient of the cost would correspond to a change orthogonal to the contour lines, i.e., that indicated by the red arrow. However, backpropagation changes the output in a different direction shown by a dashed arrow. Since the network is a complex cascaded system, optimizing the weights independently, without considering the effect of update of other weights, leads to the output activity not updating towards the target directly, due to different weight updates to different layers interfering with each other. By contrast, when updating each weight, prospective configuration considers the results of update of other weights by finding a desired configuration of neural activity first, and such mechanism is missing in backpropogation but natural in energy-based networks. Extended Data Fig. 2 shows a direct comparison of how these two models evolve in weight and output spaces during learning.

The interference can be quantified by the angle between the direction of target (from current output to target) and learning (from current output to output after learning, both measured without target provided), and we define “target alignment” as the cosine of this angle (Fig. 3b), hence high interference corresponds to low target alignment (Fig. 3c). It is useful to highlight that the target alignment is little affected by the learning rate, as shown by Fig. 3d, demonstrating that the learning rate has little effect on the direction and trajectory output neurons take. The difference in target alignment demonstrated in Fig. 3a is also present for deeper and larger (randomly generated) networks, as shown in Fig. 3e. When a network has no hidden layers, the target alignment is equal to 1 (proved in section 2.4.1 of Supplementary Information). The target alignment drops for backpropagation as the network gets deep, because changes in weights in one layer interfere with changes in other layers (as explained in Fig. 1) and the backpropagated errors do not lead to appropriate modification of weights in hidden layers (Extended Data Fig. 2). By contrast, prospective configuration maintains a much higher value along the way. This higher target alignment of prospective configuration can be theoretically explained by the following: (i) there exists a close link between prospective configuration and an algorithm called target propagation^57^ (shown in Extended Data Fig. 3 and section 2.2 of Supplementary Information); and (ii) under certain conditions target propagation^57^ has target alignment of 1^58^ (demonstrated in Extended Data Fig. 4 and Section 2.4.2 of Supplementary Information). Thus, the link with target propagation^57^ provides a theoretical insight (with numerical verification) on why prospective configuration has a higher target alignment.

The effectiveness of target alignment directly translates to the efficiency of learning: Fig. 3f shows that the test error during training in a visual classification task with a deep neural network of 15 layers decreases faster for prospective configuration than backpropagation. (“test error” refers to the ratio of incorrectly classified samples in all samples on the test set).

Throughout the whole paper, if learning rate is not presented in a plot, the plot corresponds to the best learning rate optimized independently for each rule under the setup, via a grid search. The optimization target is either the learning performance or approximation to experimental recordings, depending on the nature of the experiment (details can be found in method section for each experiment). Thus for example, Fig. 3f shows the test errors as training progress, and they are with the learning rates optimized independently for each learning rule. The optimization target is the “mean of test error” during training (reflecting how fast the test error decreases during training). Fig. 3g plots this “mean of test error” for different learning rates for both learning rules, and the learning rates giving the minima of the curves have been used in Fig. 3f.

Fig. 3h repeats the experiment on networks of other depths, and shows the mean of the test error during training (reflecting how fast the test error drops), as a function of network depth. The mean error is higher for low depths, as these networks are unable to learn the task, and for greater depths, because it takes longer to train deeper networks. Importantly, the gap between backpropagation and prospective configuration widens for deeper networks, paralleling the difference in target alignment. Efficient training with deeper networks is important for biological neural systems, known to be deep, e.g., primate visual cortex^59^.

In the Supplementary Information we develop a formal theory of prospective configuration and provide further illustrations and analyses of its advantages. Extended Data Figs. 5 formally defines prospective configuration and demonstrates that it is indeed commonly observed in different energy-based networks. Extended Data Figs. 6 and 7 empirically verify and generalize the advantages expected from the theory: they show that prospective configuration yields more accurate error allocation and less erratic weight modification, respectively.

### Advantages of prospective configuration: effective learning in biologically relevant scenarios

Inspired by these advantages, we show empirically that prospective configuration indeed handles various learning problems that biological systems would face better than backpropagation. Since the field of machine learning has developed effective benchmarks for testing learning performance, we use variants of classic machine learning problems that share key features with the learning in natural environments. Such problems include online learning where the weights must be updated after each experience (rather than a batch of training examples)^60^, continual learning with multiple tasks^61, 62^, learning in changing environments^63^, learning with limited amount of training examples, and reinforcement learning^10^. In all the aforementioned learning problems, prospective configuration demonstrates a notable superiority over backpropagation.

Firstly, based on the example in Fig. 1, we expect prospective configuration to require fewer episodes for learning than backpropagation. Before presenting the comparison, we describe how backpropagation is used to train artificial neural networks. Typically, the weights are only modified after a batch of training examples, based on the average of updates derived from individual examples (Fig. 4a). In fact, back-propagation relies heavily on averaging over multiple experiences to reach human-level performance^66–68^ as it needs to stabilise training^69^. By contrast, biological systems must update the weights after each experience, and we compare the learning performance in such a setting. The sampling efficiency can be quantified by mean of test error during training, which is shown in Fig. 4b as a function of batch size (number of experiences that the updates are averaged over). The efficiency strongly depends on batch size for backpropagation, because it requires batch-training to average out erratic weight updates, while this dependence is weaker for prospective configuration, where the weight changes are intrinsically less erratic and the batch-averaging is less required (see Extended Data Figs. 7). Importantly, prospective configuration learns faster with smaller batch sizes, as in biological settings. Additionally, the final performance can be quantified by the minimum of the test error, which is shown in Fig. 4c, when trained with batch size equal to one. Here, prospective configuration also demonstrates a notable advantage over backpropagation.

**Fig. 4.**
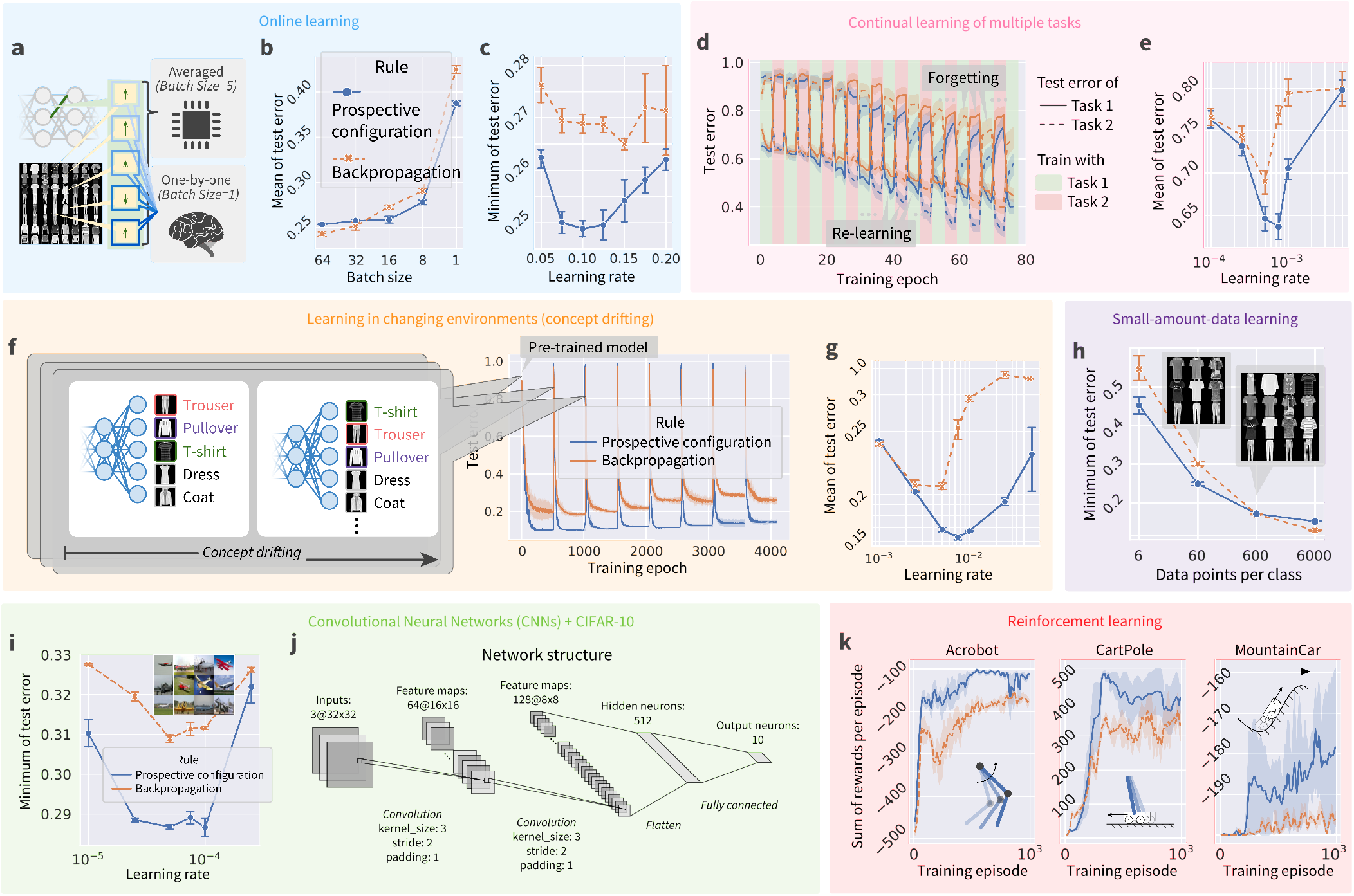
Prospective configuration achieves a superior performance over backpropagation in various learning situations faced by biological systems. These situations are: online learning^60^ (a–c), continual learning of multiple tasks^61,62^ (d–e), learning in changing environments^63^ (f–g), learning with a limited amount of training examples (h), and reinforcement learning^10^ (k). Panels corresponding to each situation are grouped together with the same background colour. Simulations of each situation differ from the “default setup” described in the Methods in a single aspect unique to this task. For example, the default setup involves training with mini-batches, so the batch size was only set to 1 in (a–c) for investigating online learning, while it was set to a larger default value in rest of the groups (panels). In supervised learning setups, fully-connected networks (a–h) are evaluated on FashionMNIST^56^ dataset and convolutional neural networks^64^ (i–j) are evaluated on CIFAR-10^65^ dataset. In reinforcement learning setup (k), fully-connected networks are evaluated on three classic control problems. If the learning rate is not presented in a plot, each point (a setup of experiment) in the plot corresponds to the best learning rate optimized independently for the each rule under that setup. ▶ **a** | Difference in training setup between computers that can average weight modifications for individual examples to get a “statistically good” value, and biological systems which must apply one modification before computing another. ▶ **b** | Mean of the test errors during training, as a function of batch size. ▶ **c** | Minimum of the test error during training as a function of learning rate. ▶ **d** | Test error during continual learning of two tasks. ▶ **g** | Mean of test error of both tasks during training as a function of learning rate. ▶ **f** | Test error during training when learning with concept drifting. ▶ **g** | Mean of test error during training with concept drifting as a function of learning rate. ▶ **h** | Minimum of the test errors during training, with different amounts of training examples (datapoints per class). ▶ **i** | Minimal of test error during training of a convolutional neural network trained with with prospective configuration and backpropagation on CIFAR-10^65^ dataset. ▶ **j** | The structure detail of the convolutional neural network used in the last panel. ▶ **k** | Sum of rewards per episode during training on three classic reinforcement learning tasks (insets). An episode is a period from initialization of environment to reaching a terminate state.

Secondly, biological organisms need to sequentially learn multiple tasks, while artificial neural networks show catastrophic forgetting: when trained on a new task, performance on previously learnt tasks is largely destroyed^36, 70–72^. Fig. 4d shows the performance when trained on two tasks alternately (task 1 is classifying five randomly selected classes in FashionMNIST dataset, and task 2 is classifying the remaining five classes). It shows that prospective configuration outperforms backpropagation in both terms of avoiding forgetting previous tasks and re-learning current tasks. Fig. 4e summarizes the results.

Thirdly, biological systems often need to rapidly adapt to changing environments. A common way to simulate this is “concept drifting”^63^, where a part of the mapping between the output neurons to the semantic meaning is shuffled every period of time (Fig. 4f left). Fig. 4f right shows the test error during training with concept drifting. Before epoch 0, both learning rules are initialized with the same pre-trained model (trained with backpropagation), thus, the epoch 0 is the first time the model experiences concept drift. Fig. 4g summarizes the results, and shows that for this task there is a particularly large difference in mean error (for optimal learning rates). This large advantage of prospective configuration is related to it being able to optimally detect which weights to modify (see Extended Data Figs. 6), and to preserve existing knowledge while adapting to changes (Fig. 1). This ability to maintain important information while updating other is critical for survival of animals in natural environments that are bound to change, and prospective configuration has a very substantial advantage in this respect.

Furthermore, biological learning is also characterized by a limited data availability. Fig. 4h show that prospective configuration outperforms backpropagation when the model is trained with fewer examples. To demonstrate the advantage of prospective configuration also scales up to larger networks and problems, we evaluate convolutional neural networks^64^ on CIFAR-10^65^ trained with both learning rules (Fig. 4i), where prospective configuration shows notable advantages over backpropagation. The detailed structure of the convolutional networks are given in Fig. 4j.

Another key challenge for biological systems is to decide which actions to take. Reinforcement learning theories (e.g., *Q*-learning) propose that it is solved by learning the expected reward resulting from different actions in different situations^73^. Such prediction of rewards can be made by neural networks^10^, which can be trained with prospective configuration or backpropagation. The sum of rewards per episode during training on three classic reinforcement learning tasks is reported in Fig. 4k, where prospective configuration demonstrates a notable advance over backpropagation. This large advantage may arise because reinforcement learning is particularly sensitive to erratic changes in network’s weights (as the target output depends on reward predicted by the network itself for a new state - see Methods).

Based on the superior learning performance of prospective configuration, we may expect that this learning mechanism has been favored by evolution, thus in the next sections we investigate if it can account for neural activity and behaviour during learning better than backpropagation.

### Evidence for prospective configuration: inferring of latent state during learning

Prospective configuration is related to theories proposing that before learning, the brain first infers a latent state of environment from feedback^74–76^. Here, we propose that this inference can be achieved in neural circuits through prospective configuration, where following feedback, neurons in “hidden layers” converge to a prospective pattern of activity that encodes this latent state. We demonstrate that data from various previous studies, which involved the inference of a latent state, can be explained by prospective configuration. These data were previously explained by complex and abstract mechanisms, such as Bayesian models^74, 75^, while here we mechanistically show with prospective configuration how such inference can be performed by minimal networks encoding only the essential elements of the tasks.

The dynamical inference of latent state from feedback has been recently proposed to take place during sensorimotor learning^75^. In this experiment, participants received different motor perturbations in different contexts, and learned to compensate for these perturbations. Behavioural data suggest that after receiving the feedback, the participants were first employing it to infer the context, and then adapted the force for the inferred context. We demonstrate that prospective configuration is able to reproduce these behavioural data, while backpropagation cannot.

Specifically, in the task (Fig. 5a), participants were asked to move a stick from a starting point to a target point, while experiencing perturbations. The participants experienced a sequence of blocks of trials (Fig. 5c-e) including training, washout, and testing. During the training session, different directions of perturbations, positive (+) or negative (-), were applied in different contexts, blue (B) or red (R) backgrounds, respectively. We denote these trials as B+ and R-. These trials may be associated with latent states, which we denote by [B] and [R]; e.g., the latent state [B] may be associated with both background B and perturbation +. The next stage of the task was designed to investigate if this latent state [B] can be activated by the perturbation + even if no background B is shown. Thus, participants experienced different trials including R+ (i.e., perturbation + but no background B). Specifically, following a washout session (during which no perturbation was provided), in the testing session the participants experienced one of the four possible test trials: B+, R+, B-, and R-. To evaluate learning on the test trials, motor adaptation (i.e., the difference between the final and target stick positions) was measured before and after the test trial, on two trials with blue background (Fig. 5e). The change of the adaptation between these two trials is a reflection of learning about blue context that occurred at the test trial. If participants just associated feedback with the colour of background (B), then the change of adaptation would only occur with test trials B+ and B-. However, experimental data (Fig. 5f, right) show that there was substantial adaptation change also with R+ trials (which was even bigger than with B-trials).

**Fig. 5.**
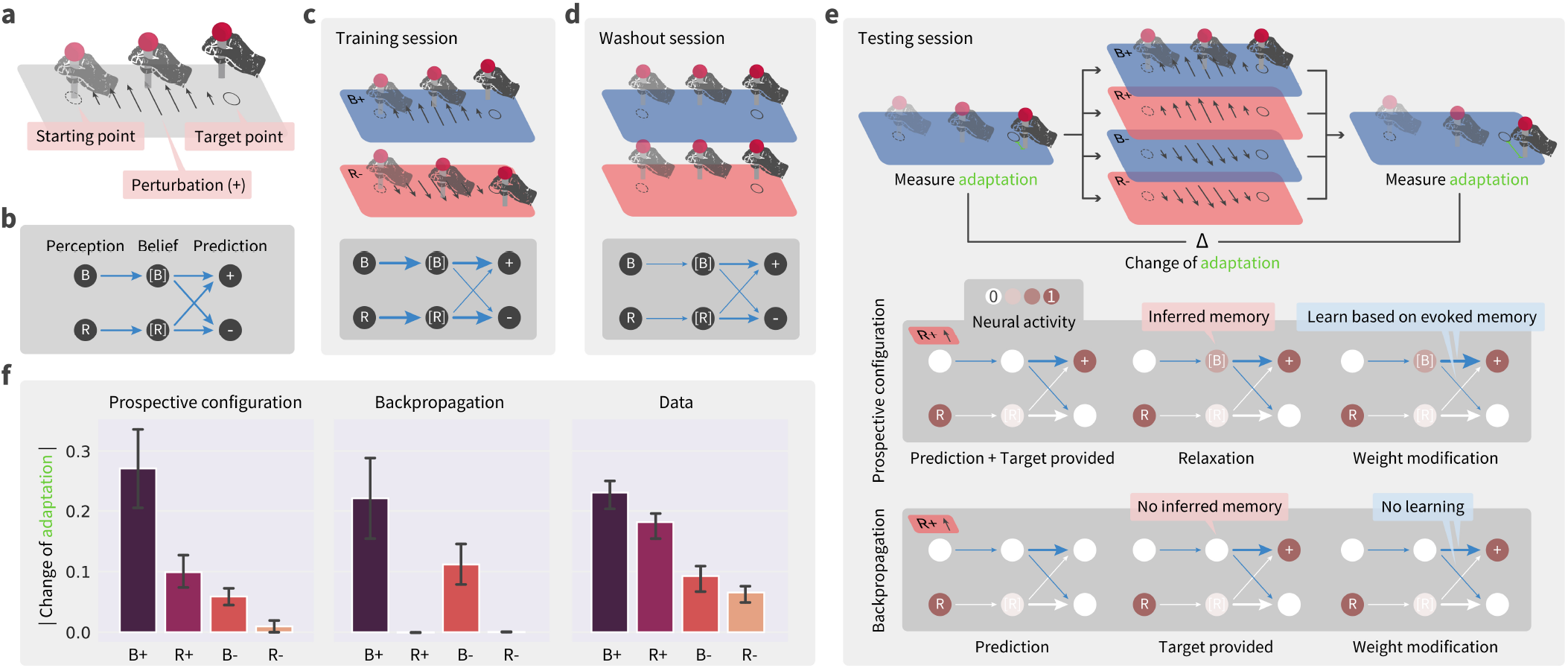
Prospective configuration explains contextual inference in human sensorimotor learning. ▶ **a** | The structure of an experimental trial, where participants were asked to move a stick from the starting point to the target point while experiencing perturbations. ▶ **b** | The minimal network for this task, including six connections encoding the associations from the backgrounds (B and R) to the belief of contexts ([B] and [R]), as well as from belief of contexts to prediction of perturbations (+ and -). ▶ **c-e** | A sequence of sessions the participants experienced: training, washout, and testing. Inside each panel, the darker box demonstrates the expected network after the session, where thickness represents the strength of connections. In the testing session, the darker box explains how the two learning rules learn differently on the R+ trial, leading to the differences in panel f. ▶ **f** | Predictions of the two learning rules compared against behavioural data measured from human participants, where prospective configuration reproduces the key patterns of data but backpropagation cannot.

To model learning in this task, we consider a neural network (Fig. 5b) where input nodes encode the colour of background, and outputs encode movement compensations in the two directions. Importantly, this network also includes hidden neurons encoding belief of being in the contexts associated with the two backgrounds ([B] and [R]). Trained with the exact procedure of the experiment^75^ from randomly initialized weights, prospective configuration with this minimal network can reproduce the behavioural data, while backpropagation cannot (cf., Fig. 5f left and middle).

Prospective configuration can produce change in adaptation with R+ test trial, because after + feedback, it is able to also activate context [B] that was associated with this feedback during training, and then learn compensation for this latent state. To shed light on how this inference takes place in the model, the bottom parts of Fig. 5c-d show evolution of the weights of the network over sessions (thickness represents the strength of connections). Fig. 5e bottom, shows the difference between the two learning rules at the exposure to R+: although B is not perceived, prospective configuration infers a moderate excitation of the belief of blue context [B], because the positive connection from [B] to + was built during the training session. The activity of [B] enables the learning of weights from [B] to + and -; while backpropagation does not modify any weights originating from [B].

For simplicity of explanation, we simulated the above experiment with minimal networks necessary to perform the task, but networks in the brain include multiple neurons, and it is important to establish if task structure that was reflected in these minimal networks can be discovered and learned by the networks themselves. Indeed, Extended Data Fig. 8 shows that networks with general fully-connected structure and more hidden neurons can replicate the above data on motor learning when employing prospective configuration, but not when using backpropagation. Thus, prospective configuration can discover task structure automatically and learn the task, while backpropagation cannot.

Studies of animal conditioning have also observed that feedback in learning tasks involving multiple stimuli may trigger learning about non-presented stimuli^77–81^. For example, in one study^77^ rats were trained to associate fear (electric shock) with noise and light; and then, in one group, fear related to light was eliminated in an extinction session (Fig. 6a). Remarkably, the data suggested that eliminating the fear to light increased the fear to noise (Fig. 6b). Such learning is not predicted by the standard Rescorla-Wagner model^82^. We consider a neural network (Fig. 6c) that includes two input neurons encoding the two stimuli, two hidden neurons, and one output neuron encoding the fear. Trained with the exact procedure of animal experiment^77^ from randomly initialized weights, prospective configuration with this simple network can reproduce the data, while backpropagation cannot (cf., Fig. 6b blue and orange). In the network employing prospective configuration, the feedback changes the activity of a hidden neuron previously associated with this feedback and with non-presented stimulus (noise), and hence enables modification of connections of this neuron (a learning mechanism analogous to that in sensorimotor learning Fig. 5, see Extended Data Figs. 9 for details).

**Fig. 6.**
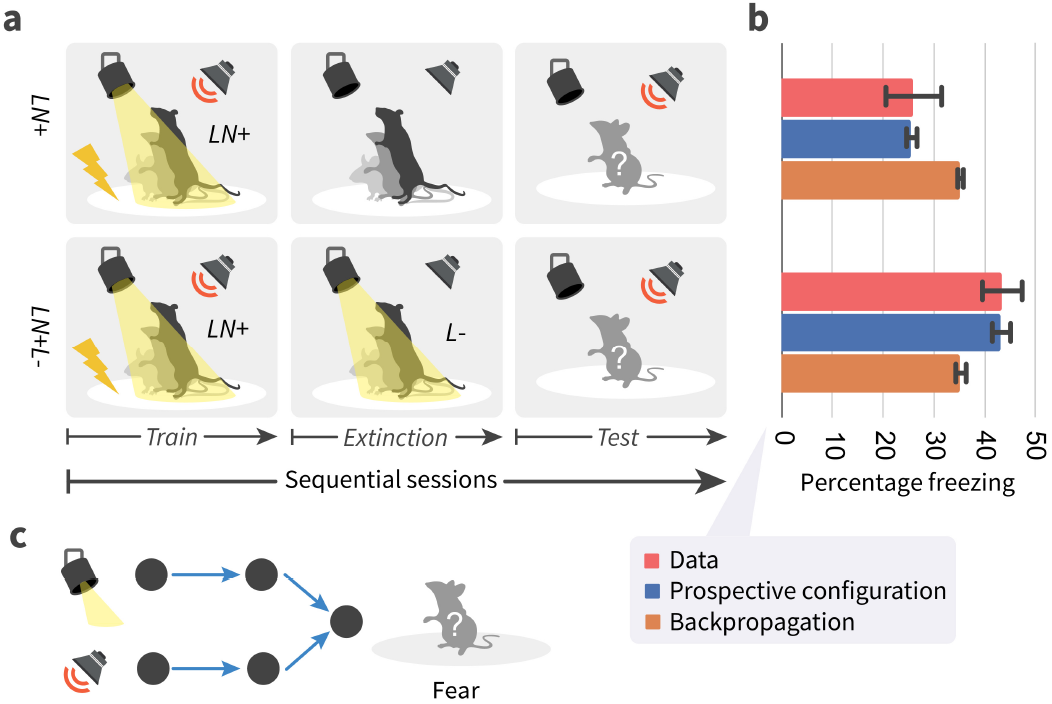
Prospective configuration infers latent state during fear conditioning. ▶ **a** | The fear conditioning task, where rats are first trained to associate fear (electric shock) with noise and light; then in one of the groups, fear related to light is eliminated in extinction session; finally, the predicted fear (percentage of rats freezing) of noise is measures in test session. ▶ **b** | The predicted fear from networks trained with prospective configuration and backpropagation, compared against the fear (percentage freezing) measured in rats. Prospective configuration reproduces the key finding that eliminating the fear to light changes the fear to noise. ▶ **c** | The architecture of simulated networks.

### Evidence for prospective configuration: discovering task structure during learning

Prospective configuration is also able to discover the underlying task structure in reinforcement learning. Particularly, we consider a task where reward probabilities of different options were not independent^74^. In this study humans were choosing between two options, whose reward probabilities were constrained such that one option had higher reward probability than the other (Fig. 7a). Occasionally the reward probabilities were swapped, so if one probability was increased, the other was decreased by the same amount. Remarkably, the recorded fMRI data suggested that participants learned that the values of two options were negatively correlated, and on each trial updated the value estimates of both options in opposite ways. This conclusion was drawn from the analysis of the signal from medial prefrontal cortex which encoded the expected value of reward. Fig. 7c, right compares this signal after making a choice on two consecutive trials: a trial on which reward was not received (“Punish trial”) and the next trial. If the participant selected the same option on both trials (“Stay”), the signal decreased, indicating the reward expected by the participant was reduced. Remarkably, if the participant selected the other option on the next trial (“Switch”), the signal increased, suggesting that negative feedback for one option increased the value estimate for the other. Such learning is not predicted by standard reinforcement learning models^74^.

**Fig. 7.**
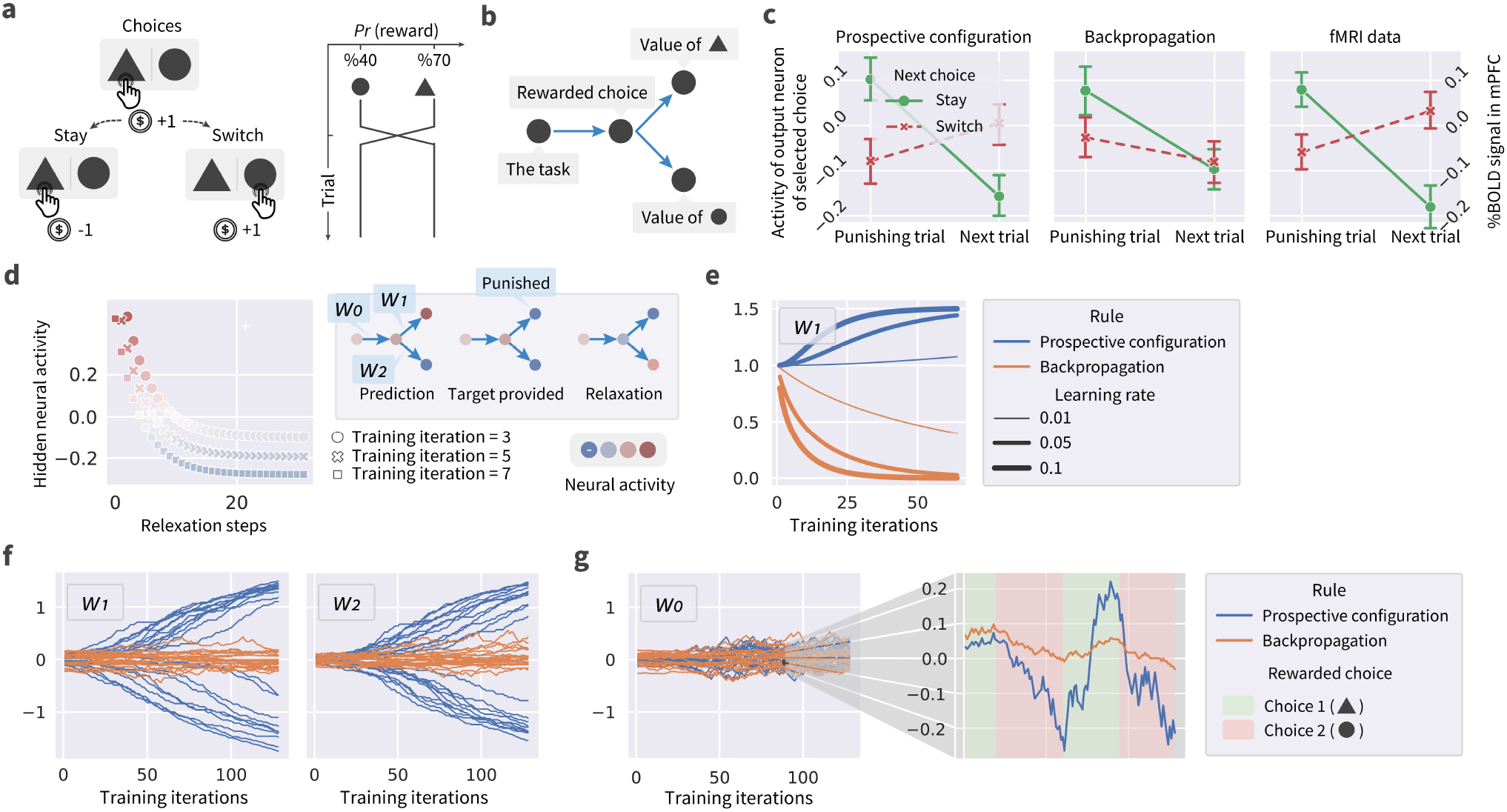
Prospective configuration can discover the underlying task structure during reinforcement learning. ▶ **a** | The reinforcement learning task, where human participants need to choose between two options, leading to either reward (gaining coins) or punishment (losing coins) with different probabilities. The probability of reward is occasionally reversed between the two options. ▶ **b** | The minimal network encoding the essential elements of the task. ▶ **c** | The activity of output neuron corresponding to the selected option, from networks trained with prospective configuration and backpropagation, compared against the fMRI data measured in human participants, i.e., peak blood oxygenation level-dependent (%BOLD) signal in the medial prefrontal cortex (mPFC). Prospective configuration reproduces the key finding that the expected value (encoded in %BOLD signal in mPFC) increases if the next choice after a punishing trial is to switch to the other option. ▶ **d** | Prospective configuration at the first few training iterations in an “idealized” version of the task: during relaxation, the hidden neuron is able to infer its prospective configuration, i.e., negative hidden activity encoding that the rewarded choice has reversed. ▶ **e** | Such inference by prospective configuration results in an increase of *W*_1_. By contrast, in backpropagation *W*_1_ is decreased. Similar behavior also applies to *W*_2_. ▶ **f** | The *W*_1_ and *W*_2_ in the simulation of the full task with stochastic rewards. Different lines correspond to different simulations. ▶ **g** | The evolution of *W*_0_ in the full task. In prospective configuration, this weight remains closer to 0 than *W*_1_ and *W*_2_. Inset shows *W*_0_ on one of the simulation in the main plot, where it is demonstrated that prospective configuration easily flips *W*_0_ as the rewarded choice changes, while backpropagation has difficulty in accomplishing this.

This task can be conceptualized as having a latent state encoding which option is superior, and this latent state determines the reward probabilities for both options. Consequently, we consider a neural network reflecting this structure (Fig. 7b) that includes an input neuron encoding being in this task (equal to 1 in simulations), a hidden neuron encoding the latent state, and two output neurons encoding the reward probabilities for the two options. Trained with the exact procedure of the experiment^74^ from randomly initialized weights, prospective configuration with this minimal network can reproduce the data, while backpropagation cannot (cf., Fig. 7c left and middle).

To shed light on the difference between the models, we simulate an “idealized” version of the task in Fig. 7d-e: the network shown in the inset starts from ({*W*_0_ = 1,*W*_1_ = 1,*W*_2_ = −1}) and is trained for 64 trials in total. The rewards and punishments are delivered deterministically, and the reversal only occurs once at the beginning of training. Fig. 7d inspects prospective configuration at the first few training iterations: during relaxation, the hidden neuron is able to infer its prospective configuration, i.e., negative hidden activity encoding that the rewarded choice has reversed. In Fig. 7e, such inference by prospective configuration results in an increase of *W*_1_: since it has inferred from the punishment that the rewarded choice has reversed to a non-rewarded one, such punishment strengthens the connection from the latent state representing non-rewarded choice to a punishment. By contrast, in backpropagation *W*_1_ is decreased: since it receives a punishment without updating the latent state (still encoding that the rewarded choice has not changed), it weakens the connection from the latent state to a reward. Fig. 7f shows the *W*_1_ and *W*_2_ in the simulation of the full task with stochastic rewards. The weights follow a similar pattern as in the simplified task, i.e., their magnitude increases in prospective configuration. This signifies that the network learns that the rewards from the two options are jointly determined by a hidden state. This increase of the magnitude of *W*_1_ and *W*_2_ enables the network to infer the hidden state from the feedback, and learn the task structure (as described for panel b). Fig. 7g shows the evolution of *W*_0_ in the full task. In prospective configuration, this weight remains closer to 0 than *W*_1_ and *W*_2_. Inset shows *W*_0_ on one of the simulation in the main plot, where it is demonstrated that prospective configuration easily flips *W*_0_ as the rewarded choice changes, while backpropagation has difficulty in accomplishing this. The reason of such behavior is as follows: thanks to large magnitude of *W*_1_ and *W*_2_ in prospective configuration, an error on the output unit results in a large error on the hidden unit, so the network is able to quickly flip the sign of *W*_0_ whenever the observation mismatches the expectation. This results in an increased expectation on the Switch trials (panel c).

Taken together, presented three simulations illustrate that prospective configuration is a common principle that can explain a range of surprising learning effects in diverse tasks.

## Discussion

Our paper identifies the principle of prospective configuration, according to which learning relies on neurons first optimizing their pattern of activity to match the correct output, and then reinforcing these prospective activities through synaptic plasticity. Although it was known that in energy-based networks the activity of neurons shifts before weight update, it has been previously thought that this shift is a necessary cost of error propagation in biological networks, and several methods have been proposed to suppress it^24, 25, 33, 47, 48^ to approximate backpropagation more closely. By contrast, we demonstrate that this reconfiguration of neural activity is the key to achieving learning performance superior to backpropagation, and to explaining experimental data from diverse learning tasks. Prospective configuration further offers a range of experimental predictions distinct from those of backpropagation (Extended Data Figs. 10–11). In sum, we have demonstrated that our novel credit assignment principle of prospective configuration enables more efficient learning than backpropagation by reducing interference, superior performance in situations faced by biological organisms, requires only local computation and plasticity, and can match experimental data across a wide range of tasks.

Our theory addresses a long-standing question of how the brain solves the plasticity-stability dilemma, e.g., how it is possible that despite learning and adjustment of representation in primary visual cortex^83^, we can still perceive the world and understand the meaning of visual stimuli we learned over our lifetime. According to prospective configuration, when some weights are modified during learning, compensatory changes are made to other weights, to ensure the stability of previously acquired knowledge. Previous computational models have also proposed mechanisms reducing interference between different pieces of learned information^72, 84^, and it is highly likely that these mechanisms operate in the brain in addition to prospective configuration and jointly reduce the interference most effectively.

From one view, prospective configuration could be seen as moving machine learning closer to inference and learning procedures in statistical modelling and system identification. For example, if the “energy” in energy-based schemes is variational free energy, i.e., the evidence lower bound (ELBO), prospective configuration can be seen as an implementation of variational Bayes that subsumes inference and learning^85–89^. Perhaps the closest example of this is dynamic expectation maximization^90, 91^. Dynamic expectation maximization (DEM) can be regarded as a generalization of predictive coding networks, in which the D-step optimizes representations of latent states (cf., relaxation till convergence), while the E-step optimizes model parameters (cf., weight modification). These two steps can be read as inference and learning respectively. This lends an interesting interpretation to prospective configuration, in the sense that the neuronal dynamics can be understood as inference (that prospectively precedes learning), while weight dynamics underwrite learning. This can be contrasted with backpropagation and amortization in standard machine learning approaches, which is limited to learning. In short, prospective configuration introduces inference into the optimization procedure to ensure optimal learning. It therefore shares with predictive coding networks a dual aspect optimization that can be regarded as a Bayesian filter with learnable parameters. One might ask what the M-step comprises in DEM. This corresponds to optimization of precision parameters that play the role of learning rates. In the computational neuroscience literature this corresponds to attention; namely, selecting precise prediction errors for local optimization of both neural and weight dynamics. We hope to consider this kind of extension by pursuing the close relationship between prospective configuration and (generalized) predictive coding networks in future work.

Other recent work^92, 93^ also noticed that the natural form of energy-based networks (“strong control” in their words) perform different learning comparing to backpropagation or approximations of backpropagation. Their analysis concentrates on an architecture of deep feedback control, and they demonstrated that particular form of their model is equivalent to predictive coding networks^93^. The unique contribution of our paper is to show the benefits of such strong control and explain why they arise.

Predictive coding networks require symmetric forward and backward weights between layers of neurons, so a question arises how such symmetry may develop in the brain. If predictive coding networks are initialized with symmetric weights (as in our simulations), the symmetry will persist, because the changes of a weight between neurons A and B are the same as for the feedback weight (between neurons B and A). Even if the weights are not initialized symmetrically, the symmetry may develop if synaptic decay is included in the model^94^, because then the initial asymmetric values decay away and weight values become more influenced by recent changes that are symmetric. Nevertheless the weight symmetry is not generally required for effective credit assignment, as it has been demonstrated that multilayer^26^ and recurrent^95^ neural networks can learn from errors propagated by feedback weights that are randomly generated, and hence asymmetric. Similarly weight symmetry is not essential for prospective configuration, as energy-based networks similar to predictive coding networks can work even if the weights are not symmetric^96^.

In this paper, we assumed for simplicity that the convergence of neural activity to an equilibrium happens rapidly after the stimuli are provided, so that the synaptic weight modification after convergence may take place while the stimuli are still present. However, the stimuli biological brains receive may be present very briefly or constantly change. Nevertheless, predictive coding networks can still work even if weight modification takes place while the neural activity is converging. Specifically, Song et al. demonstrate that if neural activities are only updated for the first few steps, the update of the weights is equivalent to that in backpropagation^33^. While, as a reminder, this manuscript demonstrates that if the neural activities are updated to equilibrium, the update of the weights follows the principle of prospective configuration, distinct from backpropagation and possesses the desirable properties demonstrated. Thus, a learning rule where neural activities and weights are updated in parallel will experience weights update that is equivalent to backpropagation at the start and then moves to prospective configuration as the system converges to equilibrium. We call this variant *parallel predictive coding*, which has been extensively studied in the Chapter 5 of the thesis from Song^97^. Furthermore, predictive coding networks have been extended to describe recurrent structures^98–100^, and it has been shown that such networks can learn to predict dynamically changing stimuli even if weights are modified before the activity converged for a given “frame” of the stimulus^99^.

The advantages of prospective configuration suggest that it may be profitably applied in machine learning to improve the efficiency and performance of deep neural networks. An obstacle for this is that the relaxation phase is computationally expensive. However, in recent work on parallel predictive coding we demonstrated that by modifying weights after each step of relaxation, the model becomes comparably fast as backpropagation and easier for parallelization^97^. Another approach to making energy-based networks more computationally efficient is to train them to predict their state following the relaxation^101^. Most intriguingly, it has been demonstrated that the speed of energy-based networks can be greatly increased by implementing the relaxation on analog hardware^102, 103^, potentially resulting in energy-based network being faster than backpropagation. Therefore, we anticipate that our discoveries may change the blueprint of next-generation machine learning hardware — switching from the current digital tensor base to analog hardware, being closer to the brain and potentially far more efficient.

## Methods

This section provides necessary details for replication of results in the main text.

### Models

Throughout this work, we compare the established theory of *backpropagation* to the proposed new principle of *prospective configuration*. As explained in the main text, backpropagation is used to train *artificial neural networks* (ANNs), where the activity of a neuron is *fixed* to a value based on its input, while prospective configuration occurs in *energy-based networks* (EBNs), where the activity of a neuron is *not* fixed.

Since in ANNs the activity of neurons ***x*** is determined by their input, the output of the network can be obtained by propagating the inputs “forward” through the computational graph. The output can then be compared against a target pattern to get a measure of difference known as a *loss*. Since the value of a node (activity of a neuron) in the computational graph is explicitly computed as a function of its input, the computational graph is usually differentiable. Thus, training ANNs with backpropagation modifies the weights ***w*** to take a step towards the negative gradient of loss *L*,

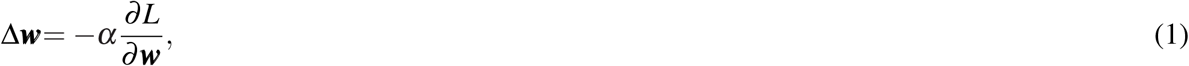

during which the activity of neurons ***x*** is fixed, and *α* is learning rate. The weights ***w*** requiring modification might be many steps away from the output on the computational graph, where the loss *L* is computed; thus, 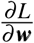 is often obtained by applying the chain rule of computing a derivative through intermediate variables (activity of output and hidden neurons). For example, consider a network with 4 layers and let ***x***^*l*^ denote the activity of neurons in layer *l*, while ***w***^*l*^ denote the weights of connections between layers *l* and *l* + 1. Then the change in the weights originating from the first layer is computed: 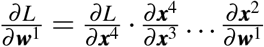. This enables the loss to be backpropagated through the graph to provide a direction of update for all weights.

In contrast to ANNs, in EBNs, the activity of neurons ***x*** is not fixed to the input from a previous layer. Instead, an energy function *E* is defined as a function of the neural activity ***x*** and weights ***w***. For networks organized in layers (considered in this paper), the energy can be decomposed into a sum of local energy terms *E*^*l*^:

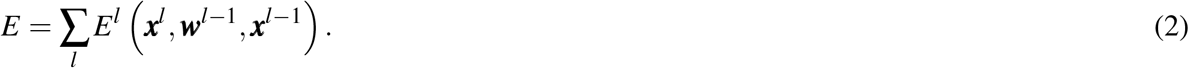

Here, *E*^*l*^ is called local energy, because it is a function of ***x***^*l*^, ***x***^*l*−1^, and ***w***^*l*−1^ that are neighbours and connected to each other. This ensures that the optimization of energy *E* can be implemented by local circuits, because the derivative of *E* with respect to any neural activity (or weights) results in an equation containing only the local activity (or weights) and the activity of adjacent neurons. Predictions with EBNs are computed by clamping the input neurons to an input pattern, and then modifying the activity of all other neurons to decrease the energy:

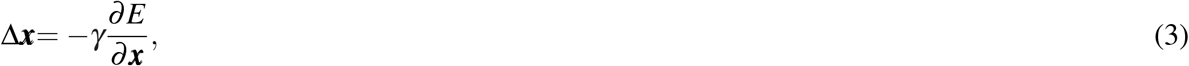

where *γ* is the integration step of the neural dynamics. Since the terms in *E* can be divided into local energy terms, this results in an equation that can be implemented with local circuits. This process of modifying the neural activity to decrease the energy is called *relaxation*, and we refer to the equation describing relaxation as *neural dynamics* — because it describes the dynamics of the neural activity in EBNs. After convergence of relaxation, the activities of the output neurons are taken as the prediction made by the EBN. Different EBNs are trained in slightly different ways. In case of *predictive coding network*^25, 40, 52^ (PCN), training involves clamping the input and output neurons to input and target patterns, respectively. Then, relaxation is run until convergence 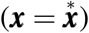, after which the weights are updated using the activity at convergence to further decrease the energy:

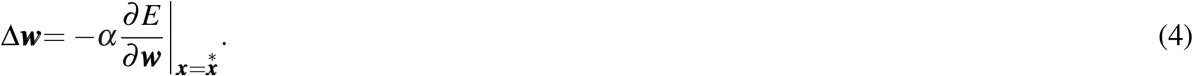

This will also result in an equation that can be implemented with local plasticity since it is just a gradient descent on the local energy. We refer to such an equation as *weight dynamics*, because it describes the dynamics of the synaptic weights in EBNs.

Backpropagation and prospective configuration are not restricted to specific models. Depending on the structure of the network, and the choice of the energy function, one can define different models that implement the principle of backpropagation or prospective configuration. In the main text and most of the Extended Data, we investigate the most standard layered network. In this case, both ANNs and EBNs include *L* layers of weights ***w***^1^, ***w***^2^, …, ***w***^*L*^, and *L* + 1 layers of neurons ***x***^1^, ***x***^2^, …, ***x***^*L*+1^, where ***x***^1^ and ***x***^*L*+1^ are the input and output neurons, respectively. We consider the relationship between activities in adjacent layers for ANNs given by

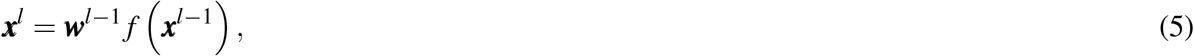

and the energy function for EBNs described by

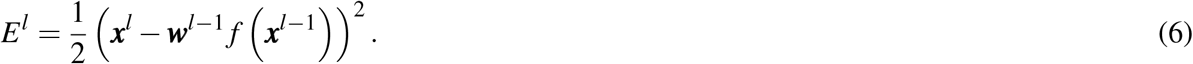

This defines the ANNs to be the standard *multilayer perceptrons* (MLPs) and the EBNs to be the PCN. In Eq. (6) and below, (***ν***)^2^ denotes the inner product of vector ***ν*** with itself. The comparison between backpropagation and prospective configuration in the main text is thus between the above MLPs and PCNs. This choice is justified by that (1) they are the most standard models^104^ and also (2) it is established that they two are closely related^25, 33^ (i.e., they make the same prediction with the same weights and input pattern), thus enabling a fair comparison. Nevertheless, we show that the theory (Extended Data Figs. 5) and empirical comparison (Extended Data Figs. 6 and 7) between backpropagation and prospective configuration generalize to other choices of network structures and energy functions, i.e., other EBNs and ANNs, such as *GeneRec*^105^ and *Almeida-Pineda*^106–108^.

Putting Eqs. (5) and (6) into the general framework, we can obtain the equations that describe MLPs and PCNs, respectively. Assume the input and target patterns are ***s***^in^ and ***s***^target^, respectively. Prediction with MLPs is:

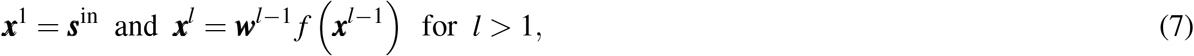

where ***x***^*L*+1^ is the prediction. Training MLPs with backpropagation is described by:

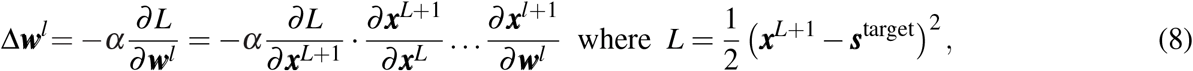

which backpropagates the error 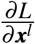 layer by layer from output neurons.

The neural dynamics of PCNs can be obtained using Eq. (2):

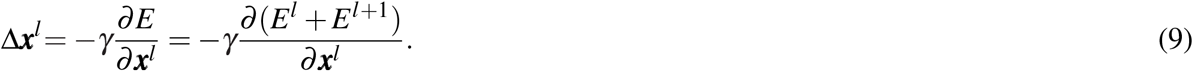

Similarly, the weight dynamics of PCNs can be found:

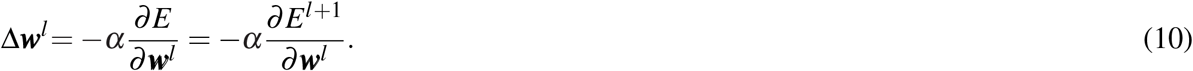

To reveal the neural implementation of PCN, we define the prediction errors to be

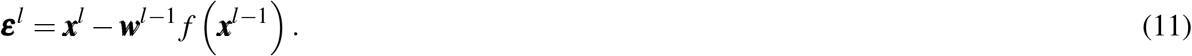

The neural and weight dynamics of PCN can be expressed (by evaluating derivatives in Eqs. (9) and (10)):

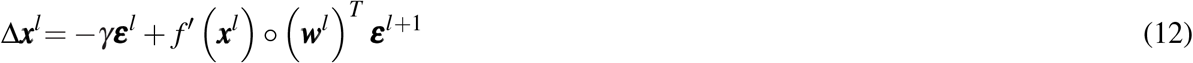

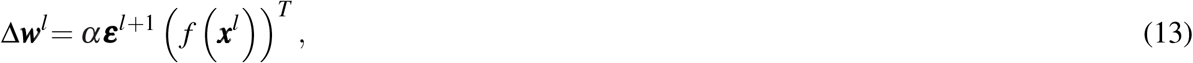

where the symbol denotes element-wise multiplication. Assuming that ***ε***^*l*^ and ***x***^*l*^ are encoded in the activity of error and value neurons, respectively, Eqs. (11) and (12) can be realized with the neural implementation in Fig. 2c bottom. Particularly, error ***ε*** and value ***x*** neurons are represented by red and blue nodes, respectively; excitatory + and inhibitory − connections are represented by connections with solid and hollow nodes, respectively. Thus, Eqs. (11) and (12) are implemented with red and blue connections, respectively. It should also be noticed that the weight dynamics is also realized locally: weight change described by Eq. (13) corresponds to simple Hebbian plasticity^109^ in the neural implementation of Fig. 2c bottom, i.e., the change in a weight is proportional to the product of activity of pre-synaptic and post-synaptic neurons. Thus, a PCN, as an EBN, can be implemented with local circuits only, due to the local nature of energy terms (as argued earlier in this section).

Full algorithm of PCN is summarized in Algorithm 1. In all simulations in this paper (unless stated otherwise), the integration step of the neural dynamics (i.e., relaxation) is set to *γ* = 0.1, and the relaxation is performed for 128 steps (𝒯 in Algorithm 1). During the relaxation, if the overall energy is not decreased from the last step, the integration step is reduced by 50%; if the integration step is reduced two times (i.e., reaching 0.025), the relaxation is terminated early. By monitoring the number of relaxation steps performed, we notice that in most of the tasks we performed, the relaxation is terminated early at around 60 iterations.

#### Algorithm 1: Learn with *predictive coding network*^25, 40, 52^ (PCN)

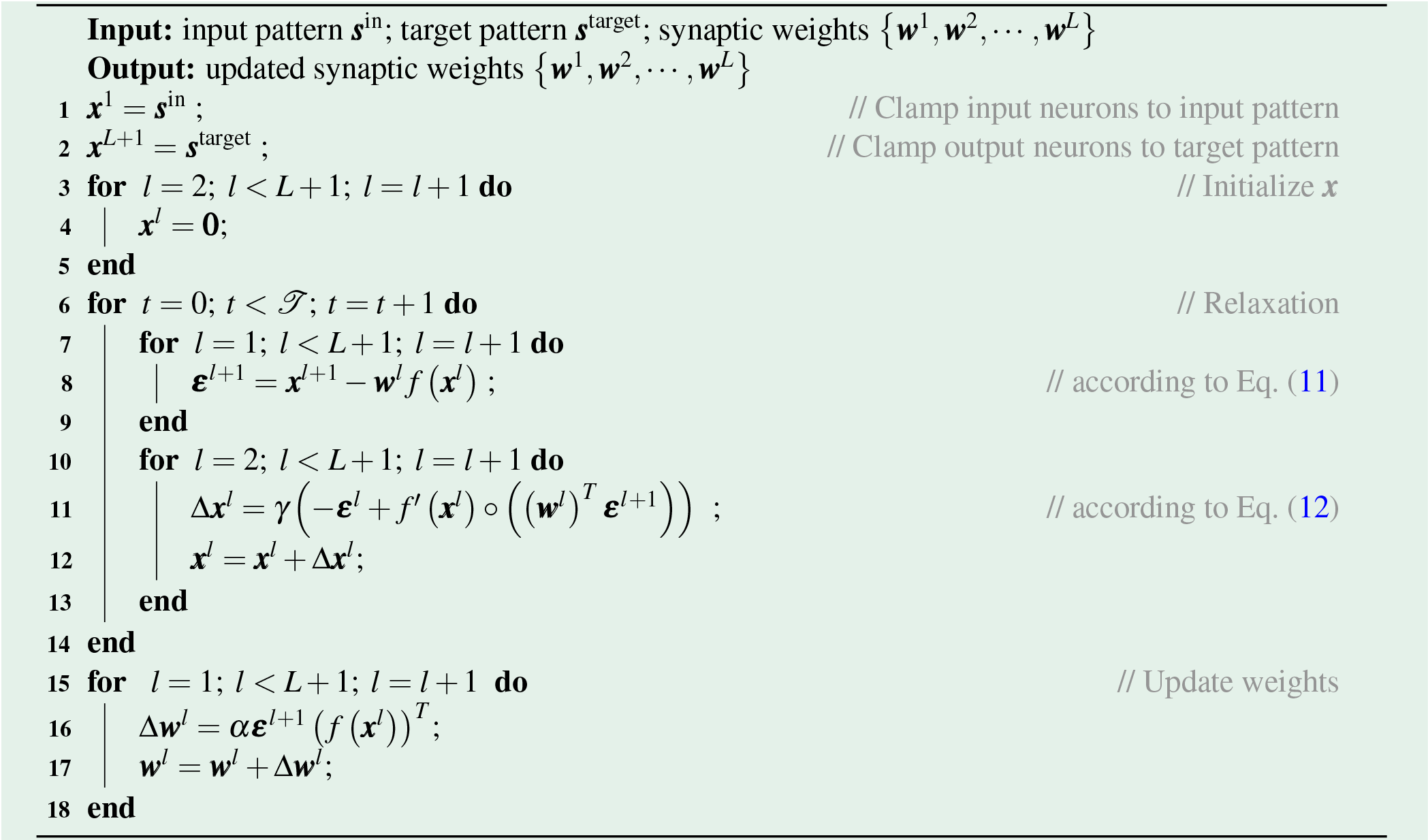

In the Extended Data, we also investigate other choices of network structures and energy functions, resulting in other ANNs and EBNs. Overall, the EBNs investigated include PCNs^25, 40, 52^, target-PCNs, and *GeneRec*^105^, and the ANNs investigated include backpropagation and *Almeida-Pineda*^106–108^. Details of all the models can be found in corresponding previous work, and are also given in the Supplementary Materials (Supplementary Information) 2.1.

### Interference and measuring interference (i.e., target alignment) (Fig. 3)

In Fig. 3a, since it simulates the example in Fig. 1, structure of the network is 1-1-2; weights are all initialized to 1; input pattern is [1] and target pattern is [0, 1]. Learning rates of both learning rules are 0.2, and the weights are updated for 24 iterations. Fig. 3d repeats the same experiment as Fig. 3a but with learning rate searched from (0.005, 0.01, 0.05, 0.1), which is wide enough to cover essentially all learning rates used to train deep neural networks in practice.

In Fig. 3e, there are 64 neurons in each layer (including input and output layers) for each network; weights are initialized via standard Xavier uniform initialization^110^. No activation function is used, i.e., linear networks are investigated. Depths of networks (*L*) are searched from {1, 2, 4, 6, 8, 10, 12, 14, 15}, as reported on the x-axis. Input and target patterns are a pair of randomly generated patterns of mean 0 and standard deviation 1. Learning rates of both learning rules are 0.001. Weights are updated for one iteration and target alignment is measured for this iteration for each of the 64 datapoints, then averaged over the 64 datapoints to produce the reported target alignment value. The whole experiment is repeated 3 times and the error bars report the standard error.

Simulations in Fig. 3f–h follow the setup of experiments in Fig. 4a–h, thus, are described at the end of the next section.

### Biologically relevant tasks (Fig. 4)

In supervised learning simulations, fully connected networks in Fig. 4a–h are trained and tested on FashionMNIST^56^, and convolutional neural networks^64^ (i–j) are trained and tested on CIFAR-10^65^. With FashionMNIST, models are trained to perform classification of gray-scaled fashion item images into 10 categories such as trousers, pullovers and dresses. FashionMNIST is chosen because it is of moderate and appropriate difficulty for multi-layer non-linear deep neural networks, so that the comparisons with EBNs are informative. Classification of data in CIFAR-10 is more difficult, as it contains colored natural images belonging to categories such as cars, birds and cats, thus only evaluated with convolutional neural networks. Both datasets consist of 60000 training examples (i.e., training set) and 10000 test examples (i.e., test set).

The experiments in Fig. 4a–h follow the configurations below, except for the parameters investigated in specific panels (such as batch size, size of the dataset, and size of the architecture), which are adjusted as stated in the description of specific experiments. The neural network is composed of 4 layers and 32 hidden neurons in each hidden layer. Note that the state-of-the-art MLP models of FashionMNIST are all quite large^111^. However, they are highly overparameterized, and thus, are not suitable to base our comparison on, because the accuracy reaches more than 95% regardless of the learning rule, due to the overparameterization. Thus, there is no space for demonstrating any meaningful comparison in these state-of-the-art overparameterized models. Overall, the size of the model on FashionMNIST demonstrated in this paper is a reasonable choice, with baseline models reaching reasonable performance (∼0.12 test error for standard machine learning setup) while keeping enough room for demonstrating performance difference in different learning rules. The size of the input layer is 28 × 28 for FashionMNIST^56^ gray-scaled. The size of the output layer is 10, as the number of classes for both datasets. The weights are initialized from a normal distribution with mean 0 and standard deviation 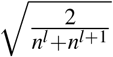, where *n*^*l*^ and *n*^*l*+1^ are the number of neurons of the layer before and after the weight, respectively. This initialization is known as Xavier normal initialization^110^. The activation function *f* () is *Sigmoid*. We define one *iteration* as updating the weights for one step based on a mini-batch. The number of examples in a mini-batch, called the batch-size, is by default 32. One *epoch* comprises presenting the entire training set, split over multiple mini-batches. At the end of each epoch, the model is tested on the test set and the classification error is recorded as the “test error” of this epoch. The neural network is trained for 64 epochs; thus, ending up with 64 test errors. The mean of the test error over epochs, i.e., during training progress, is an indicator of how fast the model learns. The minimum of the test errors over epochs is an indicator of how well the model can learn, ignoring the possibility of over-fitting due to training for too long. Learning rates are searched independently for each configuration and each model. Each experiment is repeated 10 times (unless stated otherwise), and the error bars represent standard error.

We now describe settings specific to individual experiments. In Fig. 4b different batch sizes are tested (as shown on x-axis). In Fig. 4c the batch size is set to 1. In continual learning of Fig. 4d, training alternates between two tasks. Task 1 is classifying five randomly selected classes in a dataset, and task 2 is classifying the remaining five classes. The whole network is shared by the two tasks, thus, differently from the network used in other panels, the network only has 5 output neurons. This better corresponds to continual learning with multiple tasks in nature, because, for example, if humans learn to perform two different tasks, they typically use the one brain and one pair of hands (i.e., the whole network is shared), since they do not have two different pairs of hands (i.e., humans share the output layers across tasks). Task 1 is trained for 4 iterations and then task 2 is trained for 4 iterations, and the training continues until total of 84 iterations is reached. After each iteration, error on the test set of each task is measured, as “test error”. In Fig. 4e, the mean of test error of both tasks during training of Fig. 4d at different learning rates is reported. In Figs. 4f–g investigating concept drifting^63, 112, 113^, changes to class labels are made every 512 epochs, and the models are trained for 4096 epochs in total. Thus, every 512 epochs, 5 out of 10 output neurons are selected, and the mapping from these 5 output neurons to the semantic meaning is pseudo-randomly shuffled. In Fig. 4h different numbers of data points per class (data points per class) are included into the training set (subsets are randomly selected according to different seeds).

In Fig. 4i, we train a convolutional network with prospective configuration and backpropagation, with the structure detailed in Fig. 4j. For each learning rule, we independently searched 7 learning rates ranging from {0.0005, 0.00025, 0.0001, 0.000075, 0.00005, 0.000025, 0.00001}. Both learning rules are trained for 80 epochs, with batch size 200. Weight decay of 0.01 is applied for both learning rules. Each configuration (each learning rule and each learning rate) are repeated for three times with different seeds.

To extend PCN to a convolutional neural network (or to any network with layered structure^34, 100^), we can define the forward function of a layer (i.e., how input of layer *l* + 1 is computed from the neural activity of layer *l*) with weights ***w***^*l*^ to be 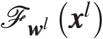. For example, for the MLPs described above, 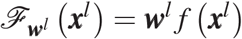. For convolutional network 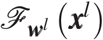 is a more complex function of ***w***^*l*^ and ***x***^*l*^, and also ***w***^*l*^ and ***x***^*l*^ are not simple matrix and vector anymore (to be defined later). Defining an ANN with ℱ () would be (i.e., Eq. (5) becomes): 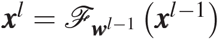. Defining energy function of PCN with ℱ () would be (i.e., Eq. (6) becomes): 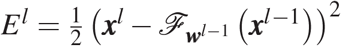. Thus, neural and weight dynamic would be (i.e., Eqs. (12) and (13) become): 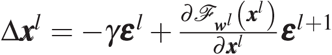 and 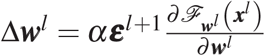, respectively. As 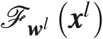 is defined, 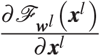 and 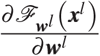 are obtained via auto differentiation in PyTorch (https://pytorch.org/tutorials/beginner/basics/autogradqs_tutorial.html). Thus, training a convolutional PCN is as simple as replacing Lines 11 and 16 in Algorithm 1 with the above corresponding equations.

In the following, we define 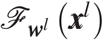 for convolutional networks. First, 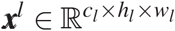, where *c*_*l*_, *h*_*l*_ and *w*_*l*_ are number of features, height and width of the feature map. These numbers for each layer are presented in Fig. 4j in the format of: *c*_*l*_@*h*_*l*_ × *w*_*l*_. For example, for the first layer (input layer), the shape is 3@32 × 32 as it is 32 × 32 colored images, i.e., with three feature maps representing red, green and blue. We denote kernel size, stride and padding of this layer as *k*_*l*_, *s*_*l*_ and *p*_*l*_, respectively. These numbers for each layer are presented in Fig. 4j. Thus, 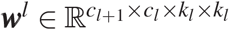. Finally, ***x***^*l*+1^ is obtained via:

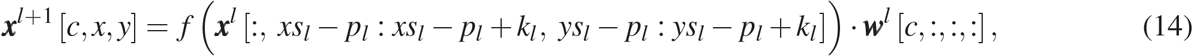

where [*a, b*, …] means indexing the tensor along each dimension, : means all indexes at that dimension, *a* : *b* means slice of that dimension from index *a* to *b* − 1, and · is dot product. In the above equation, if the slicing of ***x***^*l*^ on the second and third dimensions, i.e., ***x***^*l*^ [:, *xs*_*l*_ − *p*_*l*_ : *xs*_*l*_ − *p*_*l*_ + *k*_*l*_, *ys*_*l*_ − *p*_*l*_ : *ys*_*l*_ − *p*_*l*_ + *k*_*l*_] is outside its defined range 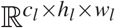, the entries outside range are considered to be zeros, known as padding mode of *zeros*.

In Fig. 3f, networks of 15 layers are trained and tested on FashionMNIST^56^ dataset. Learning rates in this Fig. 3f are optimized independently by a grid search over (5.0, 1.0, 0.5, 0.1, 0.05, 0.01, 0.005, 0.001, 0.0005, 0.0001, 0.00005, 0.00001, 0.000005) for each learning rule, as shown Fig. 3g, i.e., each learning rule in Fig. 3f uses the learning rate that gives minimal point in the corresponding curve in Fig. 3g. Fig. 3h investigates other network depths ({1, 2, 4, 6, 8, 10, 12, 14, 15}) in the same setup. Similarly as Fig. 3f, the learning rate for each learning rule and each “number of layers” is the optimal value (in terms of mean of test error as the y axis of the figure) independently searched from (5.0, 1.0, 0.5, 0.1, 0.05, 0.01, 0.005, 0.001, 0.0005, 0.0001, 0.00005, 0.00001, 0.000005). Hidden layers are always of size 64 in the above experiments. In the above experiment, only part of the training set was used (60 datapoints per class) so that the test error is evaluated more frequently to reflect the difference on efficiency of the investigated learning rules. The activation function *f* () used is *LeakyReLU*, instead of the standard Sigmoid, because Sigmoid results in difficulty in training deep neural networks. Other unmentioned details follows the defaults as described above.

In the reinforcement learning experiments (Fig. 4k), we evaluate performance on three classic reinforcement learning problems: Acrobot^114, 115^, MountainCar^116^, and CartPole^117^. We interact with these environments via a unified interface by OpenAI Gym^118^. The observations *s*_*t*_ of these environments are vectors describing the status of the system, such as velocities and positions of different moving parts (for details refer to the original articles or documentation from OpenAI Gym). Each entry of the observation *s*_*t*_ is normalized to mean 0 and standard deviation 1 via Welford’s online algorithm^119, 120^. The action space of these environments is discrete. Thus, we can have a network taking in observation *s*_*t*_ and predicting the value (*Q*) of each action *a*_*t*_ with different output neurons. Such a network is known as an action-value network, in short, a *Q* network. In our experiment, the *Q* network contains two hidden layers, each of which contains 64 neurons, initialized the same way as the network used for supervised learning, described before. One can acquire the value of an action *a*_*t*_ at a given observation *s*_*t*_ by feeding in *s*_*t*_ to the *Q* network and reading out the prediction on the output neuron corresponds to the action *a*_*t*_, such value is denoted by *Q* (*s*_*t*_, *a*_*t*_). The training of *Q* is a simple regression problem to target 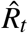, obtained via *Q*-learning with experience replay (summarized in Algorithm 2). Considering *s*_*t*_ to be ***s***^in^ and 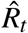 to be ***s***^target^, the *Q* network can be trained with prospective configuration or backpropagation. Note that 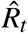 is the target of the selected action *a*_*t*_ (i.e., the target of one of the output neurons corresponds to the selected action *a*_*t*_), thus, 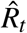 is in practice considered to be ***s***^target^ [*a*_*t*_]. For prospective configuration, it means the rest of the output neurons except the one corresponding to *a*_*t*_ are freed; for backpropagation, it means the error on these neurons are masked out.

PCN of slightly different settings from the defaults is used for prospective configuration: the integration step is fixed to be half of the default (=0.05), and relaxation is performed for a fixed and smaller number of steps (=32). This change is introduced because *Q*-learning is more unstable (so smaller integration step) and more expensive (so smaller number of relaxation steps) than supervised learning tasks. To produce a smoother curve of “Sum of rewards per episode” in Fig. 4k from the *SumRewardPerE pisode* in Algorithm 2, the *SumRewardPerE pisode* curve along *TrainingE pisode* are averaged with a sliding window of length 200. Each experiment is repeated with 3 random seeds and the shadows represents standard error across them. Learning rates are searched independently for each environment and each model from the range {0.05, 0.01, 0.005, 0.001, 0.0005, 0.0001}; and the results reported in Fig. 4k are for the learning rates yielding the highest mean of “Sum of rewards per episode” over training episodes.

#### Algorithm 2: *Q*-learning with experience replay.

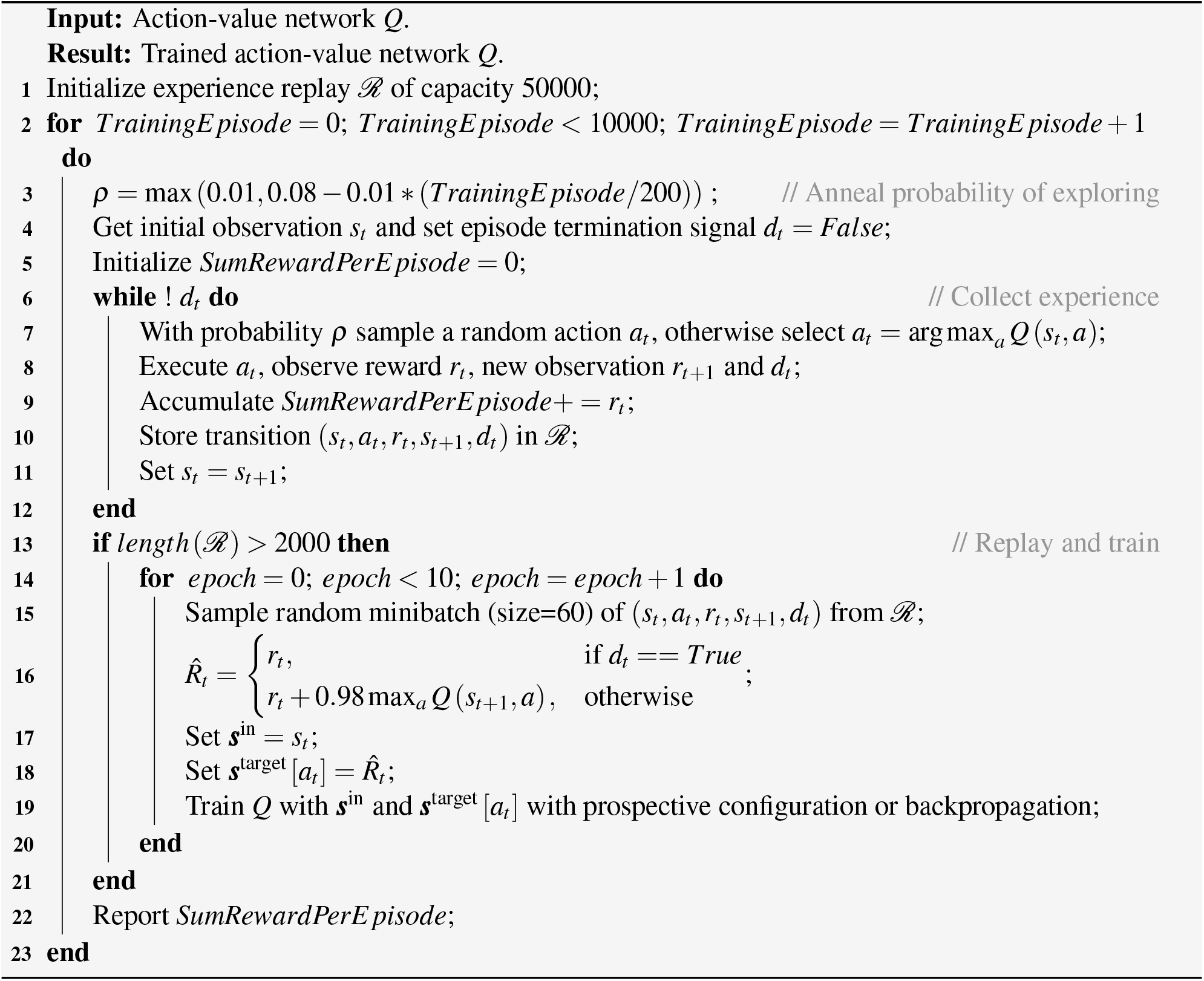

### Simulation of motor learning (Fig. 5)

As shown in Fig. 5, we train a network that includes 2 input, 2 hidden, and 2 output neurons. The two input neurons are one-to-one connected to the two hidden neurons, and the two hidden neurons are fully connected to the two output neurons. The two input neurons are considered to encode presenting the blue and red background, respectively. The two output neurons are considered to encode the prediction of the perturbations towards positive and negative directions, respectively. Presenting or not presenting a background color are encoded as 1 and 0 respectively; presenting or not presenting perturbations of a particular direction are encoded as 1 and 0, respectively. The weights are initialized from a normal distribution with mean 0 and standard deviation fitted to behavioural data (see below), simulating that the participants have not built any associations before the experiments. Learning rates are independent for the two layers, as we expect the connections from perception to belief and the connections from belief to predictions to have different degree of plasticity. The two learning rates are also fitted to the data (see below).

The number of participants, training and testing trials follow exactly the human experiment^74^. In particular, for each of 24 simulated participants, the weights are initialized with a different seed of the random number generator. They each experience two stages: training and testing. Note that the pre-training stage performed in the human experiment is not simulated here as its goal was to make human participants familiar with the setup and devices.

In the training stage, the model experiences 24 blocks of trials. In each block, the model is presented with the following sequence of trials, matching the original experiment^74^.

- The model is trained with two trials without perturbation: B0 and R0, with order counterbalanced across consecutive blocks. Note that in the human experiment there were two trial types without perturbations (channel and washout trials), but they are simulated in the same way here as B0 or R0 trials, because they both did not include any perturbations.
- The model is trained with 32 trials with perturbations, where there are equal number of B+ and R-within each 8 trials in a pseudorandom order.
- The model experiences two trials: B0 and R0, with order counterbalanced across consecutive blocks.
- The model experiences *n* ← {14, 16, 18} washout trials (equal number of B0 and R0 trials in a pseudorandom order), where *n* ← {*a, b, c*} denotes sampling without replacement from a set of values *a, b* and *c*, and replenishing the set whenever becomes empty.
- The model experiences one triplet, where the exposure trial is either B+ or R-, counterbalanced across consecutive blocks. Here, a triplet consists three sequential trials: B0, the specified exposure trial and again B0.
- The model experiences again *n* ← {6, 8, 10} washout trials (equal number of B0 and R0 trials in a pseudorandom order).
- The model experiences again one triplet, where the exposure trial is either B+ or R-, whichever was not used on the previous triplet.

Then, in the testing stage, the model experiences 8 repetitions of four blocks of trials. In each block, one of combinations B+, R+, B- and R- is tested. The order of the four blocks is shuffled in each of the 8 repetitions. In each block, the model first experiences *n* ← {2, 4, 6} washout trials (equal number of B0 and R0 trials in a pseudorandom order). Then the model experiences a triplet of trials, where the exposure trial is the combination (B+, R+, B- or R-) tested in a given block, to assess single trial learning of this combination. The change in adaption in the model is computed as the absolute value of the difference in the predictions of perturbations on the two B0 trials in the above triplet, where the prediction of perturbation is computed as the difference between the activities of the two output neurons. The predictions are averaged over participants and the above repetitions.

The parameters of each learning rule are chosen such that the model best reproduces the change in adaptation shown in Fig 5f. In particular, we minimize the sum over set *C* of the 4 exposure trial types of the squared difference between average change in adaptation in experiment (*d*_*c*_) and in the model (*x*_*c*_):

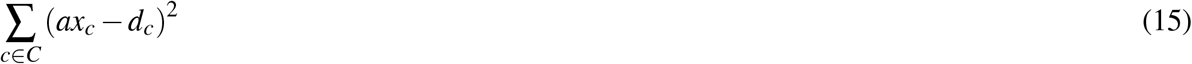

The model predictions are additionally scaled by a coefficient *a* fitted to the data, because the behavioural data and model outputs have different scales. Exhaustive search was performed over model parameters: standard deviation of initial weights could take values from {0.01, 0.05, 0.1}, and two learning rates for two layers could take values from {0.00005, 0.0001, 0.0005, 0.01, 0.05}. Then, for each learning rule and each combination of the above model parameters, the coefficient *a* is resolved analytically (restricted to be positive) to minimize the sum of the squared errors of Eq. (15).

### Simulation of fear conditioning (Fig. 6)

As shown in Fig. 6c, the simulated network includes 2 input, 2 hidden, and 1 output neurons. The weights are initialized from a normal distribution of mean 0 and standard deviation 0.01, reflecting that the animals have not built an association between stimulus and electric shock before the experiments. Presenting or not presenting the stimulus (noise, light, or shock) is encoded as 1 and 0, respectively. The two input neurons are considered to be the visual and auditory neurons; thus, their activity corresponds to perceiving light and noise, respectively. The output neuron is considered to encode the prediction of the electric shock. The training and extinction sessions are both simulated for 32 iterations with the learning rate of 0.01. In the test session, the model makes a prediction with the presented stimulus (noise only). As in the previous section, we denote by *x*_*c*_ the prediction for each group *c* from a set *C* = {*N*+, *LN*+, *LN* + *L*−}. To map the prediction to the percentage of freezing, it is scaled by a coefficient *a* (as the neural activity and the measure of freezing have different units) and shifted by a bias *b* (as the rats may have some tendency to freeze after salient stimuli even if they had not been associated with a shock). The numbers reported in Fig. 6b are these scaled predictions. The coefficient *a* (constrained to be positive) and bias *b* are optimized for prospective configuration and backpropagation independently, analogously as described in the previous section, i.e. their values that minimize summed squared error given below are found analytically.

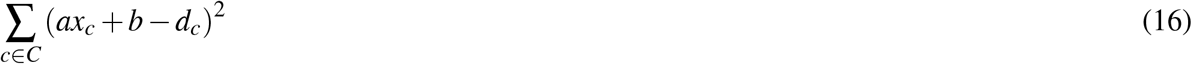

### Simulation of human reinforcement learning (Fig. 7)

As shown in Fig. 7b, we train a network that includes 1 input, 1 hidden, and 2 output neurons. The input neuron is considered to encode being in the task, so it is set to 1 throughout the simulation. The two output neurons encode the prediction of the value of the two choices. Reward and punishment are encoded as 1 and −1, respectively, because the participants were either winning or losing money. The model selects actions stochastically based on the predicted value of the two choices (encoded in the activity of two output neurons) according to the softmax rule (with temperature of 1). The weights are initialized from a normal distribution of mean 0 and standard deviation fitted to experimental data (see below), simulating that the human participants have not built any associations before the experiments. Number of simulated participants (number of repetitions with different seeds) was set to 16 as in the human experiment^74^. The number of trials is not mentioned in the original paper, so we simulate for 128 trials for both learning rules.

To compare the ability of the two learning rules to account for the pattern of signal from mPFC, for each of the rules, we optimized the parameters describing how the model is set up and learns (the standard deviation of initial weights and the learning rate). Namely, we searched for the values of these parameters for which the model produces the most similar pattern of its output activity to that in the experiment. In particular, we minimized the sum over set *C* of four trial types in Fig. 7c of the squared difference between model predictions *x*_*c*_ and data *d*_*c*_ on mean mPFC signal (Eq. (16)). The model predictions are additionally scaled by a coefficient *a* and offset by a bias *b*, because the fMRI signal had different units and baseline than the model. To compute the model prediction for a given trial type, the activity of the output neuron corresponding to the chosen option is averaged across all trials of this type in the entire simulation. The scaled average activity from the model is plotted in Fig. 7c, where the error bars show the standard error of the scaled activity. To fit the model to experimental data, the values of model parameters and the coefficient were found analogously as described in the previous section. In particular, we employ exhaustive grid search on the parameters. The models are simulated for all possible combinations of standard deviation of initial weights, and the learning rate, from the following set: {0.01, 0.05, 0.1}. Then, for each learning rule and each combination of the above model parameters, the coefficient *a* (restricted to be positive) and the bias *b* are resolved analytically to minimize sum of the squared error of Eq. (16).

## Data availability

Learning tasks analysed in Fig. 4a-j were built using the publicly available FashionMNIST^56^ and CIFAR-10^65^ datasets. They are incorporated in most machine learning libraries, and their original releases are available at https://github.com/zalandoresearch/fashion-mnist and https://www.cs.toronto.edu/~kriz/cifar.html, respectively. Reinforcement learning tasks analysed in Fig. 4k were built using the publicly available simulators by OpenAI Gym^118^.

## Code availability

Complete code and full documentation reproducing all simulation results will be made publicly available at https://github.com/YuhangSong/A-New-Perspective upon publication of this work. It will be released under GNU General Public License v3.0 without any additional restrictions (for license’s details see https://opensource.org/licenses/GPL-3.0 by the open source initiative).

## Acknowledgements

We thank Timothy Behrens for comments on the manuscript, and Andrew Saxe for discussions. Yuhang Song was supported by the China Scholarship Council under the State Scholarship Fund and J.P. Morgan AI Research Awards. Beren Millidge and Rafal Bogacz were supported by the the Biotechnology and Biological Sciences Research Council grant BB/S006338/1 and Medical Research Council grant MC UU 00003/1. Thomas Lukasiewicz and Tommaso Salvatori were supported by the Alan Turing Institute under the EPSRC grant EP/N510129/1 and by the AXA Research Fund. Zhenghua Xu was supported by National Natural Science Foundation of China under the grant 61906063, by the Natural Science Foundation of Hebei Province, China, under the grant F2021202064, by the Natural Science Foundation of Tianjin City, China, under the grant 19JCQNJC00400, by the “100 Talents Plan” of Hebei Province, China, under the grant E2019050017, and by the Yuanguang Scholar Fund of Hebei University of Technology, China.

## JPMorgan Chase & Co

This research was funded in part by JPMorgan Chase & Co. Any views or opinions expressed herein are solely those of the authors listed, and may differ from the views and opinions expressed by JPMorgan Chase & Co. or its affiliates. This material is not a product of the Research Department of J.P. Morgan Securities LLC. This material should not be construed as an individual recommendation for any particular client and is not intended as a recommendation of particular securities, financial instruments or strategies for a particular client. This material does not constitute a solicitation or offer in any jurisdiction.

## Author contributions

Y.S. and R.B. conceived the project. Y.S., R.B., B.M. and T.S. contributed ideas for experiments and analysis. Y.S. and B.M. performed simulations. Y.S., B.M. and R.B. performed mathematical analyses. Y.S., T.L, and R.B. managed the project. T.L, and Z.X. advised on the project. Y.S., R.B. and B.M. wrote the paper. T.S., T.L, and Z.X. provided revisions to the paper.

## Additional information

**Extended Data Figures/Tables** is available for this paper in the same file (Section 1).

**Supplementary Information** is available for this paper in the same file (Section 2).

**Correspondence and requests** for materials should be addressed to Y.S. and R.B.

## 1 Extended Data

**Extended Data Fig. 1.**
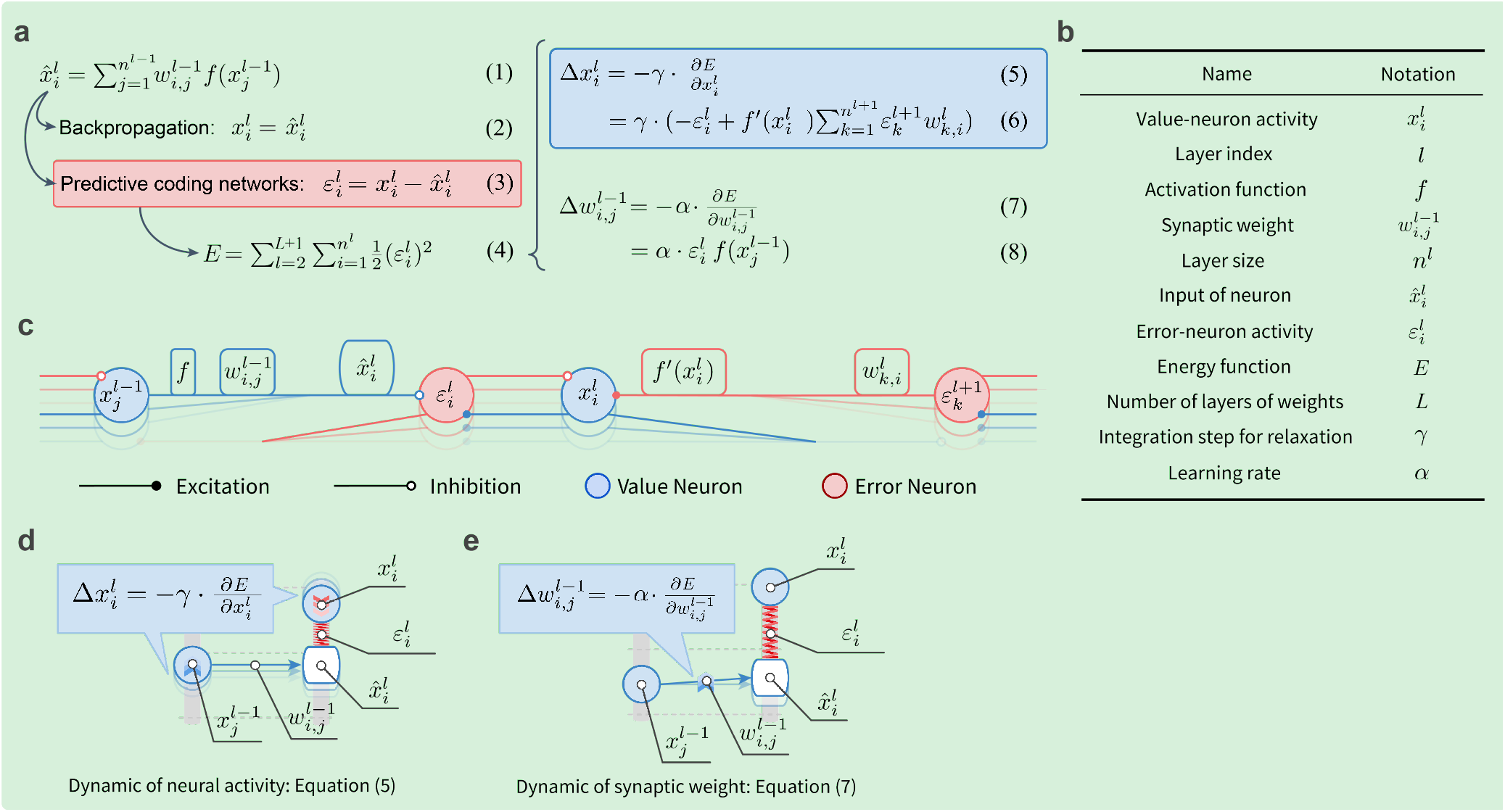
Predictive coding networks, neural implementation and corresponding energy machine. The figure shows a list the equations describing the equilibrium-seeking dynamics and plasticity of predictive coding networks (panels a-b), how these equations map to a neural implementation, and how they map to the machine analog introduced in Fig. 2. ▶ **a** | List of equations describing predictive coding networks. Eq. (1) in this figure describes the input to a given layer 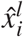 from the neuron in the previous layer 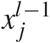. In artificial neural networks trained with backpropagation, neural activities of a given layer 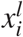 are set as the input to this layer 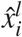 (Eq. (2)). In contrast, in predictive coding networks, neural activities of this layer 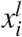 are **not** set as the input to this layer 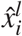, instead an error 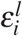 is defined between them (Eq. (3)). Additionally, predictive coding networks define the energy *E* of the network to be the sum of all the squared errors 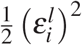 (Eq. (4)). The dynamic of neural activity 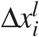 in predictive coding networks is set to change the neural activity in proportion to the negative gradient of the energy with respect to the neural activity, so as to reduce the energy (Eq. (5)), which can be further derived as Eq. (6). The dynamic of synaptic weights 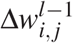 of predictive coding networks is set to be in proportion to the negative gradient of the energy with respect to the weight, so as to reduce the energy (Eq. (7)), which can be further derived as Eq. (8). ▶ **b** | A list of symbols shared by all panels in the figure for easy reference. ▶ **c** | Mapping of equations describing predictive coding networks in panel a to a neural implementation. The neural implementation includes value neurons (blue) performing computations in Eq. (6), and separate error neurons encoding prediction errors (red) performing computations in Eq. (3), where positive sign is encoded by excitatory connections while negative sign is encoded by the inhibitory connections. It should be noticed that the weight dynamics 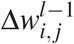 is also realized locally: weight change described by Eq. (8). corresponds to simple Hebbian plasticity^109^ in the architecture shown in panel a, i.e., the change in a weight is proportional to the product of activity of pre-synaptic and post-synaptic neurons. Different suggestions have been made on how this architecture could be realized in cortical circuits. An influential study^121^ has suggested that error and value neurons correspond to separate neurons, so in such architecture the plasticity rule is precisely Hebbian, as explained above. Some other models^22^ implementing predictive coding networks^32^ include an error compartment (in apical dendrite) and a value compartment (in soma) within a single neuron. In such architecture the plasticity is still local as it depends on the product of activity in one neurons and potential of the apical dendrite in the other neuron. ▶ **d-e** | Mapping of equations describing predictive coding networks in panel a to the machine analog introduced in Fig. 2. The exact same set of equations describing predictive coding networks also describe a physical machine connected with rods, nodes and springs. ▶ **d** | The dynamic of neural activity 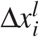 of predictive coding networks (Eq. (5)) describes relaxing the physical machine by moving the nodes. ▶ **e** | The dynamic of synaptic weight 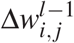 of predictive coding networks (Eq. (7)) describes relaxing the physical machine by tuning the rods.

**Extended Data Fig. 2.**
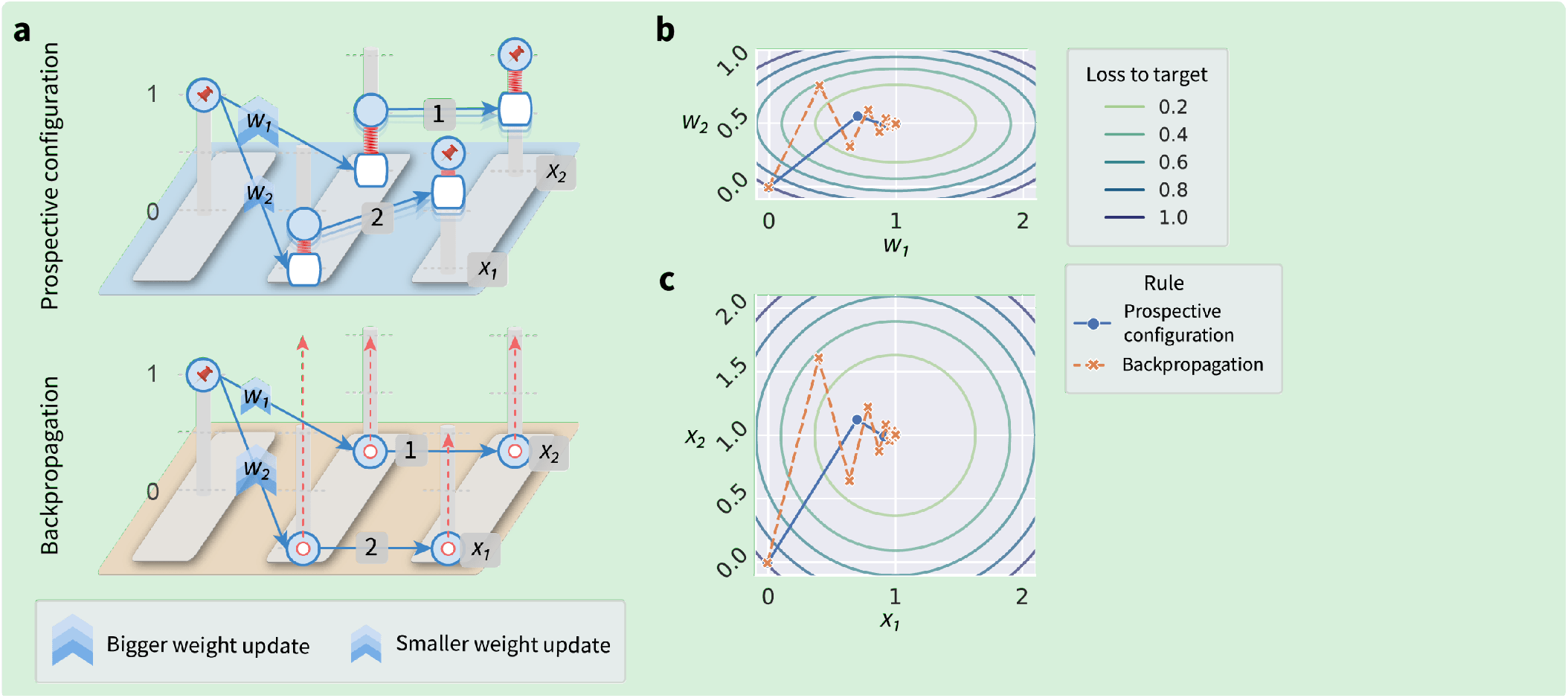
Differences in learning between prospective configuration and backpropagation. This figure shows an example of a simple network revealing striking differences in how errors are propagated and weights modified by the two algorithms. For this network it is possible to explicitly visualize how learning changes weights and outputs, and explicitly show that although backpropagation follows the gradient of loss in the space of weights, it does not in the space of outputs. ▶ **a** | Setup of the example. In this example, we consider a network consisting of 1 input neuron, 2 hidden neurons and 2 output neurons, with the structure shown with the energy machine. The input is always 1 and the target of both output neurons are both 1. The weights in the first layer are initialized to 0, while in the second layer to 1 (top) and 2 (bottom). We visualize with the energy machine how prospective configuration and backpropagation learn differently in this example. Prospective configuration assigns larger error to the top hidden neurons than the bottom, and hence would increase *w*_1_ more than *w*_2_. By contrast, backpropagation does the opposite: since the backpropagated errors are scaled by the weights to output layer, the error for the bottom hidden neuron is higher than for the top. Importantly, in this learning problem, weight *w*_2_ does not need to be modified as much as *w*_1_, because any changes in *w*_2_ will be amplified by the high weight to the output neuron. Prospective configuration indeed modifies *w*_2_ less than *w*_1_, while backpropagation does the opposite. This suggests that backpropagation does not modify the weights optimally to move output toward the target, and we will illustrate it in the following panels. ▶ **b** | Landscape of the weights (*w*_1_ and *w*_2_). We consider a setup in which the network only learns the two weights on the first layer: *w*_1_ and *w*_2_, while the weights in the second layer are fixed all the time during the training. This is so that the weight space is small (only two dimensional, so that we can visualize the landscape of weights); and we choose to learn the two weights in the first layer instead the second (last) layer so that the problem is not trivial. All the combinations of weights on the same contour line gives the same loss to the target (in short, loss), where we can see the combination of *w*_1_ = 1 and *w*_2_ = 0.5 gives loss of 0. Assuming the weights (*w*_1_, *w*_2_) start from (0, 0), backpropagation (orange) takes steps following the direction orthogonal to the contour lines, i.e. the direction of local gradient descent. It is well-known that backpropagation cannot have more global vision of the minimal point of the landscape: thus, often forms the trajectory of learning as the orange curve, “bouncing” towards the global minimum point. Prospective configuration (blue), on the contrary, although does not follow gradient in the weight space (blue line is not orthogonal to the contour lines), it moves more directly to the global minimum of the landscape. This is exactly due to the mechanism of prospective configuration giving the learning rule a more global view of the system: as mentioned above, prospective configuration infers that since the bottom weight of second layer is larger (= 2) than the top one (= 1), it only needs small error being assigned to the bottom neuron of the hidden layer so as to correct the error on the bottom output. ▶ **c** | Landscape of the outputs (*x*_1_ and *x*_2_). The panel shows changes in output neurons’ activity, *x*_1_ and *x*_2_, resulting from the weight updates in panel b. As in panel b, the contour lines indicate the loss. Comparing panels b and c reveals that changes of backpropagation (orange) are orthogonal to the loss contour lines in weight space, but not in output neuron space; while changes of prospective configuration (blue) are not orthogonal to loss contour lines in weight space, but are closer to being orthogonal in output neuron space. Overall, the comparison reveals fundamental difference between backpropagation and prospective configuration: backpropagation does local gradient decent in weight space (local means it only sees the infinitely small area around it current state); while prospective configuration infers the configuration of neuron activities that reduces the loss in the output space, thus, the trajectory in the weight space is fundamentally different from that for backpropagation. This fundamental difference leads to advantage of prospective configuration over backpropagation: it moves more directly towards the minimal point in the weight space and output space, instead of “bouncing” towards it (as backpropagation does). Learning rate in this panel is the same as the learning rate used in the corresponding learning rule in panel b. **Implementation details**. Learning rate for backpropagation in this figure is set to *α* = 0.4, while that for prospective configuration is solved so that it produces the same magnitude of weight change 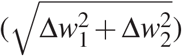 during the first iteration as backpropagation. Weights are updated for 15 iterations. Details of the learning rules are described in the Methods section and also in Supplementary Information 2.1.

**Extended Data Fig. 3.**
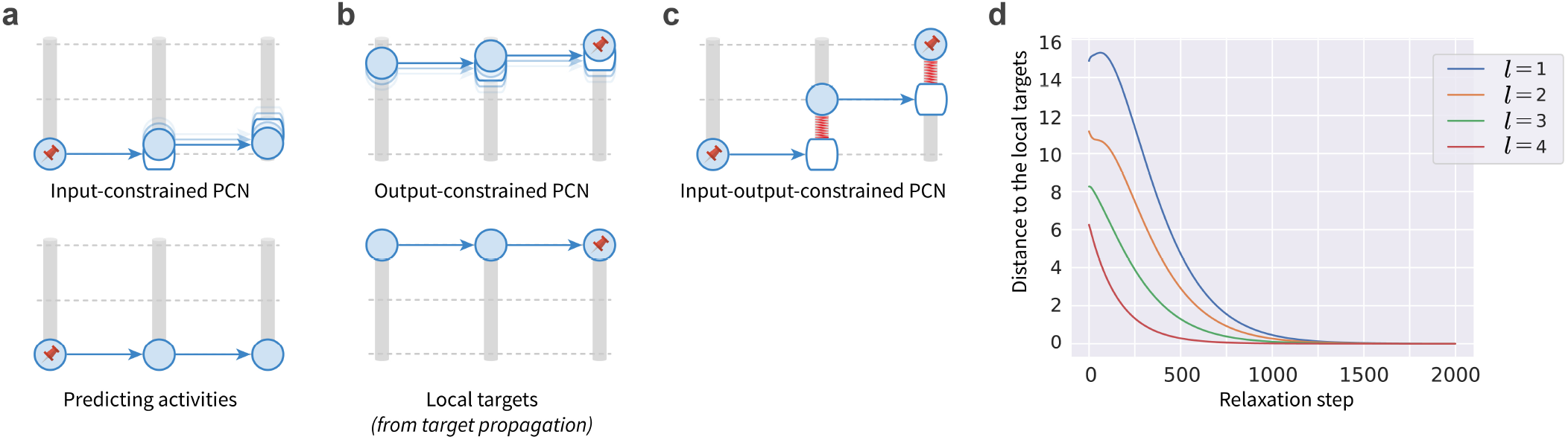
Relationship of prospective configuration to target propagation. Prospective configuration is related to another influential algorithm of credit assignment — target propagation^122^. Since target propagation has target alignment equal to 1^58^, this relationship provides an explanation for the high target alignment of prospective configuration. Target propagation is an algorithm, which explicitly computes the neural activity in hidden layers required to produce the desired target pattern. We call these values local targets. We demonstrate that one of energy-based networks, predictive coding networks^25, 40, 52^ (PCNs) tends to move the activity during relaxation towards these local targets. The relationship of PCNs to target propagation can be visualized with the proposed energy machine in Fig. 2, hence panels a–c illustrate how the neural activity in a PCN depends on whether inputs and outputs are constrained, and these properties are formally proved in Supplementary Information 2.2. ▶ **a** | With only input neurons constrained (and outputs unconstrained) PCNs can generate prediction about the output, and hence we refer to this pattern of neural activity as the predicting activity. ▶ **b** | With only output neurons constrained (and inputs unconstrained), the neural activity of PCNs relaxes to the local target from target propagation. This happens because with only outputs constrained, other nodes have a freedom to move to values that generate the outputs, and when the energy reduces to 0 (as shown in the bottom display) all neurons must have the activity generating the target output. ▶ **c** | With both input and output neurons constrained, the neural activity of PCNs relaxes to the weighted sum of the local target from target propagation and the predicting activity. Note that the position of the hidden node is in between the positions from panels a and b. ▶ **d** | The distance between the neural activity to the local target at different layers along the relaxation progress in output-constrained PCNs. Here, the neural activity of the output-constrained PCNs converges to the local target, and the layers closer to the output layer (larger *l*) converge to the local target earlier than the others, which is as expected from the physical intuition of the energy machine. **Implementation details**. We train the models to predict a target pattern from an input pattern (both randomly generated from 𝒩 (0, 1), and the input and target patterns are of 5 and 1 entries, respectively). The structure of the networks is 5→ 5→ 5→ 5→ 1. There is no activation function, i.e., it is a linear network. For the computation of the local target in target propagation, refer to the original paper^122^. The mean square difference is used to measure the distance to the local target.

**Extended Data Fig. 4.**
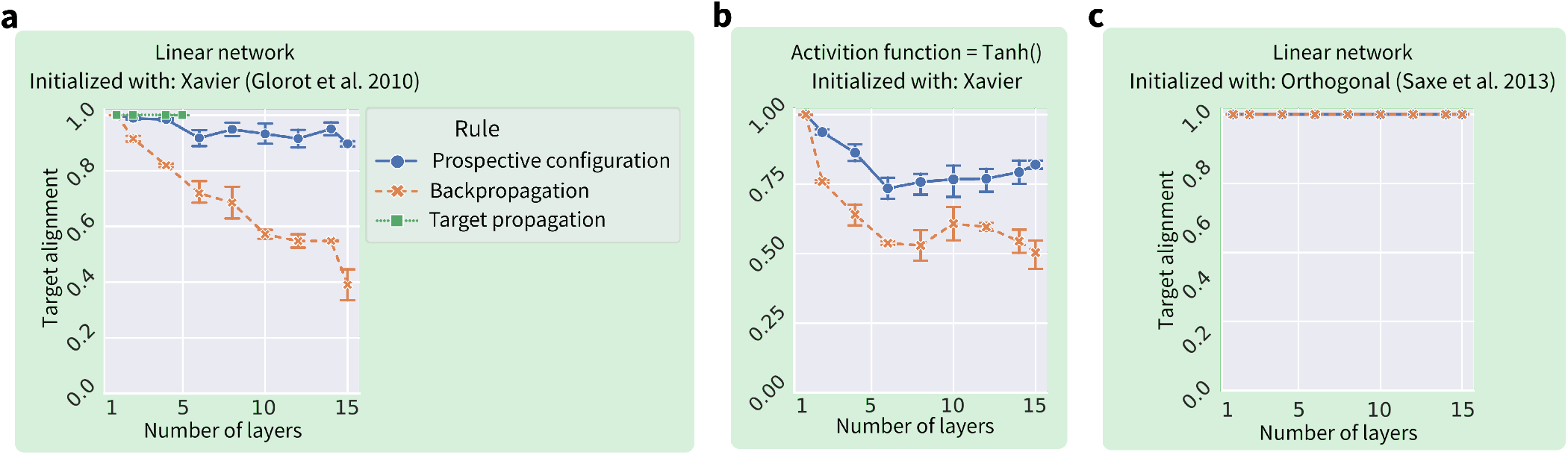
Target alignment in deep neural networks with different learning algorithms, non-linearities and initializations. This figure extends the analyses from Fig. 3e in the main paper of target alignment in randomly generated networks with different depth. ▶ **a** | Target alignment for target propagation in deep linear network initialized with standard Xavier normal initialization^110^. For comparison, the results presented in Fig. 3e of the main paper for predictive coding networks and backpropagation are also shown. The results for target propagation are only shown for networks with up to 5 layers, because the algorithm became numerically unstable for deeper networks. The target alignment of target propagation is equal to 1 as implied by previous analytic work^58^ (for details see section 2.4.2 of Supplementary Information). ▶ **b** | Target alignment for networks with a non-linear (*Tanh*) activation function, initialized with standard Xavier normal initialization^110^. The higher value of target alignment for predictive coding networks than backpropagation shown in panel a generalizes to networks with non-linearity. ▶ **c** | Target alignment of linear networks with orthogonal initialization (where weight in each layer satisfy (***w***^*l*^)^*T*^ ***w***^*l*^ = ***I***)^123^. Saxe et al.^123^ discovered that with such initialization weights evolve independently of each other during learning, thus, learning times can be independent of depth, even for arbitrarily deep linear networks. As shown in the figure, interestingly, orthogonal initialization gives target alignment of 1 for both learning rules. We also demonstrated this analytically in section 2.4.3 of Supplementary Information. This perfect target alignment can be intuitively expected, because the independence of weights mentioned above is related to a lack of interference, and it further illustrates that reduction in target alignment is caused by interference between weights.

**Extended Data Fig. 5.**
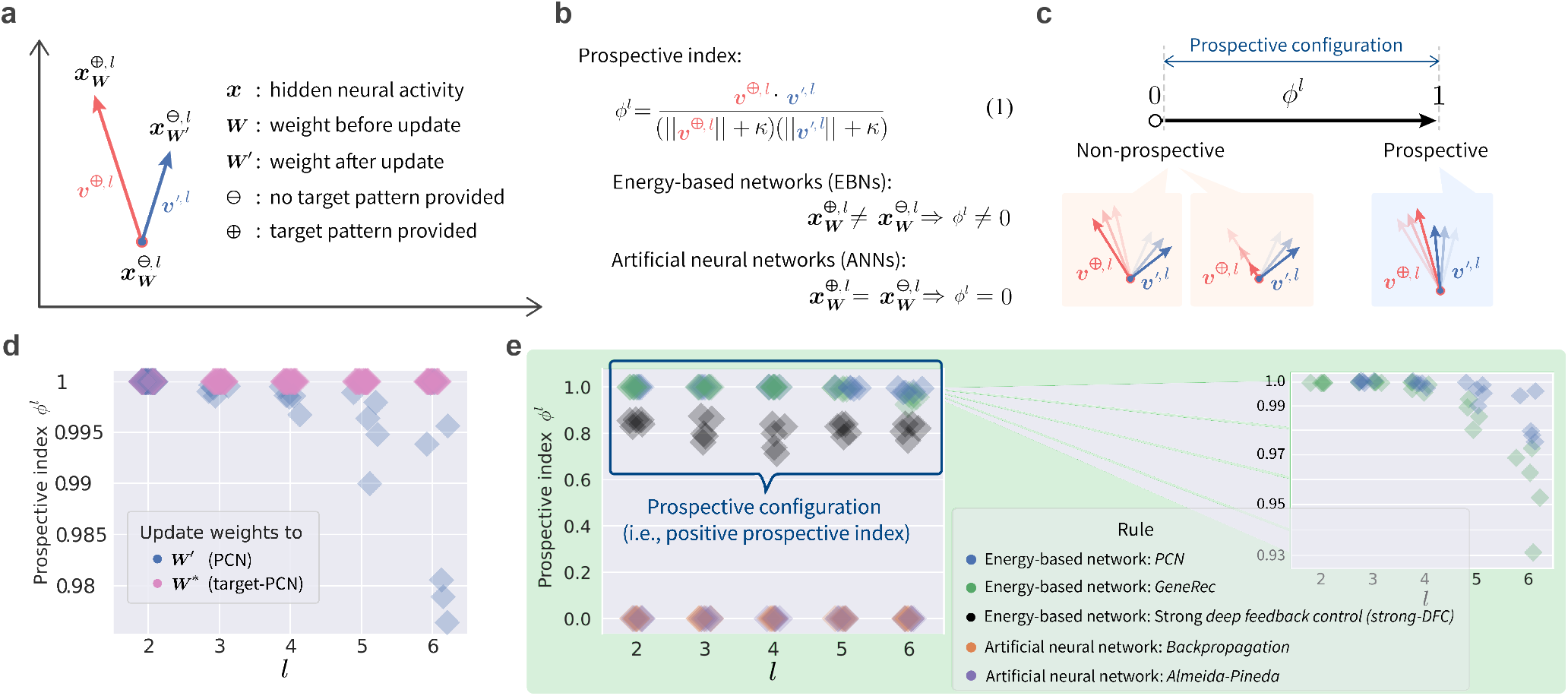
Formal definition of prospective configuration. Formal definition of prospective configuration with prospective index (panels a–c), a metric that one can measure for any learning model. With this metric, we show that prospective configuration is present in different *energy-based networks* (EBNs), but not in *artificial neural networks* (ANNs) (panels d–e). ▶ **a** | To introduce the prospective index, we consider the hidden neural activity ***x***^*l*^ in layer *l*, at three moments of time. First, a learning iteration starts from ***x***^*l*^ under the current weights ***W*** without target pattern provided 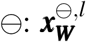. Second, a target pattern is provided ⊕, and neural activity settles to 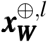. Third, ***W*** is updated to ***W*** ^′^, the target pattern is removed ⊖, and the neural activity settles to 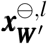. We define two vectors ***ν***^⊕,*l*^ and ***ν***^′,*l*^, representing the direction of the neural activity’s changes as a result of the target pattern being given ⊖ → ⊕ and the weights being updated ***W*** → ***W*** ^′^, respectively. ▶ **b** | The prospective index *ϕ*^*l*^ is the cosine similarity of ***ν***^⊕,*l*^ and ***ν***^′,*l*^. A small constant *κ* = 0.00001 is added in the denominator to ensure that the prospective index is still defined if the length of one of the vectors is 0 (in which case the prospective index in equal to 0). For EBNs, the neural activity settles to a new configuration when the target pattern is provided, i.e., 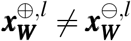, so *ϕ*^*l*^ is non-zero; for ANNs, the neural activity stays unchanged when the target pattern is provided, i.e., 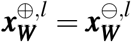, so *ϕ*^*l*^ is zero. ▶ **c** | A positive *ϕ*^*l*^ implies that ***ν***^⊕,*l*^ and ***ν***^′,*l*^ are pointing in the same direction, i.e., the neural activity after the target pattern provided 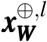 is similar to the neural activity after the weight update 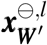, i.e., is prospective. We define the models following the principle of prospective configuration as those with positive *ϕ*^*l*^ (averaged over all layers). Additionally, prospective index close to 1 implies that a weight update rule in a model is able to consolidate the pattern of activity following relaxation, so a similar pattern is reinstated during prediction on the next trial. ▶ **d** | The prospective index *ϕ*^*l*^ of different layers *l* in PCNs and a variant of PCNs called target-PCNs. Several observations can be made, and they are explained and proved in Supplementary Information 2.3. ▶ **e** | The prospective index *ϕ*^*l*^ of different EBNs and ANNs. Here, we can see that all EBNs produce positive *ϕ*^*l*^, i.e., the prospective configuration is commonly observed in EBNs, but not in ANNs. Among the EBNs, *Deep Feedback Control*^124^ (DFC) was proposed to work with “infinitely weak nudging”, as in equilibrium propagation^24^. More recent work demonstrates that it also works with “strong control”^92, 93^ (thus, called strong-DFC), i.e., with the natural form of EBNs. The prospective index was measured for this strong-DFC model and shows it belongs to one of EBNs that process prospective configuration. Details of the simulated strong-DFC model can be found in Section 2.1 of Supplementary Information. **Implementation details**. We train various models to predict a target pattern from an input pattern (both randomly generated from 𝒩 (0, 1)). The structure of the networks is 64→ 64→ 64→ 64→ 64→ 64→ 64. The weights are initialized using Xavier normal initialization^110^ (described in the Methods). No activation function was used. Batch size is set to 1. The models were trained for one iteration (i.e., one update of the weights), the prospective index was then measured for this update. Prospective indices of input and output layers are not reported. This is because the input and output layers are held fixed during learning; thus, the prospective index is not defined for them. Experiments were repeated 5 times. The EBNs investigated include PCNs^25, 40, 52^, target-PCNs, and *GeneRec*^105^, while the ANNs investigated include backpropagation and *Almeida-Pineda*^106–108^. Details of all simulated models are given in Section 2.1 of Supplementary Information.

**Extended Data Fig. 6.**
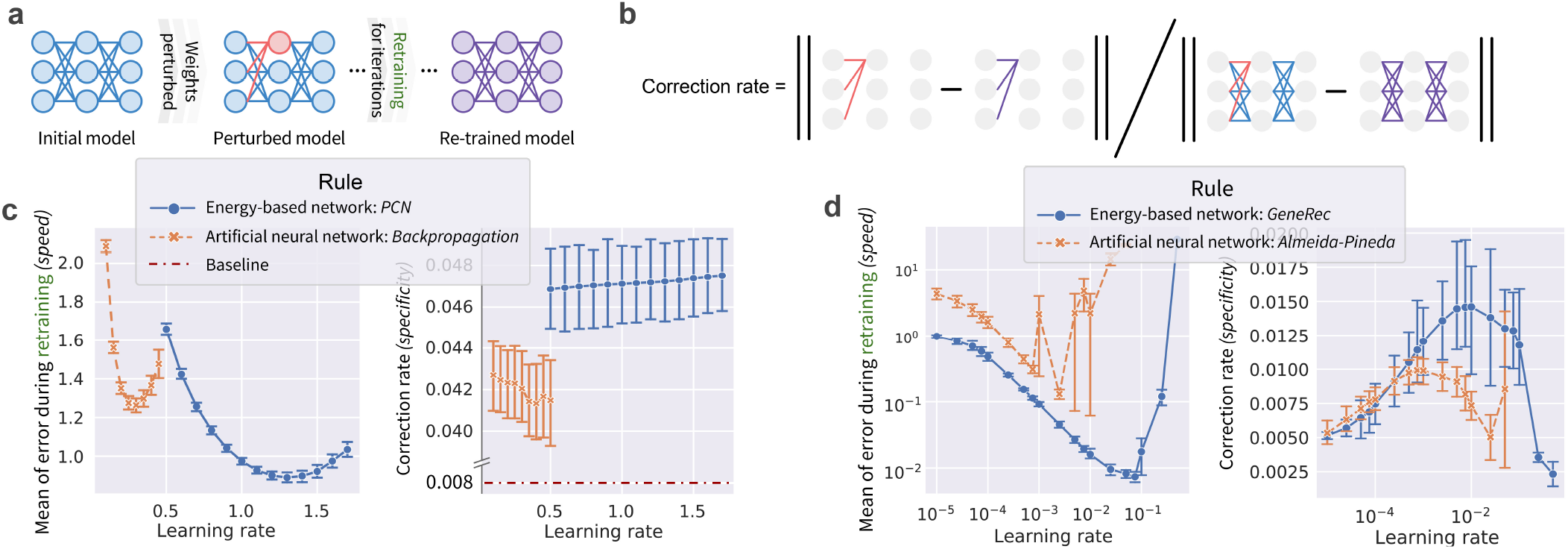
Prospective configuration yields a more accurate weight modification. A numerical experiment (panels a–b) verifies that *energy-based networks* (EBNs) yield a accurate weight modification than *artificial neural networks* (ANNs) (panels c–d). The following intuition can be provided for why the prospective configuration enables an accurate weight modification. In EBNs, if more error is assigned to a neuron, this neuron will settle to a prospective activity that reduces the error. The prospective activity of this neuron is then propagated through the network, resulting in less error being assigned to other neurons, thus the error being assigned more accurately. ▶ **a** | Experimental procedure: we take a pre-trained model (illustration here does not reflect the real size of the model), randomly select a hidden neuron and perturb the synaptic weights connecting to this neuron (red), then retrain this model on the same pattern for a fixed number of iterations. During retraining, an optimal learning agent is expected to identify that the error in the output neurons is due to the perturbed weights, thus, (1) correct the error faster, and (2) correct the perturbed weights more. We refer to the above two properties as *speed* and *specificity*. Speed can be measured with the mean of error over retraining iterations (the lower, the better). ▶ **b** | Specificity can be measured by correction rate (the higher, the better): the ratio of how much the perturbed weights are corrected compared to how much all the weights (in all layers) are corrected after all retraining iterations. ▶ **c** | A comparison between an EBN, predictive coding network^25, 40, 52^ (PCN), and an ANN, trained with backpropagation. In the right plot, there is an additional baseline, which is the number of perturbed weights divided by the number of all the weights, indicating the expected correction rate if a learning rule randomly assigns errors. ▶ **d** | The same comparison as in panel c, but for another EBN, namely, *GeneRec*^105^. GeneRec describes learning in recurrent networks, and ANN with this architecture is not trained by standard backpropagation, but by a variant of backpropagation, called *Almeida-Pineda*^106–108^. **Implementation details**. We first pre-train the models to predict a target pattern from an input pattern (both randomly generated from 𝒩 (0, 1) and of 32 entries). The structure of the networks is 32→ 32→ 32→ 32. The pre-training session is sufficiently long (1000 iterations) to reach convergence. Then, one neuron is randomly selected from the (32 + 32) hidden neurons, and all weights connecting to this neuron are “flipped” (i.e., multiplied by −1). Current weights of the network are recorded as ***W***_*b*_. The part of current weights that were just flipped are recorded as 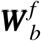. The network is then re-trained on the same pattern for 64 iterations. After each re-training iteration, the model makes a prediction. The square difference between the prediction and the target pattern is recorded as the “error during re-training” of this iteration. After the entire re-training session, the “errors during re-training” are averaged over the 64 re-training iterations, producing the left plots of panels c–d. Current weights of the network are recorded as ***W*** _*a*_. The part of current weights that were flipped before the re-training session are recorded as 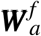. The *correction rate* is computed as 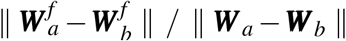, which produces the right plots of panels c–d. Each configuration was repeated 20 times, and the error bars represent standard error.

**Extended Data Fig. 7.**
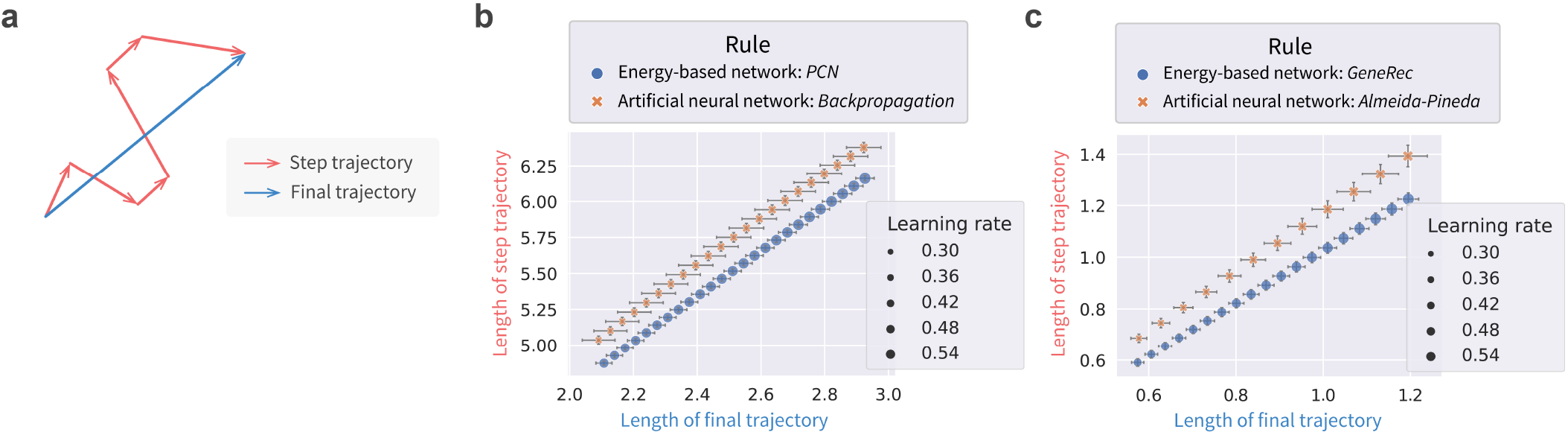
Prospective configuration produces less erratic weight modification. An experiment verifies that *energy-based networks* (EBNs) (i.e., prospective configuration), produce a less erratic weight modification than *artificial neural networks* (ANNs) (i.e., backpropagation). ▶ **a** | Experimental procedure. The weights are updated for a fixed number of steps on a fixed number of data points, which produces the step trajectory in the weight space (each red arrow corresponds to one weights update). Connecting the start and end points of the step trajectory (i.e., the initial and final weights of the model) produces the final trajectory (blue). A learning rule with less erratic weight modification would produce a shorter step trajectory relative to the final trajectory. This property of less erratic weight modification is also desirable for biological systems, because each weight modification costs metabolic energy. ▶ **b** | Comparison of the length of step and final trajectories between EBN, predictive coding network (PCN), and an ANN, trained with backpropagation. Note that the length of both trajectories depends on the learning rate. Thus, in panels b–c, we present the length of the step and final trajectory on y and x axis, respectively; each point is from a specific learning rate (represented by the size of the marker; the legend does not enumerate all sizes). In such plots, when the two learning rules produce roughly the same length of final trajectory (which could be from different learning rates), one can compare the length of their step trajectory. ▶ **c** | The same comparison as in panel c, but for another EBN, namely, *GeneRec*^105^. GeneRec describes learning in recurrent networks, and ANN with this architecture is not trained by standard backpropagation, but by a variant of backpropagation, called *Almeida-Pineda*^106–108^. **Implementation details**. We train the models to predict a target pattern from an input pattern (both randomly generated from 𝒩 (0, 1) and of 32 entries), and there are 32 pairs of them (32 datapoints). The structure of the networks is 32→ 32→ 32→ 32. The batch size is one, as biological systems update the weights after each experience. The training is conducted for 64 epochs (one epoch iterates over all 32 datapoints). At the end of each epoch, current weights of the network are recorded as one set. Thus, it results in a sequence of 64 sets of weights. Each set of weights is used as one point to construct the step trajectory. The first and last sets of weights are used to construct the final trajectory. The length of the step and final trajectories can then be computed and reported in Extended Data Figs. 7b–c. For each combination of learning rule and learning rate, simulation is repeated 20 times with different seeds, and the error bars represent standard error.

**Extended Data Fig. 8.**
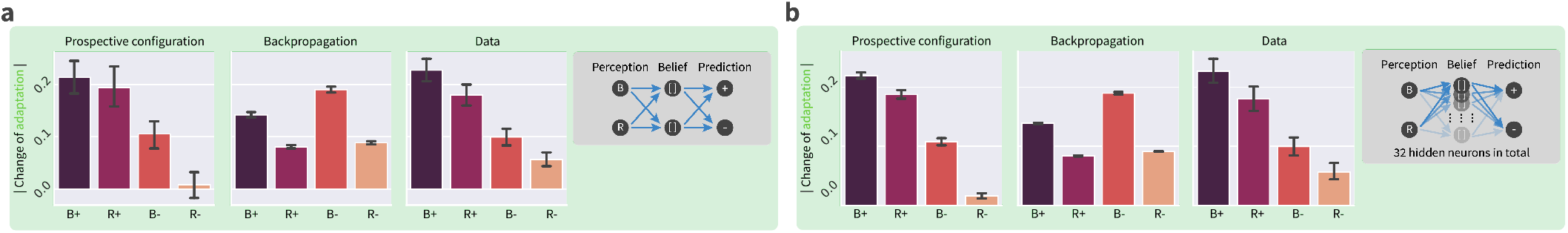
Motor learning experiment with fully-connected structure and more hidden neurons. In the experiments explaining biological observations, for simplicity, we simulated minimal networks necessary to perform these tasks, but it is important to establish if task structure can be discovered and learned by the networks without specifying network structure. Thus, here we repeat the motor learning experiment in Fig. 5 with general fully-connected structure (panel a) and 32 hidden neurons (panel b). Insets illustrate the structure of the networks. In both cases, prospective configuration is able to discover the task structure itself and reproduce the experimental observations; while backpropagation cannot.

**Extended Data Fig. 9.**
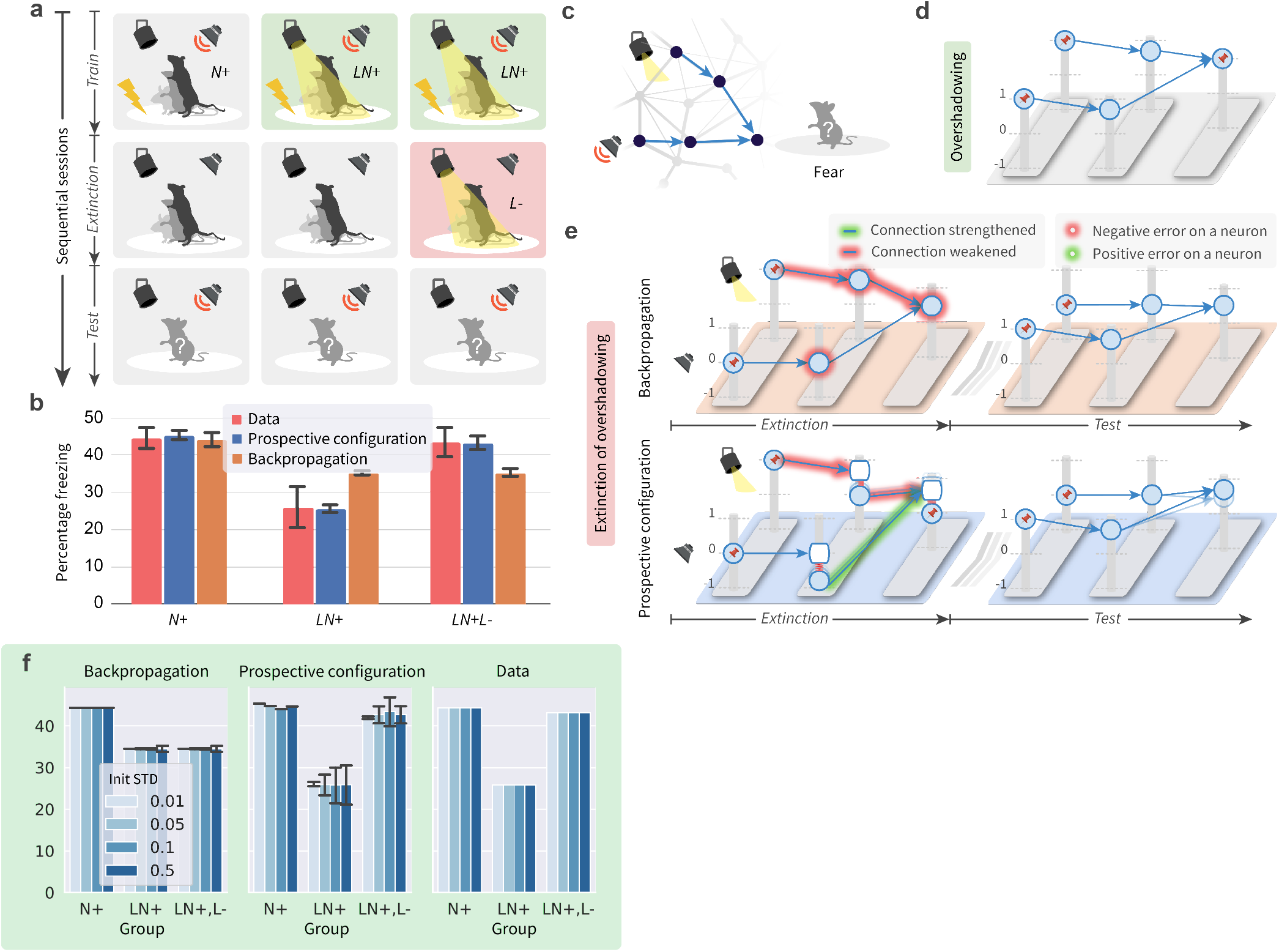
Prospective configuration explains extinction of overshadowing in fear conditioning (complete description of the experiment in Fig. 6). The extinction of overshadowing effect^77^ can be accurately reproduced and explained by prospective configuration, but not backpropagation (comparing “Data” against “Prospective configuration” and “Backpropagation” in panel b). ▶ **a** | Experimental procedure. Rats were divided into three groups, corresponding to three columns. Each group underwent three sessions sequentially, corresponding to the top three rows, namely, train, extinction, and test. The goal of the training session was to associate fear (+) with different presented stimuli *N* or *LN* depending on the group: rats experienced an electric shock paired with different stimuli, where *N* and *L* stands for noise and light, respectively. Next, during the extinction session no shock was given, and for the third group the light was presented but without the shock, aiming to eliminate the fear (-) of light (*L*). Finally, all groups underwent a test session measuring how much fear was associated with the noise: the noise was presented and the percentage of freezing of rats was measured. ▶ **b** | Experimental and simulation results. The bar chart plots the percentage of freezing during test for each group, both measured in the animal experiments^77^ (i.e., Data) and simulated by the two learning rules. Two effects are present in experimental data. First, comparing the groups *N*+ and *LN*+ demonstrates the overshadowing effect: there is less fear of noise if the noise had been compounded with light when paired with shock *LN*+ than if the noise alone had been paired with shock *N*+ (that is, light overshadows noise in a conditioned fear experiment). This effect can be accounted for by the canonical model of error-driven learning — the Rescorla-Wagner model^82^, and consequently it can be also produced by both error-driven models we consider — backpropagation and prospective configuration (explained in panel d). Second, comparing the groups *LN*+ and *LN*+*L*-shows the striking effect of extinction of overshadowing: presenting the light without the shock increases the fear response to the non-presented stimulus — noise. This effect is not produced by backpropagation, but can be reproduced by prospective configuration (explained in panel e). ▶ **c** | The neural architecture considered: both stimuli are processed by hidden neurons (i.e., intermediate neurons corresponding to visual and auditory cortices) and are then combined to produce the prediction of electric shock (i.e., fear). ▶ **d** | Explanation of overshadowing effect, i.e., the reduced percentage freezing comparing group *LN*+ against *N*+. With the energy machine introduced in Fig. 2, the diagram illustrates the state of the network after the *Train* sessions in groups *LN*+ and *LN*+*L*-. The network learns to predict a shock (i.e. produces output of 1), on the basis of two stimuli, hence each of the inputs to the output neuron must be 0.5. Therefore, if only one stimulus is presented, the output of the network is reduced to 0.5. The network shown in this panel is acting as the starting point of learning in panel e. ▶ **e** | Explanation of extinction of overshadowing effect, i.e., the increased percentage freezing after noise in group *LN*+*L*-in comparison to *LN*+. This effect suggests that during extinction trials, where light is presented without a shock, the animals increased fear prediction to noise. As shown in this panel, backpropagation (top) cannot explain this, since the error cannot be backpropagated to and drive a weight modification on a non-activated branch where no stimuli are presented; prospective configuration (bottom), however, can account for this. Specifically, on the non-activated branch, the hidden neural activity decreases from zero to a small negative value (it may correspond to a neural activity decreasing below the baseline^125^). Since a weight modification depends on the product of the presynaptic activity and the postsynaptic activity representing the error, which are both negative here, the weight on the non-activated branch is strengthened. ▶ **f** | Robustness to different standard deviations of initial weights. We also simulated networks with different standard deviations of initial weights (ranging from 0.01 to 0.5, represented by the depth of the colour). It is shown that prospective configuration fits better to the data measured in the animal experiments than backpropagation, regardless of the standard deviation of initial weights.

**Extended Data Fig. 10.**
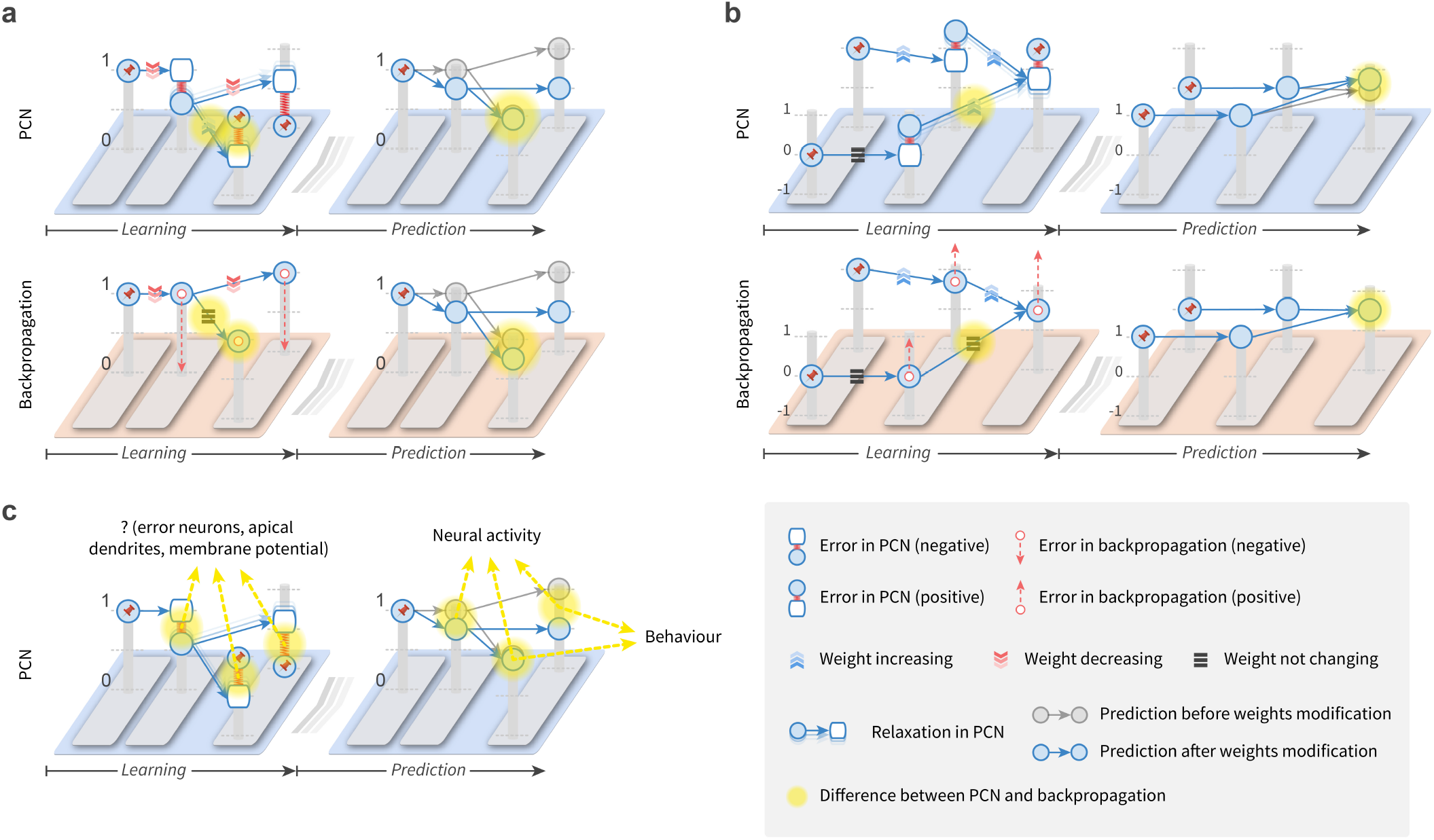
Experimental predictions of prospective configuration and backpropagation. To provide examples of experimental predictions of prospective configuration, panels a-b (and Extended Data Fig. 11) add the different behaviour of the learning rules in simple network motifs, which are minimal networks displaying given behaviour. Two motifs in this figure have been already analysed earlier in the paper, but there we focused on differences corresponding to experimentally observed effects, while in this figure we also add other qualitative differences that reveal a range of untested predictions of prospective configuration. Here, we consider a *predictive coding network*^25, 40, 52^ (PCN) with the energy machine in Fig. 2, however, a similar analysis can be applied to other energy-based networks, which also follow the principle of prospective configuration. In each panel, the top and bottom rows demonstrate the prediction of PCNs and backpropagation, respectively. The left column adds the differences in the prediction errors during learning and the resulting weight update. The right column demonstrates the neuron activity before (transparent) and after (opaque) weight update. The differences between the rules are added in yellow. Experimental predictions following from them can be derived as summarized in panel c. ▶ **a** | The error may spread to the branch where the prediction is correctly made. This motif has been compared with experimental data in Fig. 7, but here we focus on the effect illustrated in Fig. 1 and Fig. 2d, which despite being intuitive, has not been tested experimentally to our knowledge. The panel adds that an error on one output in PCN results in prediction error on the other, correctly predicted output. This produces an increase of the weight of the correct output neuron, which compensates for the decrease of the weight from the input, and enables the network to make correct prediction on the next trial. ▶ **b** | The error may cause a weight change in the sensory regions associated with absent stimuli. The panel shows a similar motif as the one investigated in Extended Data Fig. 9. The difference is that Extended Data Fig. 9 introduces negative error while this panel introduces positive error on the same architecture. Interestingly, introducing negative (Extended Data Fig. 9) or positive (this panel) error to the same architecture produce a similar effect in the PCN, i.e. an increased predicted output for the stimulus not presented during learning. ▶ **c** | Observing model behaviour in experiments. The diagram summarizes how the differences added in previous panels could be measured in experiments. The key difference in models’ behavior during learning is the difference in error signals. However, currently it is not clear how the prediction errors are represented in the cortical circuits. Three hypotheses have been proposed in the literature that errors are encoded in: activity of separate error neurons^40, 121, 126^, membrane potential of value neurons^127, 128^, membrane potential in apical dendrites of value neurons^22, 32^. Nevertheless, if the future research establishes how errors are encoded, it will be possible to test the predictions related to errors during learning. For example, one can design a task corresponding to panel a, where predictions in two modalities have to be made on the basis of a stimulus. One can then test if omission in one modality results in error signals in the brain region corresponding to the correctly predicted modality. The models also differ in the neural activity of the value nodes during the next trial following the learning. Such predictions are easier to test, because if the model makes a prediction without observing any supervised signal, then all errors are equal to 0 in PCNs, so the neural activity should reflect just the activity of value nodes. Additionally, the differences in the activity of the output value neurons should be testable in behavioural experiments. For example, panel b makes a behavioural prediction (presenting light with stronger shock should also increase freezing for tone) that can be tested in a similar way as described in Extended Data Fig. 9. Testing this prediction would also validate our explanation of the experimental result in Extended Data Fig. 9.

**Extended Data Fig. 11.**
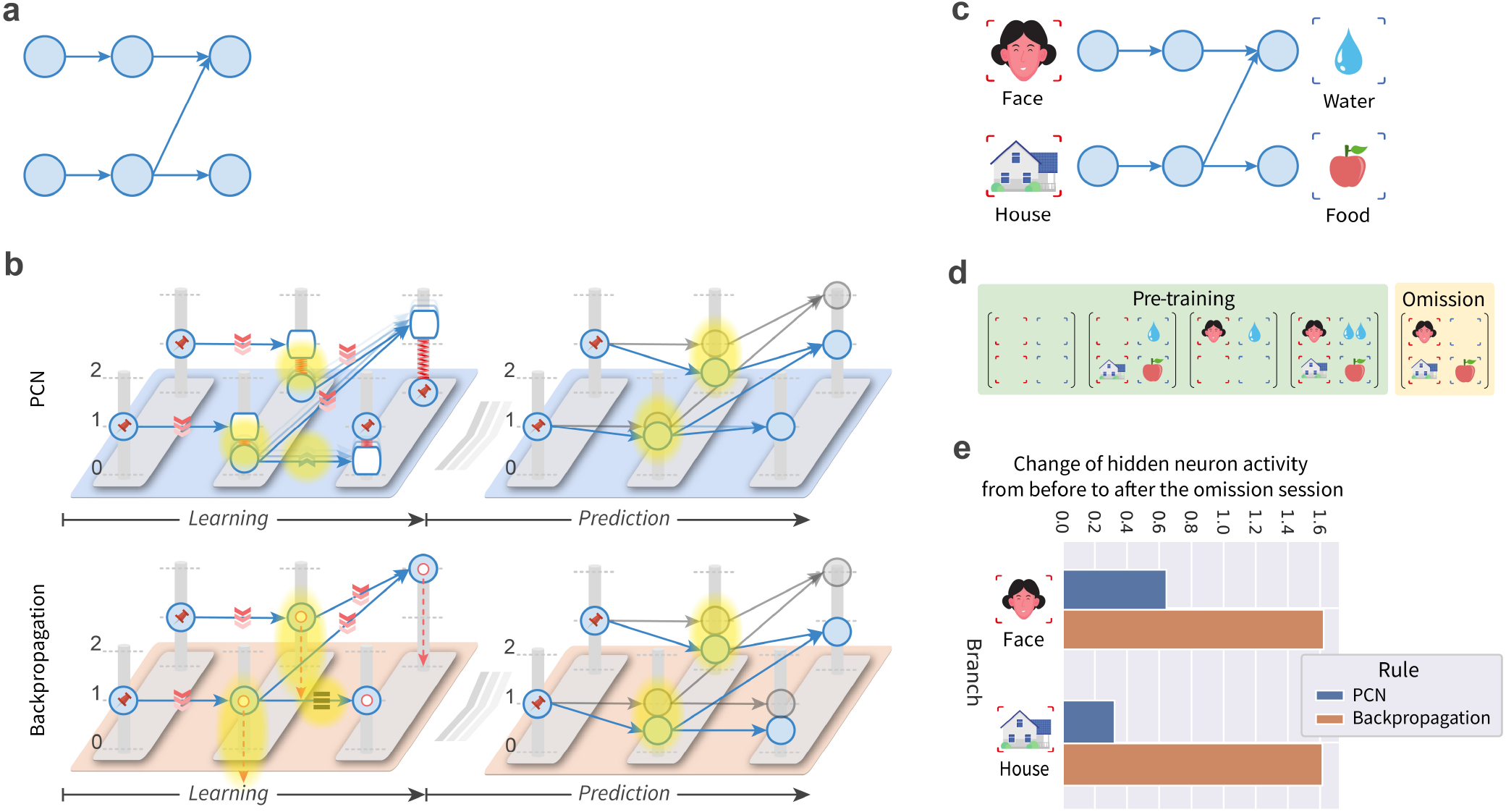
Experimental predictions concerning errors assigned to hidden nodes. The figure demonstrates a striking difference in how prospective configuration and backpropagation assign error to hidden nodes. Namely, in prospective configuration, the error assigned to a hidden node is reduced if the node is also connected to correctly predicted outputs. This difference is illustrated in a motif (panel a), for which we illustrate behaviour of learning rules with the energy machine (panel b), and describe a sample experiment testing model predictions (panels c–d). Finally, we report the simulation results of the two learning rules (panel e), confirming that they indeed make distinct predictions for this motif. ▶ **a** | In this motif, two stimuli are presented and two predictions are made. One stimulus contributes to only one prediction, while the other stimulus contributes to both predictions. ▶ **b** | Comparison of learning rules’ behaviour with the energy machine (notation as in Extended Data Fig. 10). The diagrams illustrate a network containing the motif (panel a), in a situations where one of the predicted outputs (top output) is omitted. A negative error is introduced to the prediction determined by both stimuli. Thus, we would expect the error to be assigned to hidden neurons on both branches. Both learning rules do so, however, they assign errors differently. PCNs allocate less error on the bottom hidden neuron than the top hidden neuron, because the bottom hidden neuron also contributes to another output that was correctly predicted, while backpropagation assigns the same error to both hidden neurons. This is also a nice example where prospective configuration (PCNs) demonstrates more intelligent behavior. ▶ **c** | Experimental stimuli. To test this motif, it is important to choose stimuli for which neural activity of “hidden” neurons can be easily measured. In case of a human experiment, inputs could contain faces and houses, because the hidden neurons would correspond to the brain regions known to be specifically excited by these particular types of stimuli, and the activity of these regions could be easily distinguished in an experiment^129^. The outputs could correspond to reward modalities (e.g. water and food). In case of a human experiment, these could be “virtual rewards” the participants are instructed to gather, while for animals, these could be the actual rewards. ▶ **d** | Experimental procedure. The motif shown in panel c could arise in brain networks from training with examples shown in the green box. To test differences in behaviour of learning rules, partial omission trials could be presented, in which one of the expected outputs is omitted, as shown in the orange box. ▶ **e** | Results of simulations. We pre-train the models with the examples in the green box in panel d for a sufficient number of iterations until convergence, and then we train the model with the omission using the example in the orange box in panel d for one trial. We measure the change of hidden neural activity on both branches from before to after the above omission session. The graph shows simulation results of such change in hidden activity: PCNs predict different changes on different branches, while backpropagation predicts the same change on different branches (consistent with illustration in panel b, right). **Implementation details**. Presenting and not presenting a stimulus (face, house, water, or food) are encoded as 1 and 0, respectively. Presenting two drops of water is encoded as 2. The network is initialized to the pre-trained connection pattern demonstrated in Extended Data Figs. 11c, i.e., the weights visible on the panel are set to one and other weights are set to zero. Such pattern of weights would arise from pre-training with the four examples in Extended Data Figs. 11d (in the green “Pre-training” box), but for simplicity, we do not simulate such pre-training but just set the weights as explained before. Next, to measure the activity of hidden units of such network during prediction, we set both inputs to 1 and record the hidden neural activity of the two branches. Subsequently, the model is presented with the omission trial shown in the orange box and the weights are updated once. Finally, to measure weight changes resulting from training on the subsequent prediction trial, we set both inputs to 1 and record the hidden neural activity of the two branches for the second time. The change of the hidden neuron activity from before to after the omission session can thus be computed for both branches.

## 2 Supplementary Information

In this supplement, we present additional description and analysis of the simulated models. In Section 2.1, we provide details of all models simulated in the paper. In Section 2.2, we discuss relationship between prospective configuration and target propagation. In Section 2.3, we analyse prospective index of PCNs. In Section 2.4, we analyse target alignment of various learning models.

### 2.1 Details of simulated models

This section gives more details of all simulated models. The general idea of *energy-based networks* (EBNs) and *artificial neural networks* (ANNs), and one of EBNs, *predictive coding network*^25, 40, 52^ (PCN), have been described in the Main text and Methods. PCN is again included here along with other simulated models to provide descriptions in a unified form, facilitating the reproduction of our reported results. Complete code and full documentation reproducing all simulation results will be made publicly available at https://github.com/YuhangSong/A-New-Perspective upon publication of this work.

Algorithms 3 to 7 describe how the four models simulated in this paper predict and learn. These four models are: PCN, backpropagation, *GeneRec*^105^, and *Almeida-Pineda*^106–108^. Among the four models, PCN and GeneRec are the two EBNs we investigate; backpropagation and Almeida-Pineda are the two ANNs we investigate. Specifically, PCN is compared against backpropagation, because it has been established that PCN are closely related to backpropagation^25, 33^ and they make the same prediction with the same weights and input pattern^25^. Therefore we simulated prediction in these two algorithms in the same way (Algorithm 3). However, they learn differently (c.f. Algorithms 4 and 1). The other EBN, GeneRec, describes learning in recurrent networks, and ANN in this architecture is not trained by standard backpropagation, but a modified version proposed by Almeida and Pineda^106–108^ (thus called the *Almeida-Pineda* algorithm). Thus, GeneRec should be compared against Almeida-Pineda because they make same prediction with the same weights and input pattern^105^. Therefore we simulated prediction in these two algorithms in the same way (Algorithm 5). But they learn differently (c.f. Algorithms 6 and 7). In a word, PCN and backpropagation are EBN and ANN working in feed-forward architecture, respectively; GeneRec and Almeida-Pineda are EBN and ANN working in recurrent architecture, respectively.

#### Algorithm 3: Predict with backpropagation or *predictive coding network*^25, 40, 52^ (PCN)

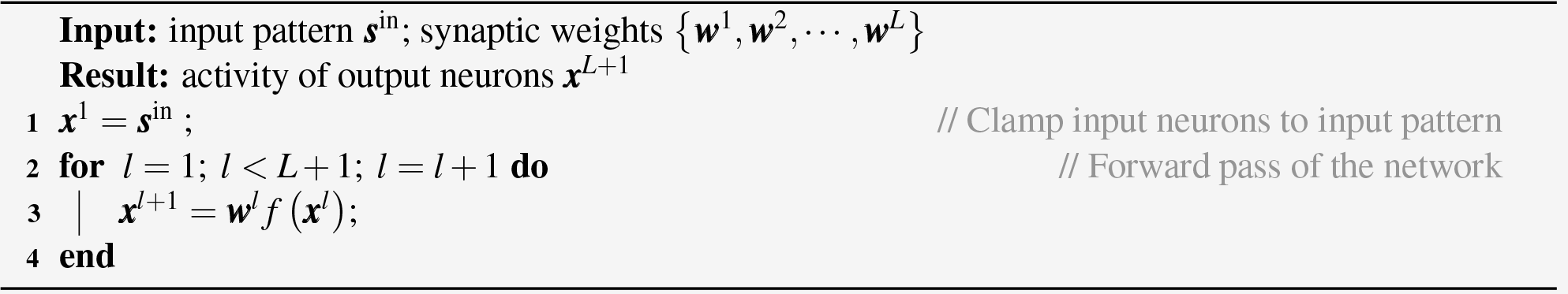

Particularly, PCN & Backpropagation work in a network where prediction is made from the input through a series of forward weights {***w***^1^, ***w***^2^, …, ***w***^*L*^ }; GeneRec & Almeida-Pineda works in a network where prediction is made from input through a mixture of forward weights {***w***^1^, ***w***^2^, …, ***w***^*L*^} and backward weights {***m***^1^, ***m***^2^, …, ***m***^*L*^}. The forward weights {***w***^1^, ***w***^2^, …, ***w***^*L*^} and backward weights {***m***^1^, ***m***^2^, …, ***m***^*L*^} are not necessarily related. This architecture is also similar to the continuous Hopfield model^130, 131^. Unlike in some previous studies^24^, here, we focus on layered networks, where the sets of neurons at adjacent layers ***x*** and ***x*** are connected by synaptic weights. Thus, we define two sets of weights for GeneRec & Almeida-Pineda that works in the recurrent network: ***w***^*l*^ is the forward weights connecting from ***x***^*l*^ to ***x***^*l*+1^; ***m***^*l*^ is the backward weights connecting from ***x***^*l*+1^ to ***x***^*l*^.

#### Algorithm 4: Learn with backpropagation

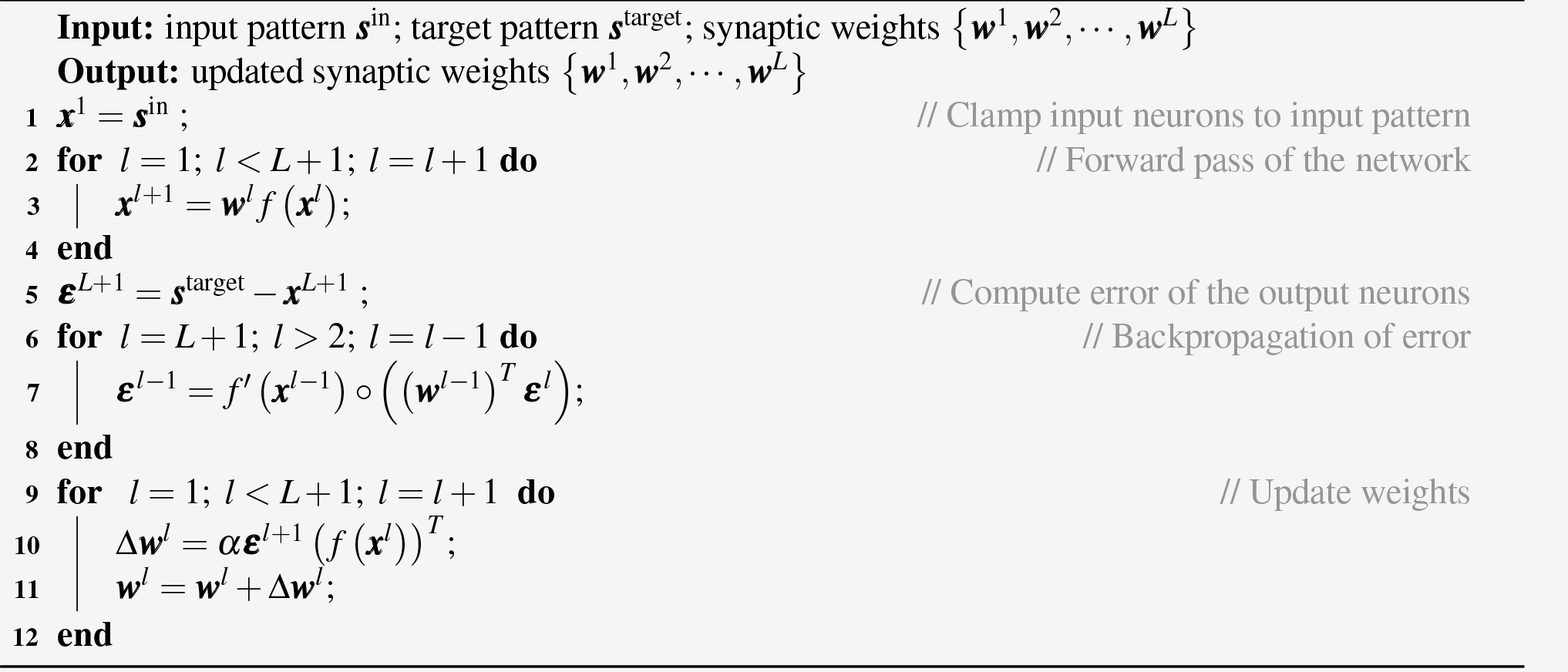

Also note that GeneRec has been explored and re-discovered in recent works^48, 132^ showing how a closely related algorithm resembles backpropagation when the backward weights are the transposes of the forward weights ***m***^*l*^ = (***w***^*l*^)^*T*^ (or for a fully-connected network in their context *w*_*i, j*_ = *w*_*j,i*_), and how the extreme version of the algorithm approximate backpropagation^24^.

Extended Data Fig. 5 additionally investigates *Strong Deep Feedback Control*^92, 93^ (strong-DFC). *Deep Feedback Control*^124^ (DFC) was proposed to work with “infinitely weak nudging”, as in equilibrium propagation^24^. More recent work demonstrates that it also works with “strong control”^92, 93^ (thus, called strong-DFC), i.e., with the natural form of EBNs. Thus, in this paper we investigate strong-DFC. In strong-DFC (or DFC in general), backward weights ***m***^*l*^ do not connect from layer *l* + 1 to layer *l* as in other models investigated in the paper. Instead, ***m***^*l*^ connects from the output layer *L* + 1 to layer *l*. We use the provided code in https://github.com/mariacer/strong_dfc to simulate strong-DFC. All hyper parameters are kept as is in the provided code. We remove the activation function of the last layer in the original implementation^124^, to keep consistent with the rest of the models investigated in this paper, thus, provides a fair comparison. Derivation and motivation of the model can be found in the original paper^92, 93^.

Some common notations in the algorithms are: *α* is the learning rate for weights update; *γ* and 𝒯 are the integration step and length of relaxation, respectively (specified to the two EBNs, PCN and GeneRec); ***s***^in^ and ***s***^target^ are the input and target patterns, respectively. For Almeida-Pineda, which requires additional iterative process to propagate error, *β* and 𝒦 are the integration step and length of this iterative process, respectively. In our simulation, we use *β* = 0.01 and 𝒦 = 1600.

All simulated models work in mini-batch mode, that is to say, one iteration is to update the weights for one step on a mini-batch of data randomly sampled from the training set for classification tasks. The above sampling is without replacement, i.e., the same examples will not be sampled again before the completion of a epoch, which is when the entire training set has been sampled once. For example, considering a dataset of 1000 examples with a batch-size (number of examples in a mini-batch) of 10, then each iteration would update weights for one step on 10 examples, and it will take 100 such iterations to complete one epoch. To implement the Algorithms 3 to 7 described below in mini-batch mode, one can simply add an extra-dimension, the size of which is batch-size, to all the neuron-specific vectors in the algorithms such as ***x***^*l*^, ***ε***^*l*^ and etc., and then reduce this dimension by summing over it when computing weight update Δ***w***^*l*^ (and Δ***m***^*l*^ if the model is GeneRec or Almeida-Pineda).

Note that learning with Almeida-Pineda involves relaxation of the model, i.e., updating neural activity, in lines 5-12 of Algorithm 6. However, its function is to make a prediction with current weights and input pattern so that the error on the output neurons can be computed (in the following line 13), similar as the function of “forward pass” in backpropagation in lines 2-4 of Algorithm 4. The neural activity in the Almeida-Pineda model is fixed during spreading of error, like in backpropagation. Thus, Almeida-Pineda is classified as an ANN rather than an EBN (which updates neural activity during spreading of error).

#### Algorithm 5: Predict with *Almeida-Pineda*^106–108^ or *GeneRec*^105^

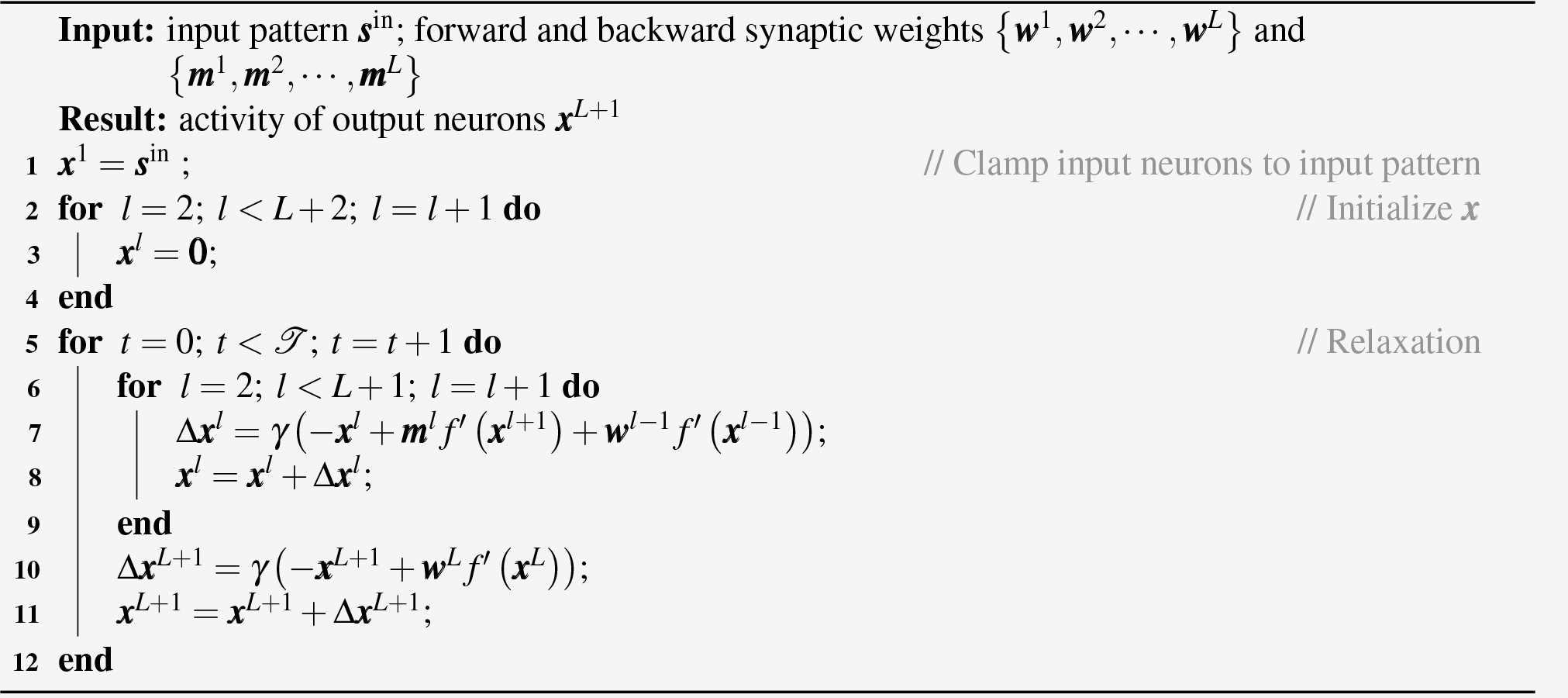

#### Algorithm 6: Learn with *Almeida-Pineda*^106–108^

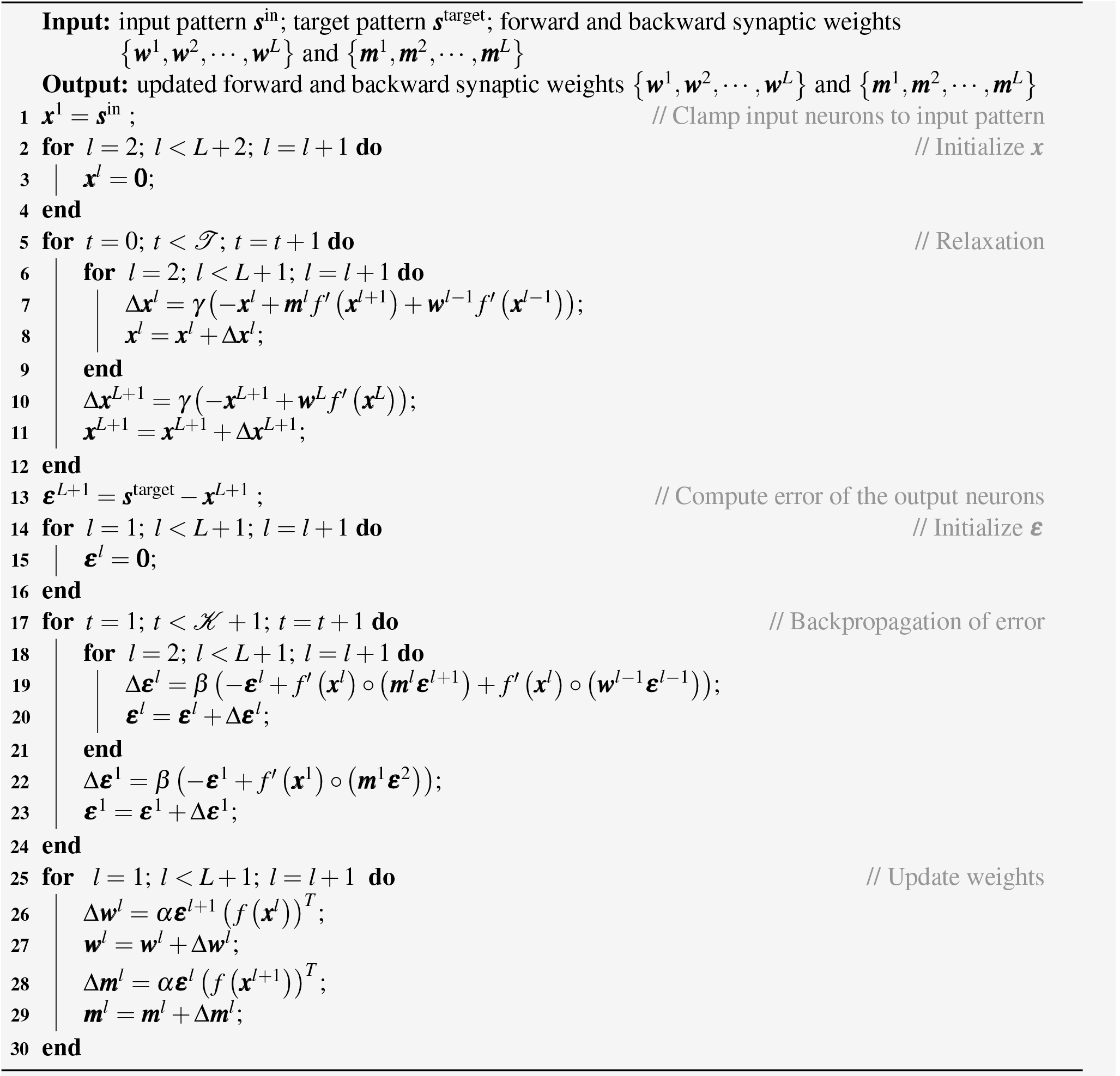

#### Algorithm 7: Learn with *GeneRec*^105^

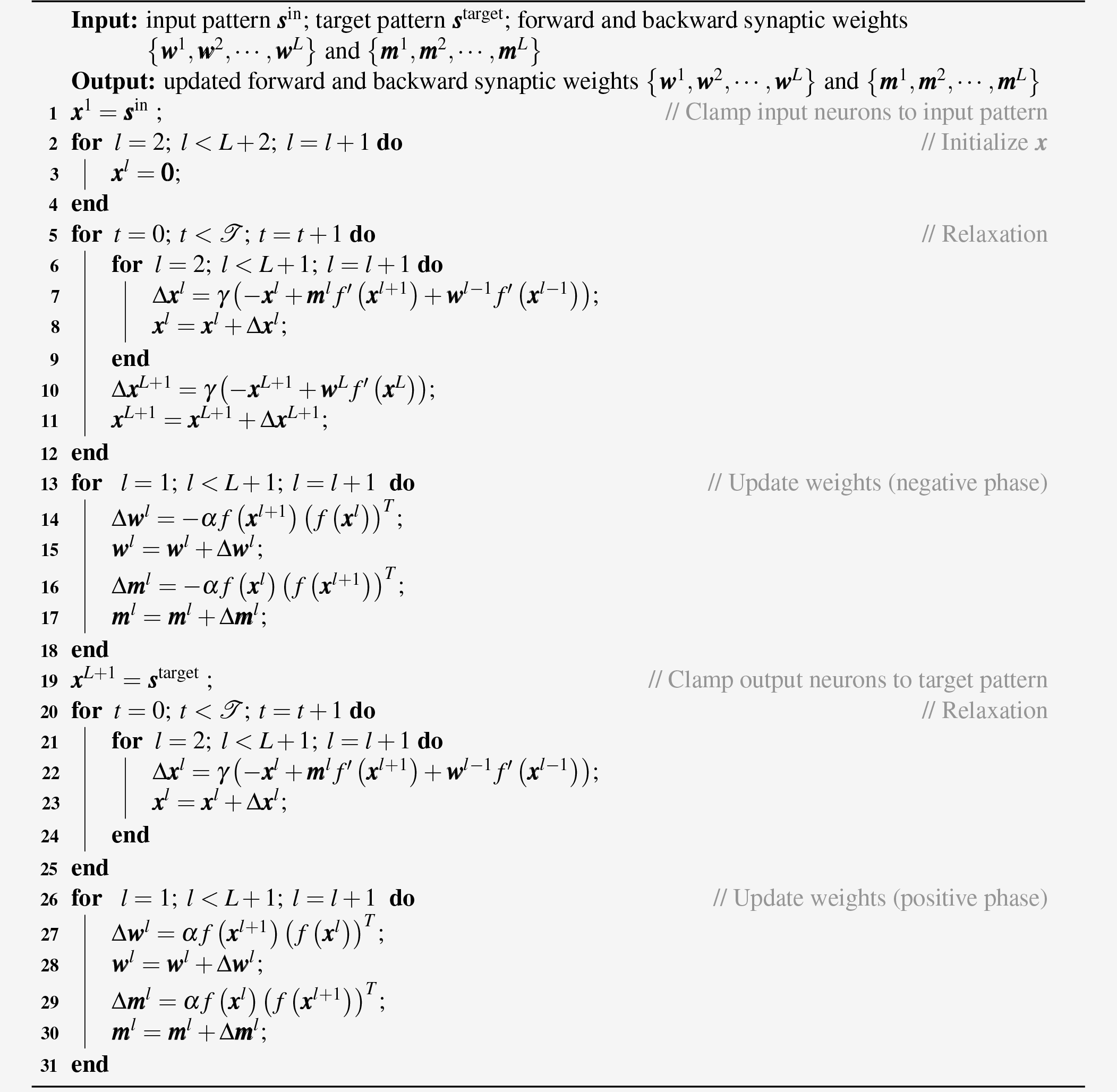

### 2.2 Relationships of predictive coding networks to target propagation (Extended Data Figs. 3)

In Extended Data Figs. 3, we illustrate that prospective configuration, particularly, *predictive coding network*^25, 40, 52^ (PCN), has close a relationship to target propagation^57^. In this section, we formally prove these observations.

Note that these relationships of predictive coding networks to target propagation on one hand build interesting connections to existing work, on the other hand serve as a step in providing a mathematical explanation of the target alignment of predictive coding networks, as discussed in the later Section 2.4.4.

#### 2.2.1 Target propagation

##### Algorithm 8: Learn with target-propagation

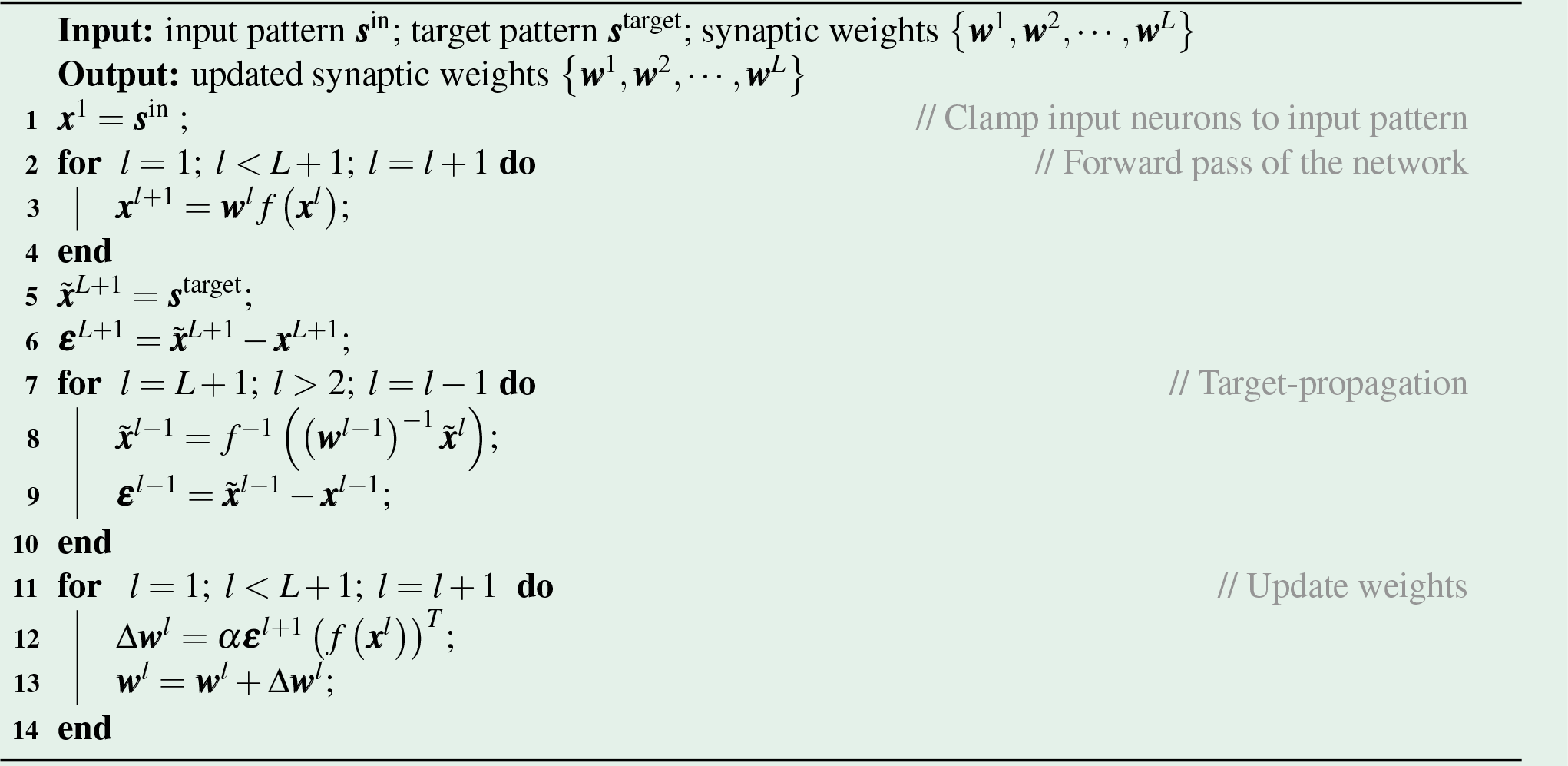

We first briefly review target propagation. The key insight behind target propagation is that rather than updating weights based on a gradient of a loss function, one can instead attempt to explicitly compute what are the optimal activity for the neurons so that they can produce the desired target pattern, and then update the weights so as to nudge the current neural activity towards the optimal activity directly. We call these optimal activity *local target* since if the neurons takes this activity, the network would produce the desired target pattern. Importantly, we can directly compute the local target in terms of the *inverses* of the weights and activation functions. Namely, suppose that we have a three-layer network with activation functions *f* (), weight matrices ***w***^1^, ***w***^2^, ***w***^3^ and an input pattern ***s***^in^. The output of this network is ***x***^4^ = ***w***^3^ *f* (***w***^2^ *f* (***w***^1^ *f* (***s***^in^))). Suppose instead that we do not want the network to output ***x***^4^ for a given ***s***^in^ but rather a given target pattern ***s***^target^. Then, the activity at the first layer 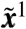 that would produce this desired activity can be exactly computed by inverting^1^ the network 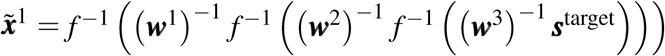. From this, we can define a recursion of one local target in terms of another at the layer above,

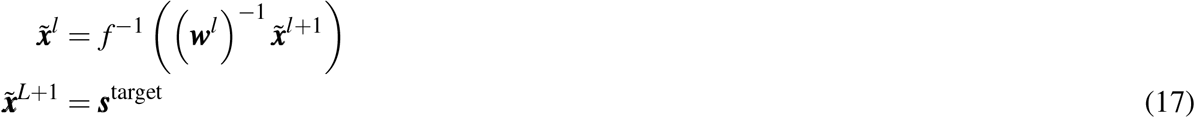

Based on these targets we can define the errors in target propagation as 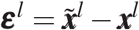. These errors drive the update of weights according to:

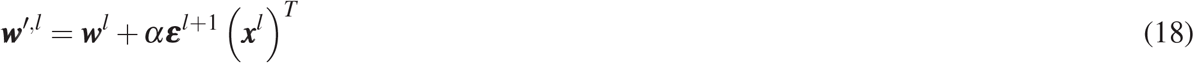

This algorithm is summarized in Algorithm 8.

#### 2.2.2 Analyses of the relationships

Now we formally prove the below observations in Extended Data Figs. 3 about how prospective configuration, particularly, *predictive coding network*^25, 40, 52^ (PCN), has close a relationship to target propagation^122^. In other words, we formally prove that

- In an output-constrained PCN, neural activity after relaxation converges to the local target;
- In an input-output-constrained PCN, neural activity after relaxation approaches to the weighted sum of the predicting activity and the local target.

In the above, predicting activity refer to the neural activity when the model is making prediction, and they are the same for both backpropagation and PCN as they compute the same neural activity when making a prediction.

##### Output-constrained PCN

As mentioned, we first investigate the “output-constrained PCN”: in this PCN input neurons are not clamped to the input pattern but output neurons are clamped to the target pattern. We show that in this PCN, the activity after relaxation is precisely equal to the local target. Since ***x***^1^ is not constrained to the input pattern, we can look at its dynamic by setting *l* = 1 in Eq. (12). Since there is no error term or error nodes at the input layer, there is only the later term left when setting *l* = 1 in Eq. (12) (note that here we write in matrix & vector form):

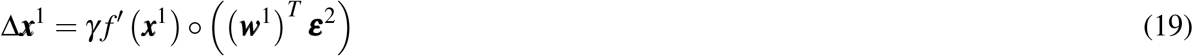

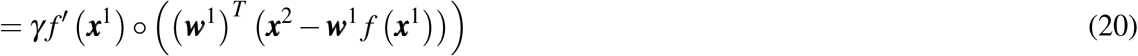

Considering the above dynamic has converged, we can set Δ***x***^1^ = **0** in the above equation and solving for ***x***^1^, then we can obtain the converged value of ***x***^1^:

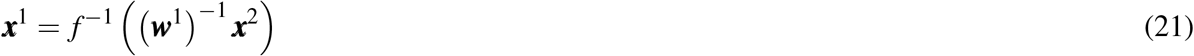

Now we look at the dynamic of ***x***^2^ by setting *l* = 2 in Eq. (12):

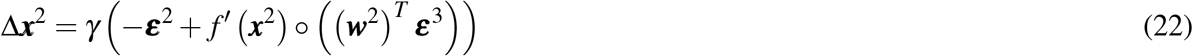

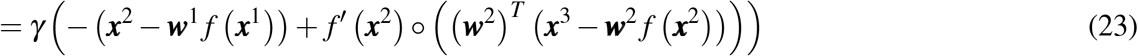

Putting the solved ***x***^1^, i.e., Eq. (21), into the above Eq., we have:

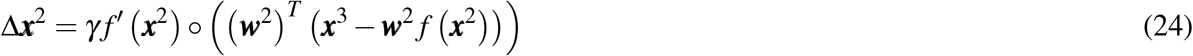

Considering the above dynamic has converged, we can set Δ***x***^2^ = **0** in the above equation and solving for ***x***^2^, then we can obtain the converged value of ***x***^2^:

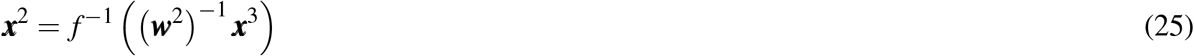

One can now see the proof goes recursively until *l* = *L* and ***x***^*L*+1^ is fixed to the target pattern ***s***^target^:

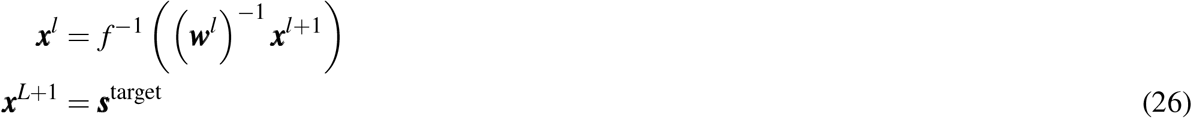

which is exactly the recursive formula of the local target in target propagation, i.e., Eq. (17). Thus, neural activity of output-constrained PCN after relaxation equals to the local target.

##### Input-output-constrained PCN

Secondly, we investigate the “input-output-constrained PCN”: in this PCN both input and output neurons are clamped to the input and target patterns, respectively. We show that in this PCN, the activity after relaxation are the weighted sum of the predicting activity and the local target. Particularly, since in a input-output-constrained PCN, we can only solve for the equilibrium after relaxation analytically in the linear case, we prove this for a linear PCN. Nevertheless, the analysis still provides useful insights. Looking at the network dynamics at a given layer *l*, i.e., Eq. (12), we can write the dynamics in the linear case as,

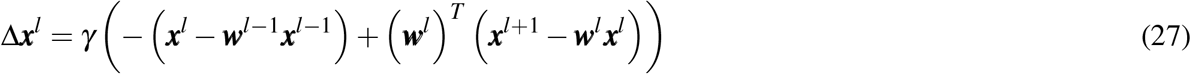

If we then set Δ***x***^*l*^ = **0** and solve for ***x***^*l*^, we obtain,

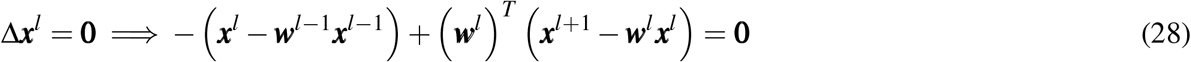

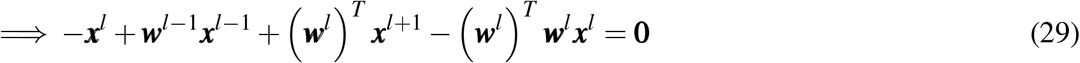

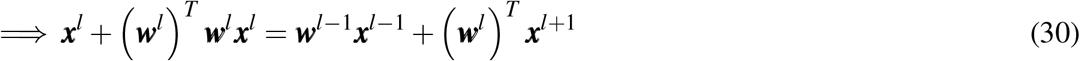

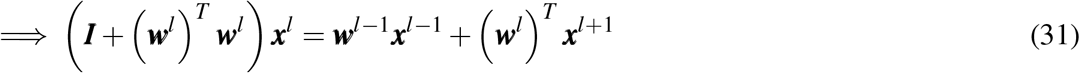

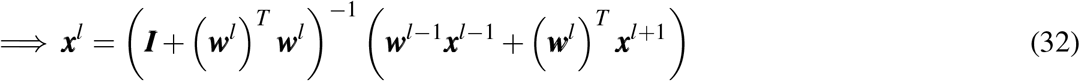

If we assume that the norm of the weights is large compared to the identity matrix ***I***, i.e., we consider (***I*** + (***w***^*l*^)^*T*^ ***w***^*l*^)^−1^ ≈ ((***w***^*l*^)^*T*^ ***w***^*l*^)^−1^, the above equilibrium solution can further be approximated by:

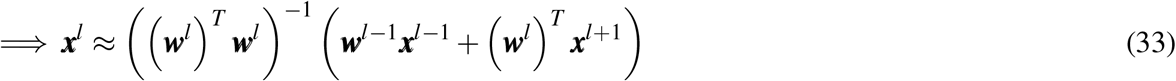

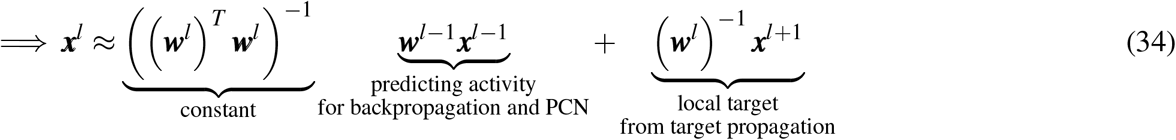

where the equilibrium solution is simply the weighted sum of the predicting activity and the local target.

In summary, during relaxation the activity in predictive coding networks tends to move from the predicting activity towards the local target that would be computed by target propagation. These relationships on one hand build interesting connections to existing work, on the other hand serve as a step in providing a mathematical explanation of the target alignment of predictive coding networks, as discussed in the later Section 2.4.4.

### 2.3 Prospective index of predictive coding networks (Extended Data Figs. 5)

This section formally proves two properties of the prospective index *ϕ*^*l*^ of a *predictive coding network*^25, 40, 52^ (PCN), that can be observed in Extended Data Figs. 5d. To briefly recap, prospective index *ϕ*^*l*^ quantifies to what extent the hidden neural activity of the network following clamping output neurons to a target pattern is shifting toward the hidden neural activity following subsequent weight modification. Below we show two properties visible in Extended Data Figs. 5d:

- Firstly, prospective index of the first hidden layer (*ϕ* ^2^) in a PCN is always one.
- Secondly, the prospective index in other layer is close to one because, the weights ***W*** in PCN are updated towards a configuration ***W*** ^*^ whose prospective index is one.

Note that these observations of high prospective index of predictive coding networks on one hand formally defines what we proposed as “prospective configuration” and distinguishes itself from backpropagation, on the other hand serve as a step in providing a mathematical explanation of the target alignment of predictive coding networks, as discussed in the later Section 2.4.4.

#### 2.3.1 Prospective index of the first hidden layer of PCN is always one

We assume that the model does not make a perfect prediction with the current weights, so that the error in the prediction drives the learning. As defined in Extended Data Figs. 5a, vectors ***ν***^⊕,*l*^ and ***ν***^′,*l*^ describe the changes in hidden neuron activity, due to target pattern being provided and learning respectively. Specifically for layer *l* = 2, these vectors are:

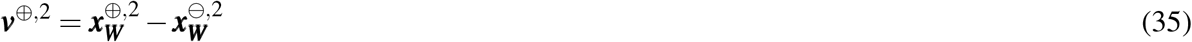

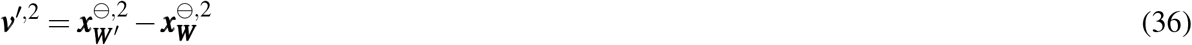

We will now show that for PCN the above vectors ***ν***^⊕,2^ and ***ν***^′,2^ point in the same direction. The change in activity due to learning ***ν***^′,2^ is equal to

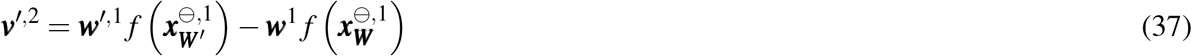

Since the value nodes of the first (input) layer ***x***^1^ are always fixed to the input signal ***s***^in^, the above Eq. (37) can further be written as,

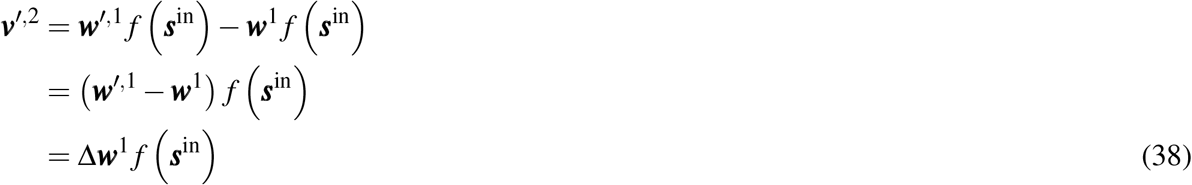

Using Eqs. (13) and (11), we write

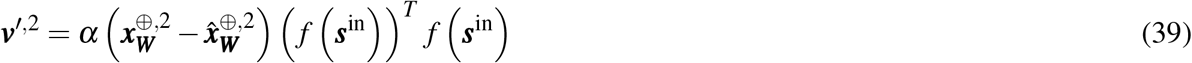

In Eq. (39), 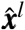 denotes inputs to neurons in layer *l*, i.e., 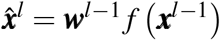. Note that 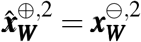, because both of these quantities are equal to ***w***^1^ *f*(***s***^in^) (the input of the first hidden layer (*l* = 2) does not change in response to output neuron being clamped). Using 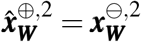, the above Eq. (39) can further be written as,

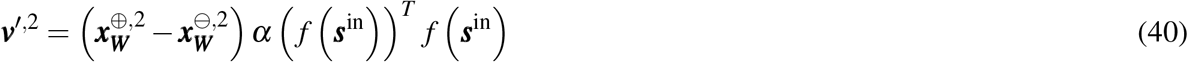

Note that *α* (*f* (***s***^in^)) ^*T*^ *f* (***s***^in^) is a positive scalar (if at least one entry in the input pattern is non-zero). Comparing Eqs. (35) and (40), we can see that vectors ***ν***^′,2^ and ***ν***^⊕,2^ are just scaled versions of each other, hence the cos of the angle between them is equal to 1, and thus prospective index is also equal to 1 (in the limit of *κ* → 0).

#### 2.3.2 Weights in PCN are updated towards a configuration with prospective index of one

As seen in Extended Data Fig. 5d, the prospective index for layers *l >* 2 is very close to one. To provide an intuition for why this is the case, in this section we demonstrate how PCNs would need to be modified to have prospective index equal to 1. We will refer to such modified model as target-PCN, and calculate its prospective index.

As in the previous section, we assume that the model does not make a perfect prediction with the current weights, so that the error in the prediction drives the learning. We start with recapping what happens in sequence in one iteration of the standard PCN.

1. Start from relaxation with only input neurons clamped to input pattern (⊖) and with current weight ***W***, the hidden neuron activity settles to: 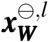
2. Both input and output neurons are clamped to the input and target pattern respectively (⊕) and then the hidden neuron activity is relaxed to: 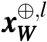
3. Weights ***W*** are updated for one step to ***W*** ^′^ to decrease the energy, while hidden neuron activity stays still from the last step: 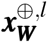
4. Output neurons are freed but the input neuron is still clamped to the input pattern and then the hidden neuron activity is relaxed to: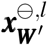

In the above step 3, weights are updated for one step from ***W*** to ***W*** ^′^. However, one can investigate the case of updating weights ***W*** for many steps until convergence ***W*** ^*^ in the above step 3. This will result in weights ***W*** ^*^ that represents: “the target towards which the weights ***W*** are updated”. Thus, we call this variant “target-PCN” and it is summarized in Algorithm 9. Specifically, the procedure of target-PCN is to replace the above steps 3 and 4 of standard PCN with:

3. Weights are updated for many steps from ***W*** to ***W*** ^*^ to decrease the energy till convergence, while hidden neuron activity stays still from the last step: 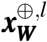;
4. Output neurons are freed but the input neuron is still clamped to the input pattern and then the hidden neuron activity is relaxed to: 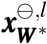;

##### Algorithm 9: Learn with target-PCN

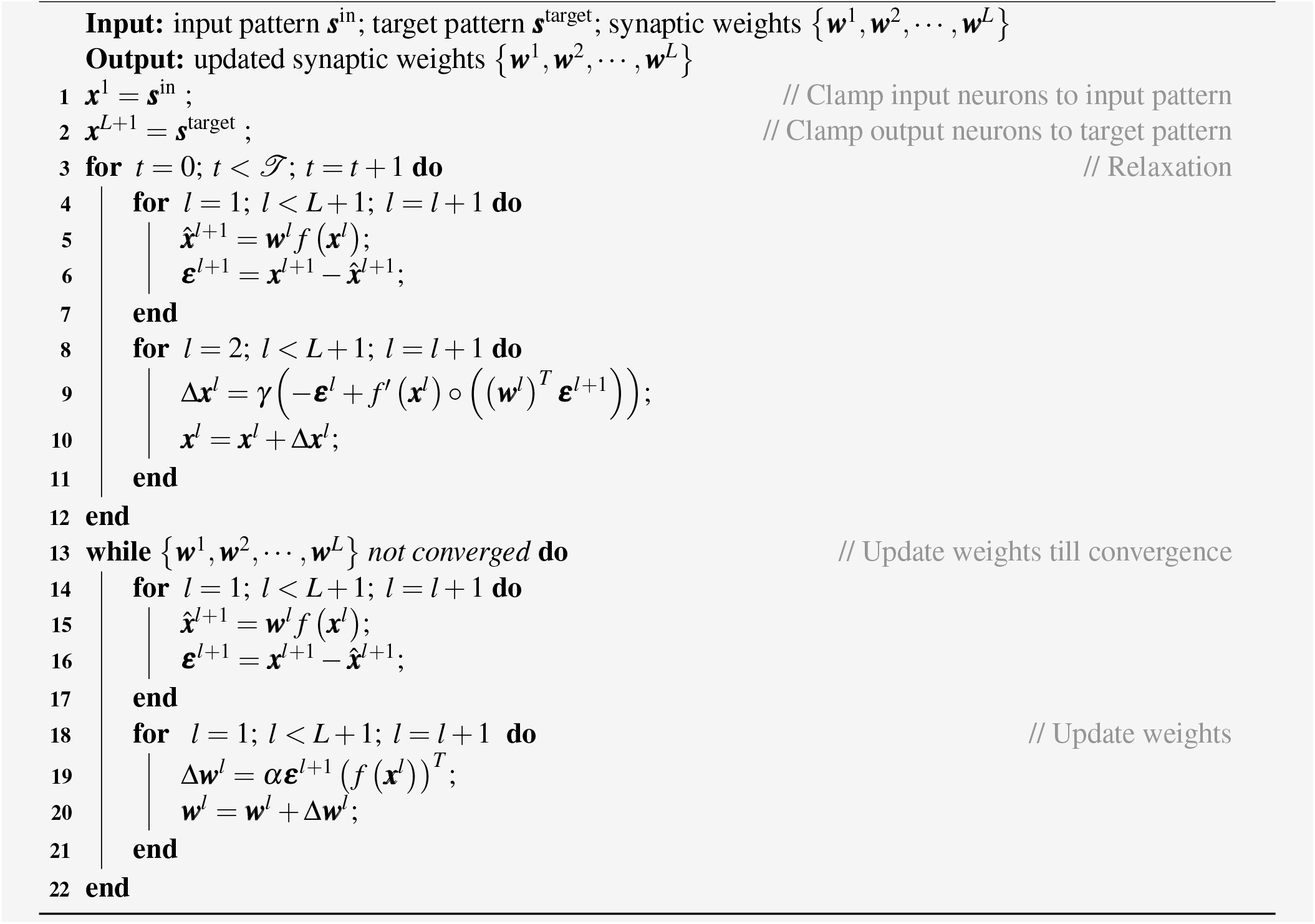

In the following, we demonstrate prospective index of target-PCN is one for all layers. First, we should notice that the minimum of energy *E* of PCN is zero, since the energy function is a sum of quadratic terms, i.e., Eq. (6). Then, we should notice that such energy *E* of PCN can be optimized to its minimum of zero by optimizing only ***W***. Particularly, the local energy term of layer *l* is:

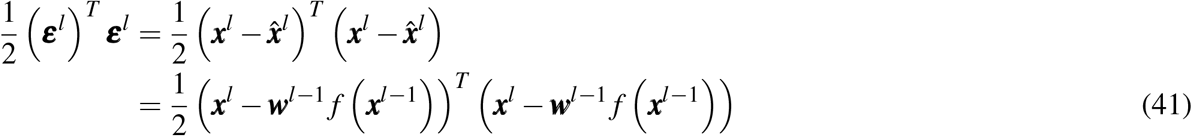

In the above Eq., ***x***^*l*^ − ***w***^*l*−1^ *f* (***x***^*l*−1^) can be optimized to produce a zero vector by optimizing only ***w***^*l*−1^, as long as *f* (***x***^*l*−1^) is not a zero vector. Specifically, let us denote all the non-zero entries in *f* ***x***^*l*−1^ by 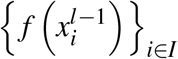, where *I* is the set of indices *i* so that 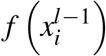 is non-zero. Since *f* (***x***^*l*−1^) is not a zero vector, *I* ≠ ∅. To demonstrate that there exists a solution for 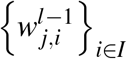 so that 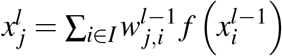 we construct an example of such solution. Such sample solution is to pick one index *g* from *I*, then have 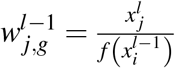 and 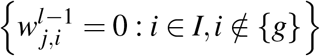. Thus, as long as *f* (***x*** ^*l*−1^) is not a zero vector (*I* ≠ ∅), there exists a solution of ***w***^*l*−1^ that makes ***x***^*l*^ − ***w***^*l*−1^ *f*(***x***^*l*−1^) a zero vector.

Thus, in step 3 of the target-PCN, the energy of the network is at its minimum of zero. This further implies that in the step 4 of the target-PCN, the neural activity does not move, i.e.,

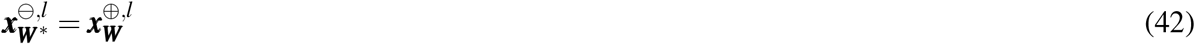

According to the definition of prospective index in Extended Data Figs. 5a-b, the prospective index of this target-PCN (*ϕ* ^*,*l*^) is:

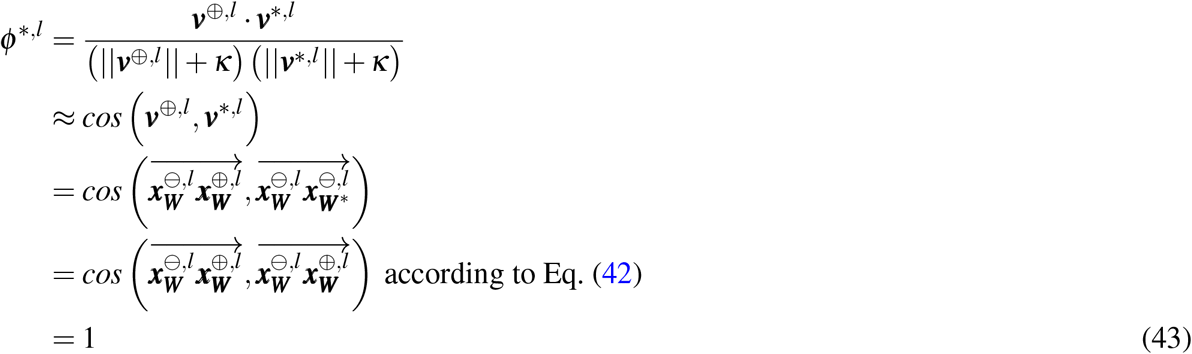

This theoretical result is further confirmed by empirical observation in Extended Data Figs. 5d. Since the standard PCN modifies the weights in a similar direction as target-PCN, it is likely to have a similar prospective index.

In summary, predictive coding networks has a high prospective index. This on one hand formally defines what we proposed as “prospective configuration” and distinguishes itself from backpropagation, on the other hand serve as a step in providing a mathematical explanation of the target alignment of predictive coding networks, as discussed in the later Section 2.4.4.

### 2.4 Target alignment

In this section we provide a mathematical analysis of target alignment. First, we show that the target alignment is equal to 1 for various networks that do not include hidden layers. Next we demonstrate that target propagation produces target alignment of 1. The third subsections identifies a special condition under which backpropagation produces target alignment of 1. The last subsection addresses the question of why predictive coding networks have higher target alignment than backpropagation, using several findings in earlier sections.

#### 2.4.1 Target alignment for networks without hidden layers (Fig. 3e)

Fig. 3e shows that target alignment for models without hidden layers, trained either with PC or BP, is exactly one, and here we prove this property analytically. Without hidden layers, PC and BP are identical algorithms. In a linear network, the change of the weight ***w***^1^ is:

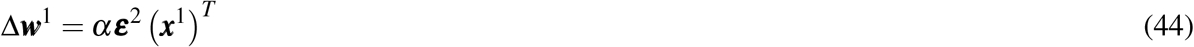

We denote output after learning by ***x***^′2^. The change of the output ***x***^′2^ − ***x***^2^ is:

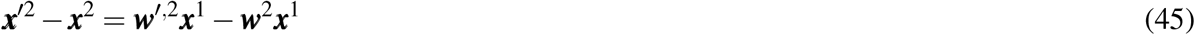

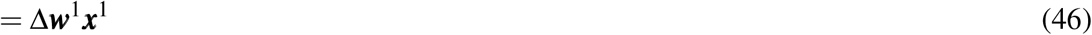

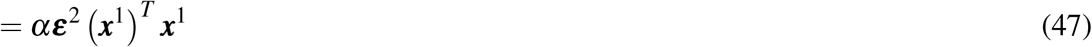

Here (***x***^1^)^*T*^ ***x***^1^ is a positive scalar (if at least one entry in ***x***^1^ is non-zero). Thus,

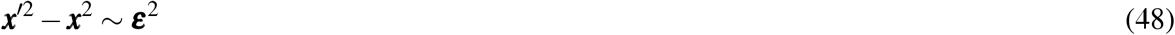

According to the definition of target alignment, which is the cosine similarity of the direction of the target (i.e., ***ε***^2^) and the direction of learning (i.e., ***x***^′2^ −***x***^2^), target alignment of this network is exactly one. This conclusion also applies to network with nonlinear activation function.

#### 2.4.2 Target alignment of target propagation (Extended Data Figs. 4a)

This subsection demonstrates that target alignment of target propagation is equal to 1. Such target alignment equal to 1 for target propagation is implied by Theorem 5 in the study of Meulemans et al.^58^. They show that if a network is linear and weights in each layer are invertible, then “parameter updates push the output activation along the negative gradient direction in the output space”^58^. Simulations in Extended Data Fig. 4a illustrate that the target alignment of target propagation is indeed equal to 1. For completeness we include in this paper a simple direct proof of this result (which we will also use in the next section).

For linear networks with invertible weights, the relationship between errors in adjacent layers in target propagation is:

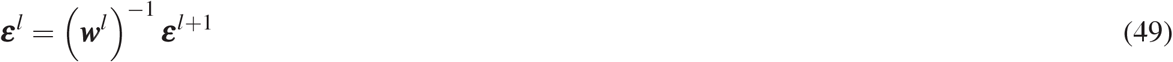

The activity of output neurons after the weight modification is:

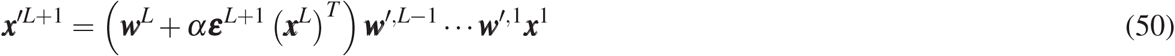

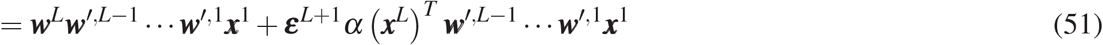

Term *α*(***x***^*L*^)^*T*^ ***w***^′,*L*−1^ ***w***^′,1^***x***^1^ is a scalar, so let us denote it by *c*_*L*_. Expanding ***w***^′,*L*−1^ and using Eq. (49), we obtain:

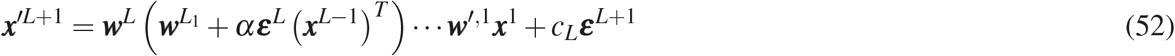

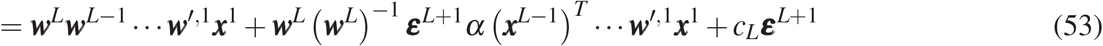

Note that ***w***^*L*^(***w***^*L*^)^−1^ is equal to the identity, so can be removed from the above equation, and *α*(***x***^*L*−1^)^*T*^ … ***w***^′,1^***x***^1^ is a scalar, so denote it by *c*_*L*−1_. Expanding all terms ***w***^′,*l*^ analogously as above, we eventually obtain:

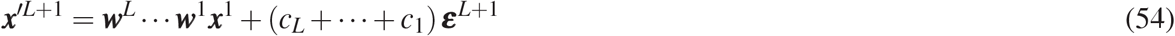

Since the output before weight update was ***w***^*L*^ … ***w***^1^***x***^1^, the change in the output is proportional to the direction towards target ***ε***^*L*+1^, hence the target alignment is equal to 1. Given the similarity between target propagation and predictive coding networks described in subsections 2.4.4 and 2.2, the predictive coding networks should also have target alignment relatively close to 1.

Since target propagation has a desirable property of perfect target alignment, one may ask if the brain can employ target propagation rather than prospective configuration as is main learning principle. However, energy-based networks have several advantages over target propagation both in terms of computational properties and relationship with experimental data. Since target propagation requires computation of multiple matrix inverses, it is numerically unstable, so for example in Extended Data Fig. 4a we only show the result for networks with up to 5 layers, because we were unable to perform target propagation in deeper networks due to numerical instabilities. Predictive coding networks offer a nice alternative which approximates target propagation, but is numerically stable. Furthermore, target propagation does not modify the activity of the neurons during relaxation, so it does not follow prospective configuration. Consequently, in the case of the network in Fig. 1 target propagation would not compensate the weight to olfactory output, because such compensation relies on updating the activity of the hidden neuron. Theory reviewed in this section implies that target propagation only produces target alignment equal to 1 if the weights are invertable, but this is not the case in the network in Fig. 1, so target propagation would not produce unity target alignment for this problem. Moreover, target propagation would not be able to reproduce the patterns of behaviour and neural activity in Figs. 5, 6 and 7, because reproducing these data relies on modifying activity of hidden neurons after feedback, and target propagation does not do it.

#### 2.4.3 Target alignment for orthogonal initialization (Extended Data Figs. 4c)

This subsection identifies one special conditions under which backpropagation produces target alignment of 1. Specifically, simulations in Extended Data Fig. 4c show that target alignment is equal to 1 for backpropagation in linear networks, when the weights are initialized to orthogonal values (***w***^*l*^)^*T*^ = ***w***^*l*^. This observation can be explained using results from the previous section: when weights are orthogonal, then (***w***^*l*^)^*T*^ = (***w***^*l*^)^−1^, hence the relationship between errors in adjacent layers is the same as for target propagation (Eq. (49)). Consequently, the same argument can be applied to backpropagation on linear networks with orthogonal initialization to show that it has target alignment equal to 1.

#### 2.4.4 Target alignment of predictive coding networks

The subsection addresses the question of why predictive coding networks have higher target alignment than backpropagation, using several findings in earlier sections. Specifically, to justify why predictive coding networks have high target alignment, we can combine 3 facts that we demonstrate in earlier sections, and summarize here:

1. Target alignment of target propagation is equal to 1. This is shown in Section 2.4.2.
2. When target pattern is provided to output neurons in predictive coding networks, during relaxation the neural activity in hidden layers converges to values related to local targets in target propagation. This is shown in Section 2.2.
3. Weight modification in predictive coding network reinforces the pattern of activity to which it converged during relaxation. In other words, predicting activity changes as a result of weight modification in the direction of the equilibrium reached during relaxation. This is shown in Section 2.3.

According to fact 3, learning in predictive coding networks reinforces the equilibrium activity, which, according to fact 2, is largely dependent on the local targets. Therefore, the changes in activity in hidden layers due to learning in predictive coding networks are similar to those in target propagation, and hence the changes in the output activity are also likely to be similar, and the two algorithms should also share a similarity in target alignment. According to fact 1, target propagation has target alignment of 1, so the predictive coding should also share a similar target alignment.

## Footnotes

1 Note that in realistic networks the weight matrices are not all square so an exact inverse (***w***^*l*^)^−1^ does not exist. Instead, we can compute approximations of the inverse using the Moore-Penrose pseudoinverse^133^ (***w***^*l*^)^†^, which is the least squares solution to the optimization problem argmin_***w***_ ***‖ I*** − ***w***^*l*^ ***w ‖***.

